# Variant effect predictions capture some aspects of deep mutational scanning experiments

**DOI:** 10.1101/859603

**Authors:** Jonas Reeb, Theresa Wirth, Burkhard Rost

## Abstract

Deep mutational scanning (DMS) studies exploit the mutational landscape of sequence variation by systematically and comprehensively assaying the effect of single amino acid variants (SAVs) for particular proteins. Different experimental protocols proxy effect through a diversity of measures. We evaluated three early prediction methods trained on traditional variant effect data (PolyPhen-2, SIFT, SNAP2) along with a regression method optimized on DMS data (Envision). On a common subset of 32,981 SAVs, all methods capture some aspects of variant effects, albeit not the same. Early effect prediction methods correlated slightly more with measurements and better classified binary states (effect or neutral), while Envision predicted better the precise degree of effect. Most surprising was that a simple approach predicting residues conserved in families (found and aligned by PSI-BLAST) in many cases outperformed other methods. All methods predicted beneficial effects (gain-of-function) significantly worse than deleterious (loss-of-function). For the few proteins with several DMS measurements, experiments agreed more with each other than predictions with experiments. Our findings highlight challenges and opportunities of DMS for improving variant effect predictions.

## Introduction

Recent human sequencing projects conclude that we all carry about 10,000 single amino acid variants (**<u>SAVs</u>**) with respect to the “reference genome” [1, 2]. Many of these SAVs are assumed to be neutral, while others might change protein function, contributing to complex phenotypes and causing diseases through a single change. Unfortunately, the gap between SAVs with and without experimental characterization continues to widen [3]: for only one in 10,000 of the known SAVs some experimental information is available [4, 5]. On top, many of those for which something is known are disease associations that may be wrong [6]. As long as the ability to interpret SAV effect does not improve, both on the level of the organism and the protein, the promise of precision medicine remains unmet [7–10].

Through the increased efficiency of sequencing, a procedure formerly used primarily *in silico* [11, 12] has become feasible for experiments, namely assessing the effect of all possible SAVs in a protein. In such deep mutational scanning (**<u>DMS</u>**) studies, a sequence library with all possible variants is created [13, 14] and exposed to a selection system. In the simplest case, the (logarithmic) difference between sequence frequencies with and without selection pressure yield an effect score for individual or combinations of variants [8, 15–17]. Variants with beneficial effect on protein function are discovered alongside the more commonly reported deleterious effect variants. DMS therefore aims at measuring the landscape of functional fitness for select proteins [18].

DMS has also been applied to screen proteins for improved drug binding, antibody affinity, using non-native chemical stresses, or non-proteinogenic amino acids, and on synthetic proteins [19–26]. DMS share objectives with directed evolution experiments which assay a wide range of mutations to engineer proteins with specific purposes. In fact, protein engineering has benefited from DMS studies [14].

One major challenge for DMS is the development of an assay to measure effect. Evaluating proteins with multiple functions requires multiple assays [8]. For instance, the effect of variants on Ubiquitin-60S ribosomal protein L40 has been assessed through their direct impact on yeast growth, as well as, through the impaired activation by the E1 enzyme [27, 28]. BRCA1 has been assayed through E3 ubiquitin ligase activity, through BARD1 binding and transcript abundance [29, 30]. Even for the same assay, specific experimental conditions might influence measurements [31]. Recently, a protocol for measuring protein abundance has been suggested as a proxy for function and applicable to many proteins [32]. The conclusions from DMS studies are limited by the validity of their functional assays; inferences of more complex effect relationships such as disease risk or clinically actionable pathogenicity often remain too speculative [8, 17].

Long before experimental DMS, prediction methods had addressed the same task *in silico* [33, 34] [35] [36–41]. These methods were developed on very limited datasets of effect and neutral variants. Many methods focused on disease-causing SAVs, e.g. from OMIM [42]. Others used databases such as the heterogeneous dbSNP [43] or PMD [44] which catalogue variants according to their direct effect upon protein function or structure. CADD solved the problem of limited data sets and some bias in databases by considering all mutations that have become fixed in the human population as neutral and a simulated set of all other variants as having an effect [35]. The training data set determines the type of effect methods can learn. Consequently, methods differ and work only on the type of SAV used for development. Given the limitations in today’s data, all methods are optimized on relatively small, unrepresentative subsets: fewer than 85,000 of all possible 217 million human SAVs have some experimental annotations [45, 46]. Methods agree much more with one another for the SAVs with known annotation than for SAVs without annotations [47].

DMS data sets constitute a uniquely valuable resource for the evaluation of current SAV effect prediction methods [17, 48], not the least, because most have not used those data. The Fowler lab has, recently, published an excellent analysis of prediction methods on DMS datasets and developed a new regression-based prediction method, Envision, trained only on DMS data [49]. Here, we focus on the analysis of a larger set of DMS studies and present trends in their correlation with SAV effects predicted by four variant effect prediction methods.

## Results and Discussion

### DMS experiments focus on subsets of all possible variants

The base dataset for our analyses of DMS studies consisted of 22 functional effect measures from 18 proteins, nine of those from human (Supplementary Online Material (SOM), Fig. S1a, Table S1) [29, 30, 32, 50–65]. The number of variants per study differed from hundreds to 10,000. In total the set contained 68,447 variants; 2,358 (3%) of these were synonymous, the other 97% constituted SAVs.

DMS still does not imply that all possible variants are tested [66]. On the contrary, only ten of the 22 sets (45%) scored some variants for at least 98% of the residues (Table S1). Four DMS studies provided functional scores for over 90% of all possible 19 non-native SAVs. On average, 66% of the residues in the 22 DMS sets were found to have SAVs with both deleterious and beneficial effects (Table S2). The majority of SAVs were beneficial for 3 of 22 studies (14%), namely for C-C chemokine receptor type 5 (CCR5), Mitogen-activated protein kinase 1 (MAPK1) and Peroxisome proliferator-activated receptor γ (PPARG). For the other 19 studies deleterious outnumbered beneficial SAVs by factors of 1.5-22.5 (Fig. S1b). Due the asymmetry in numbers and experimental fidelity, deleterious and beneficial effect SAVs were analyzed separately.

### Higher correlation for binary classification methods

SetCommon comprised 32,981 effect SAVs (17,781 deleterious) with predictions from each method (Table 1). All prediction methods reached a positive correlation on the set of deleterious SAVs (Spearman ρ≥0.1, Fig. 1a-d, Table 2). Highest was SNAP2 (ρ=0.41), followed by PolyPhen-2, SIFT and Envision [37–39, 49]. 95% confidence intervals (<u>CIs</u>) never overlapped, and these differences were statistically significant (Table S3).

**Table 1:** Number of SAVs in aggregated datasets ^Δ^.

|  | Number of SAVs |  |  |  |
| --- | --- | --- | --- | --- |
|  | Total | Neutral | Deleterious | Beneficial |
| SetAll | 66089 | 818 * | 45382 | 19889 |
| SetCommon | 32981 | 0 | 17781 | 15200 |
| SetCommonSyn90 | 15621 | 8926 | 4545 | 2150 |
| SetCommonSyn95 | 15621 | 10587 | 3209 | 1825 |
| SetCommonSyn99 | 15621 | 13506 | 1548 | 567 |

<sup>Δ</sup> SetAll depicts the total number of SAVs collected, while SetCommon contains only SAVs with predictions from every analyzed method. SetCommonSyn contains all SAVs with predictions where a thresholding scheme could be applied to yield classification of SAVs into neutral and effect (see Methods). The number of SAVs in every single DMS experiment are depicted in Fig. S1 and Table S1.
\* The ccdB set classifies variant effect in categories and contains 818 non-synonymous variants which fall in the same category as the wt. Hence these SAVs could be considered neutral.

**Table 2:** Performance measures for five prediction methods on *SetCommon* ^Δ^.

|  | deleterious SAVs (n = 17781) |  | beneficial SAVs (n = 15200) |  |
| --- | --- | --- | --- | --- |
| | $\rho$ | MSE | $\rho$ | MSE |
| PolyPhen-2 | 0.33 [0.32, 0.34] | 0.45 [0.45, 0.46] | -0.09 [-0.10, -0.07] | 0.4 [0.40, 0.41] |
| SIFT | 0.23 [0.22, 0.25] | 0.58 [0.57, 0.58] | -0.14 [-0.15, -0.12] | 0.64 [0.63, 0.64] |
| SNAP2 | 0.41 [0.40, 0.42] | 0.3 [0.30, 0.30] | 0.02 [0.01, 0.04] | 0.23 [0.23, 0.24] |
| Envision | 0.1 [0.08, 0.11] | 0.06 [0.06, 0.07] | -0.14 [-0.16, -0.13] | 0.05 [0.04, 0.05] |
| Conservation | 0.29 [0.27, 0.30] | 0.19 [0.19, 0.19] | -0.08 [-0.09, -0.06] | 0.19 [0.19, 0.20] |

<sup>Δ</sup> SetCommon denotes the set of SAVs with predictions from every method (see Methods). $\rho$ denotes Spearman $\rho$ (higher is better), MSE the mean squared error (lower is better, Methods, SOM\_Note4). Normal Dist. denotes a random prediction of scores normally distributed around the mean of the experimental values and is given as a baseline. Values in brackets are 95% confidence intervals.

**Figure 1:**
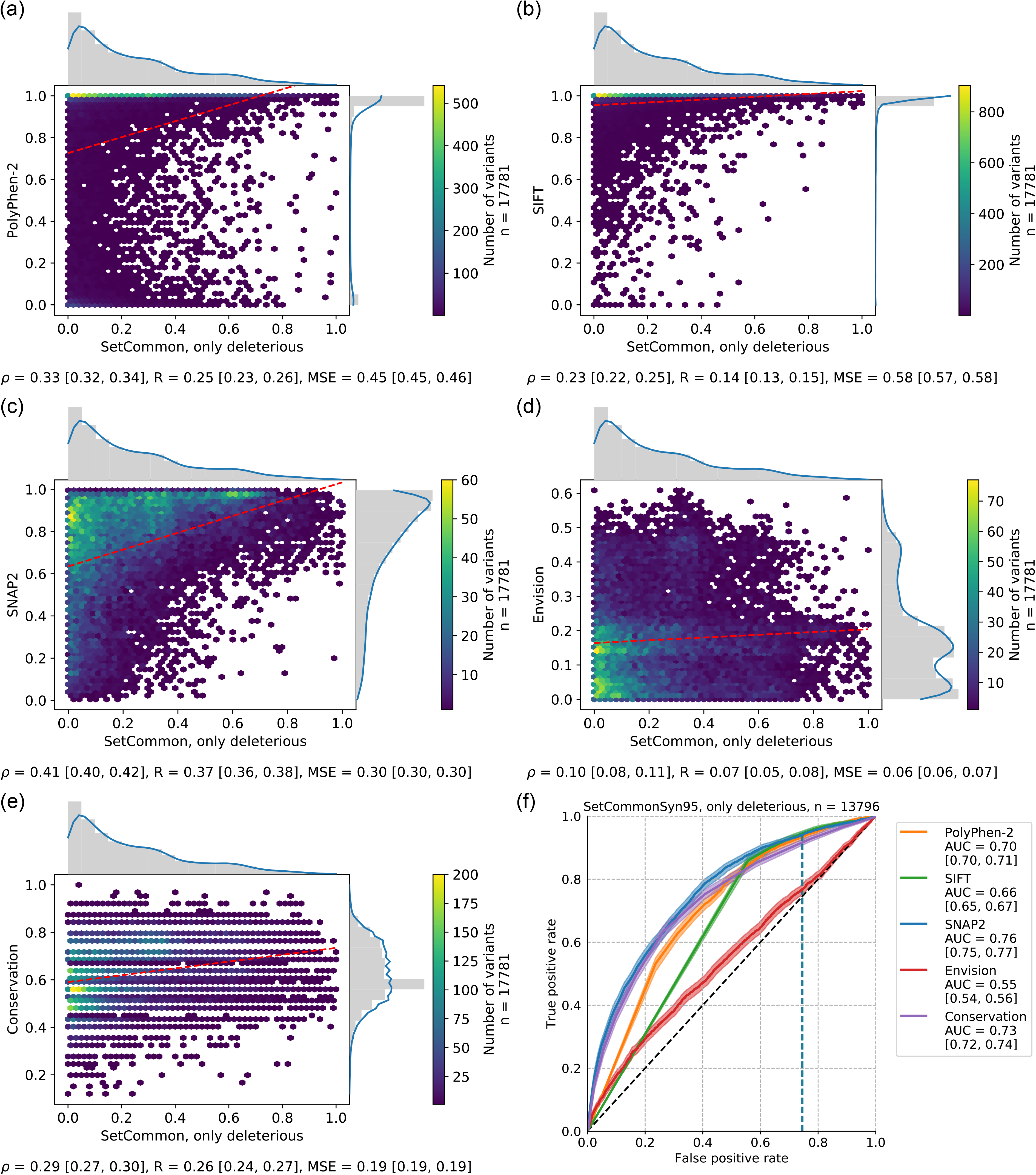
Agreements between DMS experiments and variant effect predictions. <u>Panels (a)-(e)</u> show the performance for five prediction methods (PolyPhen-2 [37], SIFT [39], SNAP2 [38], Envision [49], and naïve Conservation) through hexbin plots of 17,781 deleterious effect SAVs in SetCommon (see Methods, Processing). Values on axes range from 0 (neutral) to 1 (maximal effect). Dashed red lines give linear least-squared regressions. Marginals denote distributions of experimental and predicted scores with a kernel density estimation overlaid in blue. The footer denotes Spearman ρ, Pearson R and the mean squared error together with the respective 95% confidence intervals. The method scores are given on the y-axes and reveal the method: (a) PolyPhen-2, (b) SIFT, (c) SNAP2, (d) Envision – the only method trained on DMS data, (e) naΪve Conservation read off PSI-BLAST profiles. <u>Panel (f)</u> gives ROC curves for 13,796 deleterious effect SAVs which were classified into either neutral, defined by the middle 95% of the scores from synonymous variants, or effect (SetCommonSyn95, see Methods). Shaded areas around lines denote 95% confidence intervals. The legend denotes the AUC for each method along with the 95% confidence intervals. Horizontal dashed lines denote the default score threshold used by SNAP2 (blue) and SIFT (green).

Both SIFT and PolyPhen-2, by design, have almost binary prediction scores (either neutral or effect), and both score distributions are highly skewed toward effect. Both are optimized for capturing binary effects, not correlations, which was confirmed by two recent studies [48, 49]. The relatively high Spearman ρ values for these were, thus, somehow unexpected. On the other hand, SNAP2 and Envision scores appeared, overall, less binary (Figs. 1c-d, S3). For Envision this was expected, given that it was trained as a regression predictor on non-binary data. In contrast, SNAP2 was trained on binary classification data (effect or neutral). Nevertheless, predictions have been shown to correlate with effect strength [5, 67, 68]. SNAP2 distributions were skewed toward high effect, while Envision also succeeded in detecting SAVs with less pronounced effects (Fig. 1c-d).

### Envision approximates experimental values best

When evaluating methods by the numerical difference between experimental and predicted variant effect scores (mean squared error, MSE), Envision was best, followed at considerable distance by SNAP2, PolyPhen-2 and SIFT (Fig. 1a-d, Table 2). Envision was found to best distinguish low from high effect variants [49]. The training as regression method might have given it an advantage over classification methods. Envision’s good performance on this metric, partially originated from predicting no SAV with strong effect (the highest Envision score was 0.61, with a possible maximum of 1). Such a distribution resembled the overall experimental distribution that was skewed towards lower effect (Fig. 1d, gray distributions next to x- and y-axes). Predicting most SAVs somewhere in the middle constraints the maximally possible MSE. Indeed, shuffling the prediction scores achieved the same MSE (Fig. S2a). Furthermore, predicting a normal distribution around the mean of the experimental values performed slightly worse but still better than all other prediction methods (Fig. S2b, Table 2). On the other hand, MSE did not increase when limiting the evaluation to just the 25% strongest effect SAVs, although the benefit of predicting no strong effect SAVs should be reduced here (Table S4). When considering each DMS measurement separately, Envision also appeared to perform best except for transcriptional coactivator YAP1 (YAP1) which indeed had the most uniform distribution of effect scores (similar number of lowest, medium, and strongest effects observed; Fig. S3b, Table S5).

### Conservation captured deleterious effects well

Most surprising was the good performance of a naïve method exclusively using conservation (from PSI-BLAST profiles) to predict SAV effects. This naïve prediction appeared to correlate more with the DMS experiments than the specialized methods SIFT and Envision (Fig. 1e). Furthermore, the MSE was lower for Conservation than for all specialists except Envision (Fig. 1, Table 2). Disease causing SAVs from OMIM typically affect the most conserved residues [47], and it has been argued that machine-learning based predictions of variant effects are of limited applicability because they often largely capture conservation [17, 69–72]. Furthermore, non-specialized methods such as analyses of conservation patterns also capture some aspects of variant effect on function [73]. To some extent, our findings confirm this to be valid for DMS experiments, as well. Nonetheless, the distributions of effect predicted by Conservation and observed by DMS differed substantially (Fig. 1e, gray distributions).

### High effect SAVs from DMS not generally predicted better

The DMS sets for immunoglobulin gamma-1 heavy chain (IgG1) and modification methylase HaeIII (haeIIIM) were most skewed toward effect (Fig. S3o,v). SNAP2, PolyPhen-2 and SIFT had lower than average MSE on these two sets and also better than average correlation for haeIIIM (Tables S5, S6). Bias in the data used for developing those methods might explain this observation [5, 70, 71, 74]. However, the same trend was observed for the conservation-based prediction suggesting that for those two DMS experiments evolutionary information might capture variant effects best.

To investigate whether methods predict the strongest effect SAVs better, we analyzed subsets with the 75%, 50%, or 25% strongest effect SAVs (strongest deleterious: SetDelStrongest75∣50∣25, beneficial: SetBenStrongest75∣50∣25). MSE was higher for stronger deleterious effect SAVs for PolyPhen-2, SNAP2 and Conservation, much less so for SIFT, and not for Envision (Table S4). Therefore, the accuracy in predicting exactly how much effect a SAV has was not increased for strong effect SAVs. Furthermore, the Spearman ρ correlation decreased for all methods for SetDelStrongest75, 50 and 25. This again suggested most methods to succeed only at distinguishing lower from high effect SAVs, on average. These results were surprising, because, for instance, SNAP2 prediction scores correlated with effect strengths, i.e. stronger effect SAVs tend to be predicted better [47, 68, 75]. This correlation might originate from so-called toggle positions, i.e. residues for which SAVs have either a strong effect or none [71, 76].

### Beneficial effects difficult to predict

Unlike for deleterious SAVs, no method correlated, on average, with beneficial effect SAVs (−0.14≤ρ≤0.02, Tables 2, S7, Fig. S4) and performance was only slightly increased when focusing on high effect subsets (Table S4). This suggested beneficial effect SAVs to have distinctly different signatures from their deleterious counterparts, too distinct for current methods to capture. The conservation-based prediction also decreased substantially from Spearman ρ of 0.29 for deleterious to −0.08 for beneficial SAVs (Table 2, Fig. S4e). Furthermore, SNAP2 and PolyPhen-2 show a clear shift of prediction scores towards lower effect compared to predictions of deleterious SAVs (cf. Fig 1a,c and Fig. S4a,c, gray distributions). This was particularly pronounced for levoglucosan kinase (LGK), haeIIIM, β-glucosidase (bgl3) and β-lactamase TEM (bla) which suggests that beneficial effect SAVs were often mistaken as having no or low effect (Fig. S5j, n, o, t). Possibly, because most SAVs in the development sets of these methods were deleterious.

Unlike Spearman ρ, the MSE for beneficial effect SAVs was similar to that for deleterious variants. Envision again performed best by far (MSE=0.05, Tables 2, S3), also when treating each set separately (Table S8, Fig. S5). Envision was trained with 25% beneficial effect variants (SOM_Note1). However, the decreased correlation performance (ρ = −0.14 versus 0.1), highlighted that just including these variants in training did not suffice to learn their signatures.

### Experimental agreement sets the benchmark for prediction methods

The above comparisons of experimental and predicted SAV effects raise the question of what agreement can realistically be obtained. One proxy for an answer is the comparison of different DMS studies conducted on the same protein.

For deleterious SAVs, the correlation between experiments was high (Spearman ρ = 0.65 vs. 0.41 for the top predictions from SNAP2, Table S9, Fig. S6). The lowest correlation was that between two measurements on breast cancer type 1 susceptibility protein, BRCA1 and BRCA1_2015_E3 (ρ=0.21, Fig. S6b). This was the only set for which prediction methods outperformed experimental agreement. Rather than experimental noise, the low correlation might also originate from different experimental setups employed for multi-functional proteins such as BRCA1. The strong correlation (ρ=0.93) between two experiments that measured the same condition for bla (beta-lactamase TEM precursor; bla and bla_2014, Fig. S6h) provided a single case in strong support of such an explanation. Possibly, prediction methods only detect one of multiple aspects of effect, or the effects measured by all functional assays. The MSE between experiments was not lower than that between experiments and Envision (mean MSE=0.07, Table S9, Table S5).

In contrast to deleterious SAVs, experiments did not correlate for beneficial effect (mean ρ=0.03) although the MSE remained low (mean MSE=0.05, Table S9, Fig. S7). The major issue for this comparison were the small numbers (572 SAVs from five assays). Maybe experimental assays are biased towards measuring deleterious effects. This also put the poor predictions of beneficial SAVs effects into perspective.

### Classification into neutral and effect provides similar picture as regression analyses

Scores from binary classification methods (neutral or effect) such as SNAP2, PolyPhen-2 and SIFT primarily measure the method’s confidence in the predicted class. Classification performance is often assessed through receiver operating characteristic (ROC) curves avoiding the choice of a particular single threshold in making the decision between neutral and effect. Toward this end, we assigned classes to the SAVs in SetCommon through normalization by experimental measurements of synonymous variants [60] (Methods).

On the 3,209 deleterious effect SAVs of SetCommonSyn95 (10,587 neutral, Table 1, Fig. S8), SNAP2 achieved the highest area under the curve (AUC, 0.76, 95% CI [0.75, 0.77]). It was the only method better (with a statistically significant difference) than the naïve conservation-based method (0.73 [0.72, 0.74], Fig. 1f, Table S10). Precision-recall curves also highlight the smooth transition of SNAP2 opposed to the naïve approach of Conservation although both methods share similar peak performance (Fig S9). For this task, for which it had not been developed, Envision performed better than random, but clearly worse than the specialized classification methods (AUC = 0.55 [0.54, 0.56]). On the BRCA1 set, Envision performed on par with the best methods (Fig. S10a). The reason remained unclear, but for this set correlation was also highest (Table S6). Overall, the simple assumption that classification methods perform better on the task they were trained for was not supported by our results: The four proteins considered here (BRCA1, PPARG, PTEN and TPMT), also correlated above average for SNAP2, PolyPhen-2 and SIFT (Table S6). Using less or more stringent definitions for classifying experimental SAVs (SetCommonSyn90 and SetCommonSyn99, see Methods, Fig. S11a-b) did not change these results.

Beneficial SAVs were also difficult to classify: PolyPhen-2 and SNAP2 performed best with AUC=0.62, followed by SIFT, while Envision predictions were not better than random (Fig. S4f, Fig. S11c-d, Table S10). Naïve Conservation also performed significantly worse at a level of random predictions.

## Conclusions

Deep mutational scanning (DMS) studies set out to explore the relation between protein sequence and function by systematically probing sequence variants. We collected 22 DMS experiments from the literature and focused on single amino acid variants (SAVs). Most studies probe only a small subset of all possible variants (for a protein with N residues, there are 19*N non-native sequence variants). We set out to evaluate whether DMS studies agree with each other and with predictions. One result was that two experiments probing the same protein tend to agree more with each other than either of those agrees with predictions. Nonetheless, different experiments use different proxies for function and disagree substantially, and all experiments differ significantly from all prediction methods. On top of the problem that different experiments probe different proxies for function, we failed to identify a single measure for how similar experiments and predictions are (see SOM_Note2). Lacking a convincing solution, we discussed different aspects and scoring schemes, showing as much as possible the raw data.

We analyzed five variant effect prediction methods: *Envision*, *naïve Conservation* (essentially using PSI-Blast conservation to predict effect/neutral), *PolyPhen-2*, *SIFT* and *SNAP2*. Looking at the distributions for experimental and predicted SAV effects (Fig. 1), we noted that for deleterious SAVs all methods reached positive Spearman ρ correlations with the DMS experiments. Traditional classification methods actually achieved highest correlations with effect strength, although much of their performance can be achieved by evolutionary information, i.e. read off PSI-BLAST profiles. On the other hand, the lowest mean squared error (MSE) was achieved by Envision, the only tool currently available which has been trained and optimized for regression on DMS data. In fact, the Envision MSE was as low as that between experiments.

Sorting performance by Spearman ρ and by MSE, put different methods on top showing that methods captured diverse aspects measured by the DMS studies. All methods performed better on SAVs with deleterious than with beneficial effect. This highlighted the common negligence and lack of data for beneficial effect SAVs. However, experiments also agreed much more with each other for deleterious than for beneficial effect SAVs which demonstrates the challenge in experimentally describing beneficial SAVs comprehensively and correctly.

Although binary classification methods, surprisingly, captured aspects of non-binary measurements, they performed much better for the binary classification task (projecting DMS results onto neutral vs. effect; Fig. 1f). Notably, the naïve Conservation prediction captured effect better than some more advanced tools.

The challenge for the next generation of prediction methods will be to learn from the diversity of DMS. To give just one example: OMIM, a popular source of training data, contained ~11,000 SAVs referenced in dbSNP (02/2019). This is a magnitude matched by a single large DMS experiment. The generality of a singular SAV in both sets might not be comparable, yet DMS opens up variant effect prediction to new methodologies. For instance, increasingly popular deep learning approaches which typically require training sets magnitudes larger than what has traditionally been used for variant effect [77, 78]. Furthermore, the increase in amount and diversity of training data could help to improve the applicability of methods by being less reliant on conservation. It might also allow methods to account for different SAV types such as toggle and rheostat positions [71]. Finally, DMS studies contain many beneficial effect SAVs that have, so far, been underrepresented in databases. Clearly, DMS should play a major role for the future of variant effect prediction methods and give rise to new methods which are better estimators of SAV effects both on the level of protein function as well as organism-level disease-causing variants. Only then will we come closer to attaining the goals of precision medicine.

## Methods

### Dataset collection

We retrieved all DMS datasets available by June 2019 that report over 100 SAVs available from the literature. Functional effect scores were taken directly from the supplemental material published or requested from the authors (Table S11). The data were formatted in a variety of formats including Excel, and tab- or comma-separated files. Scores were manually mapped either to the UniProtKB identifier given in the publication or to its closest BLAST match (Table S12) [45, 79]. Six of the 22 experiments contained up to five substitutions (pairwise sequence identity ≥98%); those were maintained for prediction. We refer to the combined data as *SetAll* (66,089 SAVs) supplemented by *SetCommon* with 32,981 SAVs from ten of the 22 experiments for which we had a prediction from every method tested (Table 1). During completion of this manuscript, MaveDB, a centralized resource of multiplexed assays of variant effect has been published [8, 80]. MaveDB identifiers exist for ten of our 22 datasets (November 2019, Table S11).

SetAll contained several proteins with multiple independent experimental measurements. Inclusion of additional sets analyzed previously [49], yielded a total of three measurements for Hsp82 and BRCA1 and two for both beta-lactamase and ubiquitin (Table S1) [27, 28, 81]. Performance measures were calculated only on SAVs and not between DMS measurements from the same publication. For analysis of beneficial effect SAVs, all studies on Hsp82 had to be excluded since the sets contain only three of those SAVs each.

Three subsets of SetAll were created containing the SAVs with the 75%, 50%, or 25% highest deleterious or beneficial effect for each DMS experiment (*SetBenStrongest75∣50∣25* and *SetDelStrongest75∣50∣25*); the YAP1 data set was removed for SetBenStrongest25, because it contained fewer than 50 SAVs after filtering.

### Processing functional effect scores

Several DMS studies provide multiple effect scores for the same protein. We generally chose the scores with the highest evolutionary pressure (Table S13). In the following processing, effect scores were left as provided by the authors as much as possible but adjusted such that the wild-type score for each measurement (Table S14) became 0, and larger values denoted more effect up to the maximum at 1 (SOM_Note3). Beneficial and deleterious effects had to be analyzed separately because experimental assays were not symmetrical and further normalization might over- or underrepresent effects. The resulting score distributions differed significantly between experiments (e.g. in contrast to the more homogeneous subset used previously [49]).

We also created sets with binary classifications (effect vs. neutral) from all DMS studies with synonymous variants: The middle 95% of effect score values from synonymous variants was used to define which SAVs were considered neutral. All SAVs outside this range were considered as effect. We applied the same procedure using 90% or 99% of synonymous variants’ values and refer to the thresholding schemes as *syn90*, *syn95*, and *syn99*. Applying these schemes to the four experiments in SetCommon which have synonymous variants (BRCA1, PTEN, TPMT, PPARG) yields *SetCommonSyn90∣95∣99*. Again, deleterious and beneficial effect SAVs were analyzed separately.

### Performance measures

Experiments and predictions were compared through three measures (SOM_Note4, SOM_Note2): (1) mean squared error (**<u>MSE</u>**) calculated with the scikit-learn metrics module [82]; (2) Pearson R (pearsonr) and (3) Spearman ρ (spearmanr) both calculated with the SciPy stats module [83]. For convenience linear least-squares regression lines (linregress) were added to the correlation plots. Pearson R was added for ease of comparison to others but not discussed as it is not robust and most datasets violated both its validity assumptions (normal distribution & absence of significant outliers [84]). We further found no evidence to supplement MSE by a measure more robust to outliers (SOM_Note2). 95% confidence intervals (**<u>CIs</u>**) for R, ρ and MSE were estimated using a percentile bootstrap with 1000 random samples with replacement.

The performance of binary predictions (effect vs. neutral) was measured through receiver operating characteristic (**<u>ROC</u>**) curves and the area under those curves (**<u>AUC</u>**) calculated through the pROC package in R, which was also used to calculate 95% confidence intervals of ROC (`ci.se`) and AUC (`ci.auc`) [85, 86]. Additionally, precision-recall curves were created using scikit-learn (`precision-recall-curve`).

### Prediction methods

The sequences determined during dataset collection were used as input to a set of commonly used variant effect prediction methods. Each method was run to predict the effect of all 19 non-native amino acids at every position in the protein. *SNAP2* [38] was run locally using default parameters on UniProtKB (Release 2018_09). *SIFT* version 6.2.1 [39] was run locally (UniProtKB/TrEMBL Release 2018_10). *PolyPhen-2* [37] predictions were retrieved from the webserver in batch mode (http://genetics.bwh.harvard.edu/pph2/bgi.shtml) with classification model humdiv on genome assembly GRCh37/hg19 and default parameters. Predictions failed for all relevant residues of the three DMS studies on Hsp82. *Envision* [49] predictions were retrieved online (https://envision.gs.washington.edu/shiny/envision_new/) which requires UniProtKB identifiers as input. Therefore, Envision predictions could be analyzed only for ten proteins (Table S15). While SNAP2 and SIFT predicted all SAVs, PolyPhen-2 and Envision failed for some residues, shrinking the size of the data sets. We always report performance on the largest common subset of SAVs per dataset.

As a baseline, predictions were also created by running PSI-BLAST with three iterations on UniProtKB (Release 2018_09). Scores from the resulting profile (position-specific scoring matrix) had their signs flipped and were then directly used as a measure of effect, i.e. less frequent substitutions have a higher effect than conserved ones. We refer to this method as naïve *Conservation*.

For SIFT, scores were reversed such that higher values implied higher effect. The same was done for Envision predictions of deleterious effect. Envision predictions of beneficial effect were treated separately and mapped to the range of [0,0.2]. This yielded the same performance than scaling between [0,1] or no scaling (SOM_Note5). Finally, prediction scores of all methods were adjusted to lie between 0 (no effect) and 1 (highest effect) using the theoretical maximum and minimum prediction value of every method.

## Abbreviations used

DMS: deep mutational scanning
SAV: single amino acid variant
ROC: receiver operating characteristic
AUC: area under the ROC curve
CI: confidence interval
MSE: Mean squared error

## Acknowledgements

The authors wish to thank all groups that work on DMS and readily provided their data, either as part of their manuscript or swiftly upon personal contact. Thanks also to Michael Bernhofer and Maria Schelling for helpful discussions, Inga Weise for administrative support and Tim Karl for help with hard- and software. This work was supported by the Deutsche Forschungsgemeinschaft (DFG) – project number 5091000. Further funding was provided by the Bavarian Ministry for Education through funding to the TUM paying for the positions of the authors.

