## Supplemental Material for "Variant effect predictions capture some aspects of deep mutational scanning experiments"

### Supporting online material
### for:
### Variant effect predictions capture some aspects of deep mutational scanning experiments

**Jonas Reeb, Theresa Wirth & Burkhard Rost**

### Material

**Figure S1: Size of 22 DMS measurements comprising SetAll.**

(a) The number of (non-) synonymous variants per DMS experiment is shown with the short identifier that is used to reference the respective measurement in parentheses. (b) Number of SAVs with deleterious or beneficial effect (see Methods). Variants with effect equal to the wild-type are considered neither deleterious nor beneficial. Sets with <250 beneficial variants have those numbers written out. Note that ccdB values are categorical and only contain deleterious effect variants.

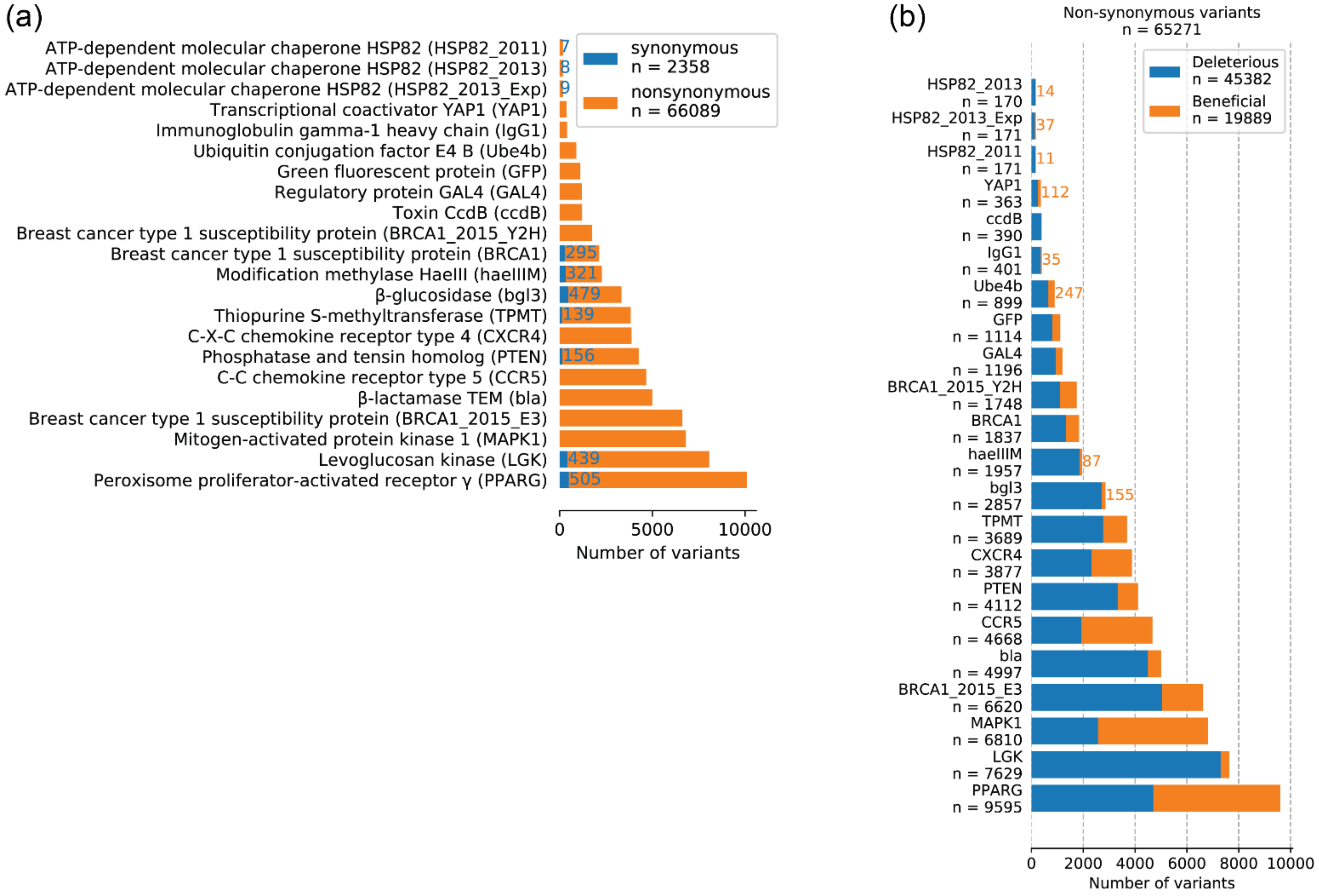

**Figure S2: Random predictions on deleterious SAVs of SetCommon.**

To assess the effect of Envisions score distribution on the resulting MSE, (a) the prediction scores for the 17,781 SAVs were randomly shuffled. Due to randomized nature of this setup MSEs slightly differed for various runs but were always close to the original MSE of 0.06 and always lower than any other prediction method's MSE. (b) Randomly generated prediction scores with a normal distribution around the mean of the experimental values. The resulting MSE of 0.13 is higher than that of Envision, but lower than the next best method, Conservation (MSE = 0.19).

(a) (b)

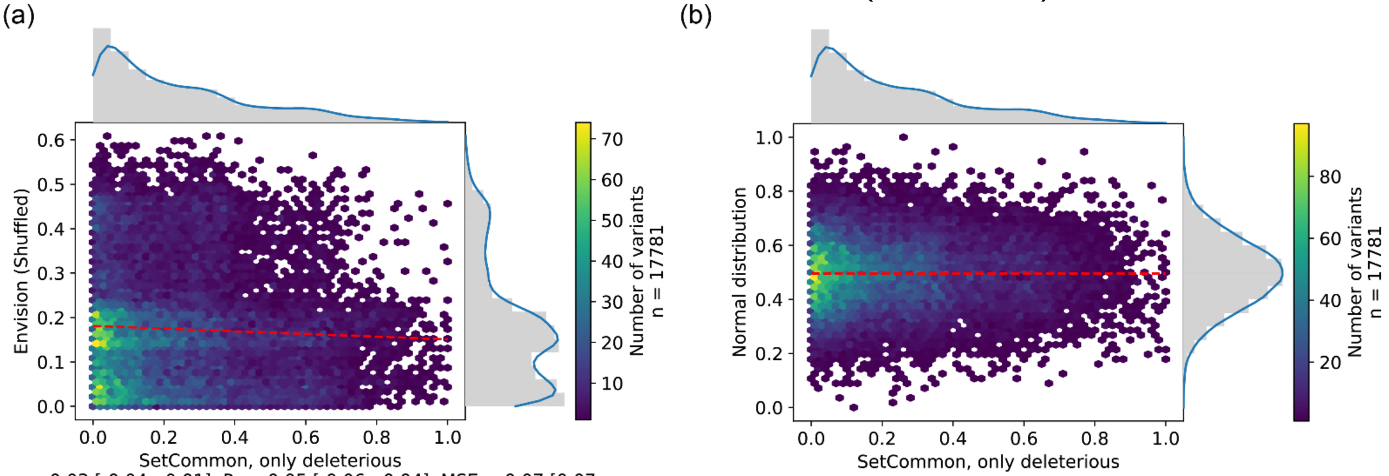

$\rho$ = -0.03 [-0.04, -0.01], R = -0.05 [-0.06, -0.04], MSE = 0.07 [0.07, 0.07]

$\rho$ = 0.01 [-0.01, 0.02], R = 0.00 [-0.01, 0.02], MSE = 0.12 [0.12, 0.13]

#### Figure S3: Agreements between prediction methods and deleterious effect scores from experiments.

Panels (a)-(v) show the performance for five prediction methods through hexbin plots for all 22 DMS datasets in SetAll (see Fig. S1b, Table S1, Methods). For every dataset only the largest common subset of deleterious effect SAVs for which a prediction was available from every method is analyzed. Missing methods did not perform any predictions at all. Values on axes range from 0 (neutral) to 1 (maximal effect). Dashed red lines give linear least-squared regressions. Marginals denote distributions of experimental and predicted scores with a kernel density estimation overlaid in blue. The footer denotes Spearman ρ, Pearson R and the mean squared error together with the respective 95% confidence intervals. The method scores are given on the y-axes and reveal the methods: PolyPhen-2, SIFT, SNAP2, Envision (the only method trained on DMS data), naïve conservation read off PSI-BLAST profiles.

S3a

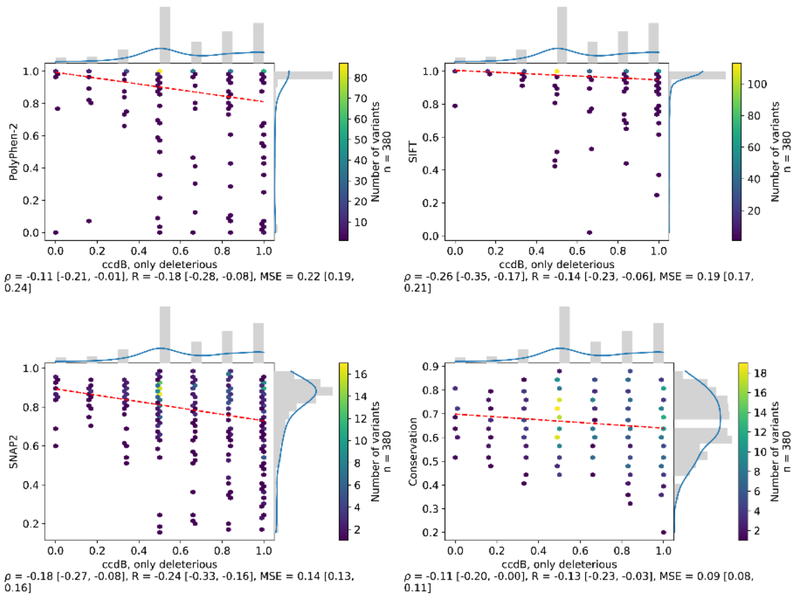
PolyPhen-2
ccdB, only deleterious
Number of variants
n = 380
ρ = -0.11 [-0.21, -0.01], R = -0.18 [-0.28, -0.08], MSE = 0.22 [0.19, 0.24]
SIFT
ccdB, only deleterious
Number of variants
n = 380
ρ = -0.26 [-0.35, -0.17], R = -0.14 [-0.23, -0.06], MSE = 0.19 [0.17, 0.21]
SNAP2
ccdB, only deleterious
Number of variants
n = 380
ρ = -0.18 [-0.27, -0.08], R = -0.24 [-0.33, -0.16], MSE = 0.14 [0.13, 0.16]
Conservation
ccdB, only deleterious
Number of variants
n = 380
ρ = -0.11 [-0.20, -0.00], R = -0.13 [-0.23, -0.03], MSE = 0.09 [0.08, 0.11]

S3b

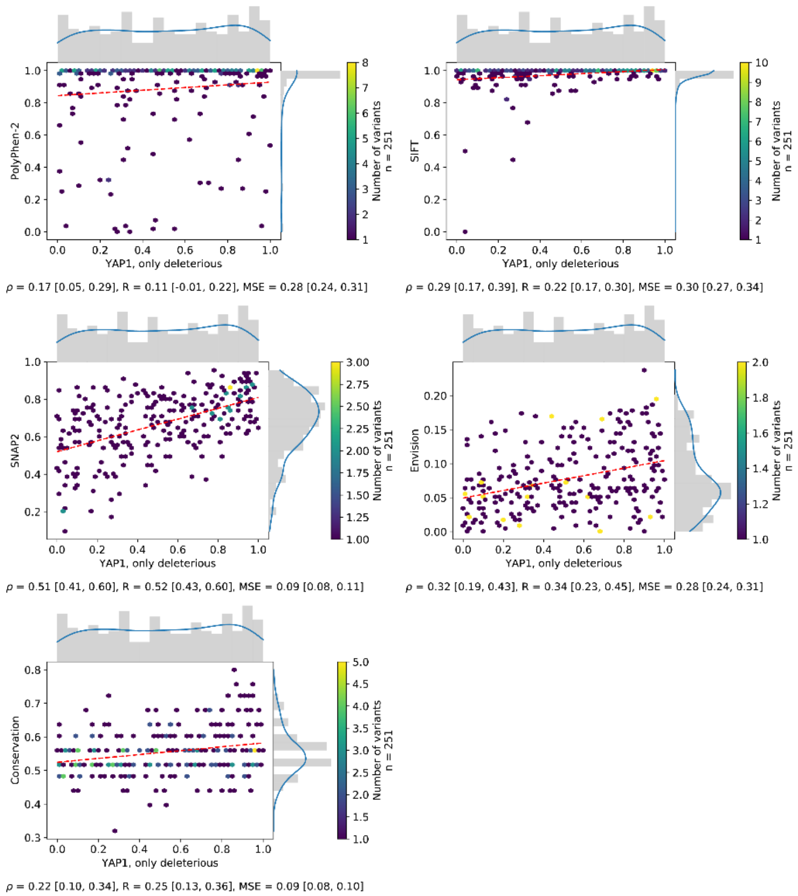
PolyPhen-2
YAP1, only deleterious
Number of variants
n = 251
ρ = 0.17 [0.05, 0.29], R = 0.11 [-0.01, 0.22], MSE = 0.28 [0.24, 0.31]
SIFT
YAP1, only deleterious
Number of variants
n = 251
ρ = 0.29 [0.17, 0.39], R = 0.22 [0.17, 0.30], MSE = 0.30 [0.27, 0.34]
SNAP2
YAP1, only deleterious
Number of variants
n = 251
ρ = 0.51 [0.41, 0.60], R = 0.52 [0.43, 0.60], MSE = 0.09 [0.08, 0.11]
Envision
YAP1, only deleterious
Number of variants
n = 251
ρ = 0.32 [0.19, 0.43], R = 0.34 [0.23, 0.45], MSE = 0.28 [0.24, 0.31]
Conservation
YAP1, only deleterious
Number of variants
n = 251
ρ = 0.22 [0.10, 0.34], R = 0.25 [0.13, 0.36], MSE = 0.09 [0.08, 0.10]

## S3c

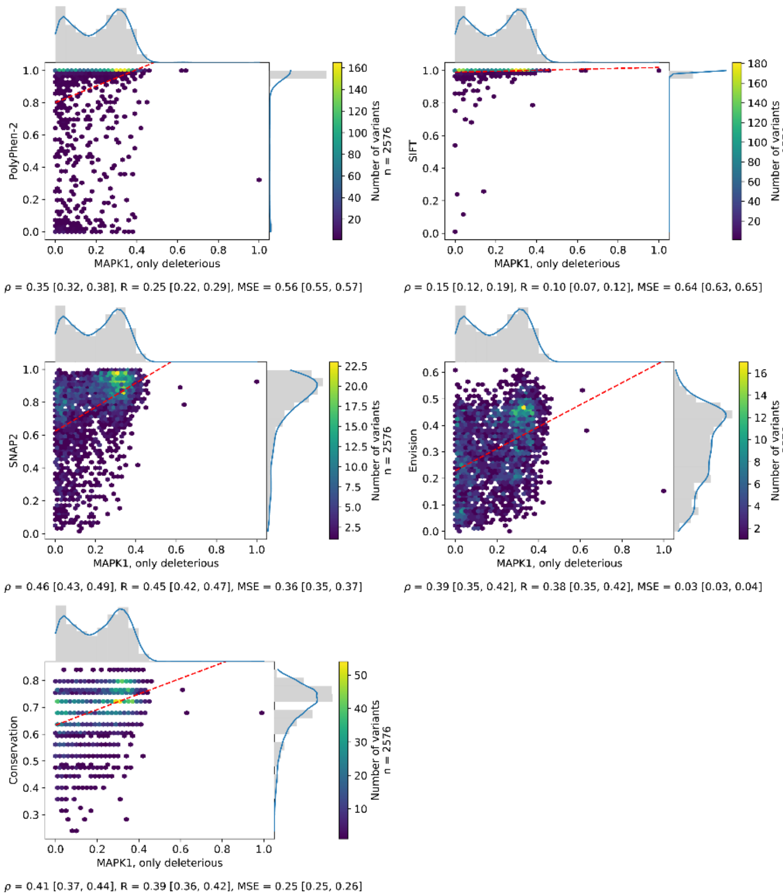
PolyPhen-2
SIFT
SNAP2
Envision
Conservation
MAPK1, only deleterious
Number of variants
n = 2576
ρ = 0.35 [0.32, 0.38], R = 0.25 [0.22, 0.29], MSE = 0.56 [0.55, 0.57]
ρ = 0.15 [0.12, 0.19], R = 0.10 [0.07, 0.12], MSE = 0.64 [0.63, 0.65]
ρ = 0.46 [0.43, 0.49], R = 0.45 [0.42, 0.47], MSE = 0.36 [0.35, 0.37]
ρ = 0.39 [0.35, 0.42], R = 0.38 [0.35, 0.42], MSE = 0.03 [0.03, 0.04]
ρ = 0.41 [0.37, 0.44], R = 0.39 [0.36, 0.42], MSE = 0.25 [0.25, 0.26]

## S3d

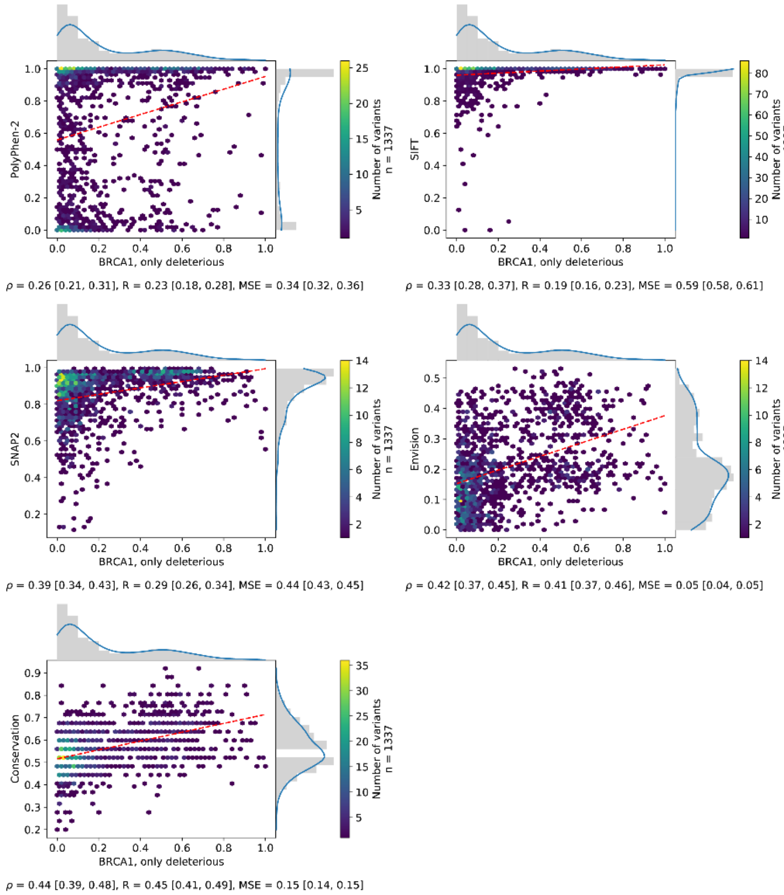
PolyPhen-2
SIFT
SNAP2
Envision
Conservation
BRCA1, only deleterious
Number of variants
n = 1337
ρ = 0.26 [0.21, 0.31], R = 0.23 [0.18, 0.28], MSE = 0.34 [0.32, 0.36]
ρ = 0.33 [0.28, 0.37], R = 0.19 [0.16, 0.23], MSE = 0.59 [0.58, 0.61]
ρ = 0.39 [0.34, 0.43], R = 0.29 [0.26, 0.34], MSE = 0.44 [0.43, 0.45]
ρ = 0.42 [0.37, 0.45], R = 0.41 [0.37, 0.46], MSE = 0.05 [0.04, 0.05]
ρ = 0.44 [0.39, 0.48], R = 0.45 [0.41, 0.49], MSE = 0.15 [0.14, 0.15]

## S3e

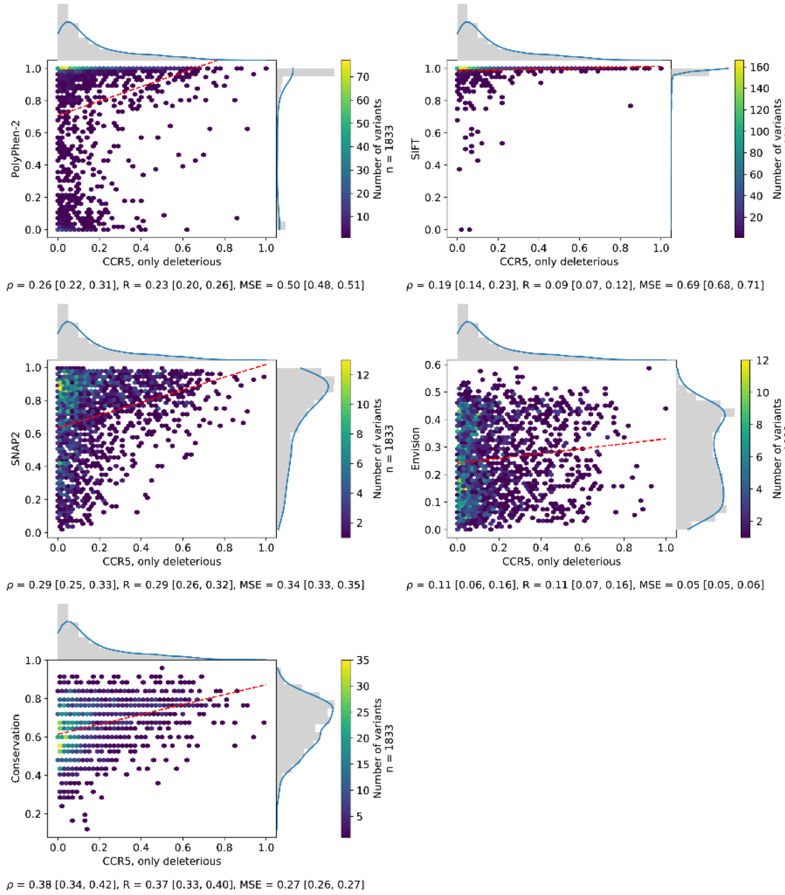
PolyPhen-2
CCR5, only deleterious
Number of variants
n = 1833
ρ = 0.26 [0.22, 0.31], R = 0.23 [0.20, 0.26], MSE = 0.50 [0.48, 0.51]
SIFT
CCR5, only deleterious
Number of variants
n = 1833
ρ = 0.19 [0.14, 0.23], R = 0.09 [0.07, 0.12], MSE = 0.69 [0.68, 0.71]
SNAP2
CCR5, only deleterious
Number of variants
n = 1833
ρ = 0.29 [0.25, 0.33], R = 0.29 [0.26, 0.32], MSE = 0.34 [0.33, 0.35]
Envision
CCR5, only deleterious
Number of variants
n = 1833
ρ = 0.11 [0.06, 0.16], R = 0.11 [0.07, 0.16], MSE = 0.05 [0.05, 0.06]
Conservation
CCR5, only deleterious
Number of variants
n = 1833
ρ = 0.38 [0.34, 0.42], R = 0.37 [0.33, 0.40], MSE = 0.27 [0.26, 0.27]

## S3f

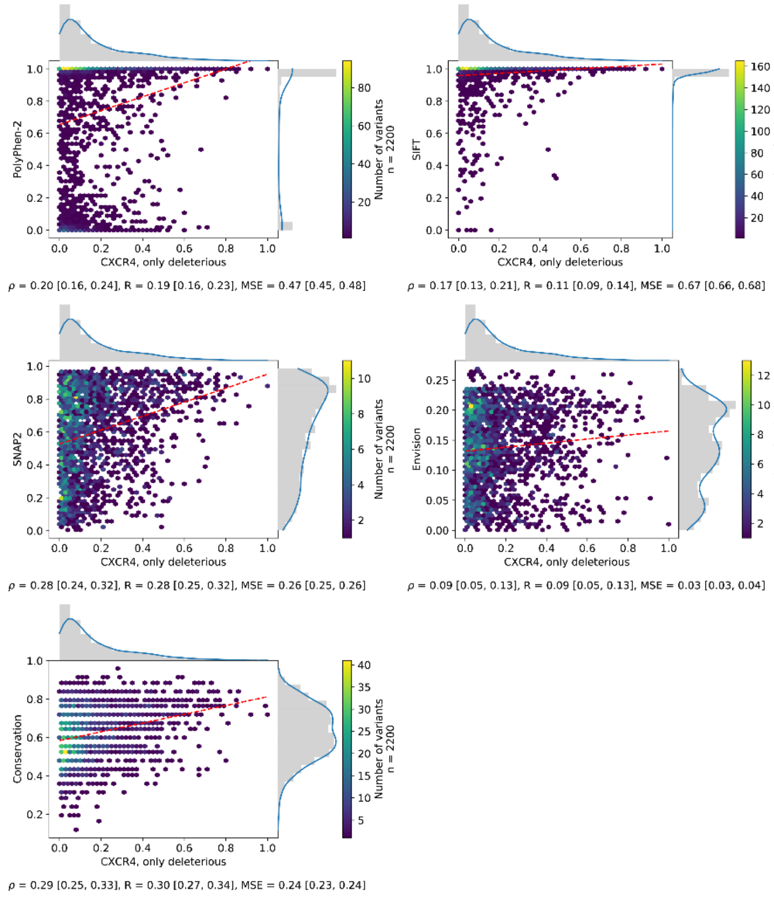
PolyPhen-2
CXCR4, only deleterious
Number of variants
n = 2200
ρ = 0.20 [0.16, 0.24], R = 0.19 [0.16, 0.23], MSE = 0.47 [0.45, 0.48]
SIFT
CXCR4, only deleterious
Number of variants
n = 2200
ρ = 0.17 [0.13, 0.21], R = 0.11 [0.09, 0.14], MSE = 0.67 [0.66, 0.68]
SNAP2
CXCR4, only deleterious
Number of variants
n = 2200
ρ = 0.28 [0.24, 0.32], R = 0.28 [0.25, 0.32], MSE = 0.26 [0.25, 0.26]
Envision
CXCR4, only deleterious
Number of variants
n = 2200
ρ = 0.09 [0.05, 0.13], R = 0.09 [0.05, 0.13], MSE = 0.03 [0.03, 0.04]
Conservation
CXCR4, only deleterious
Number of variants
n = 2200
ρ = 0.29 [0.25, 0.33], R = 0.30 [0.27, 0.34], MSE = 0.24 [0.23, 0.24]

## S3g

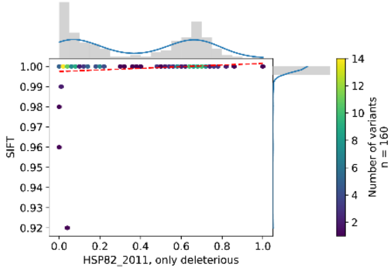

ρ = 0.27 [nan, nan], R = 0.16 [nan, nan], MSE = 0.48 [0.42, 0.53]

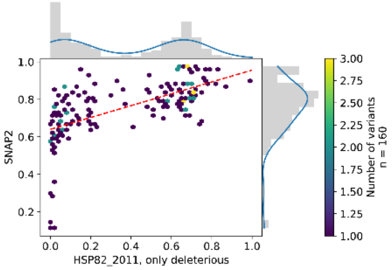

ρ = 0.69 [0.59, 0.77], R = 0.61 [0.54, 0.68], MSE = 0.20 [0.17, 0.23]

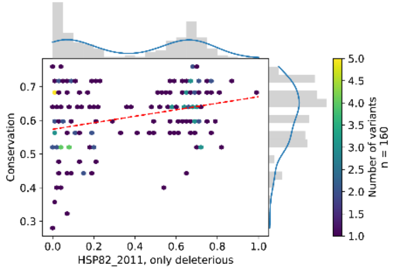

ρ = 0.25 [0.08, 0.39], R = 0.31 [0.16, 0.44], MSE = 0.13 [0.11, 0.16]

## S3h

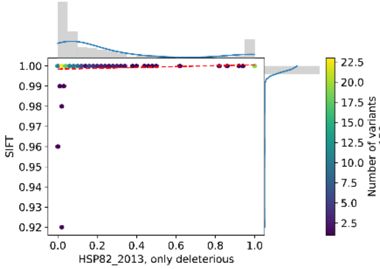

ρ = 0.22 [nan, nan], R = 0.10 [nan, nan], MSE = 0.68 [0.62, 0.73]

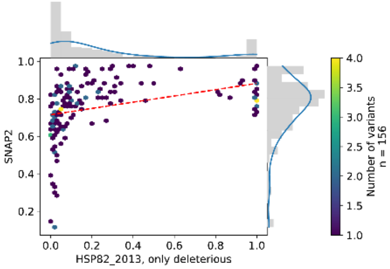

ρ = 0.61 [0.50, 0.70], R = 0.34 [0.27, 0.41], MSE = 0.36 [0.33, 0.39]

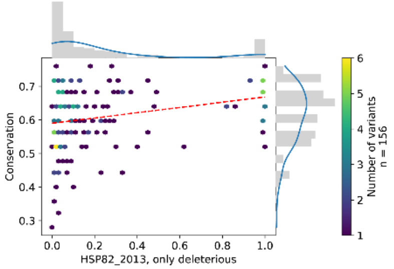

ρ = 0.28 [0.12, 0.42], R = 0.28 [0.16, 0.38], MSE = 0.23 [0.21, 0.25]

## S3i

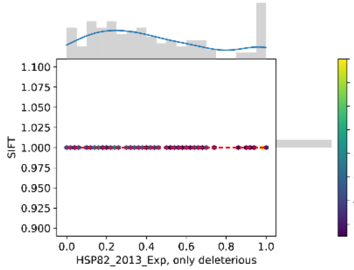

$\rho$ = nan [nan, nan], R = nan [nan, nan], MSE = 0.42 [0.37, 0.47]

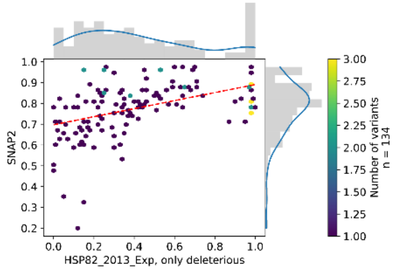

$\rho$ = 0.52 [0.38, 0.63], R = 0.44 [0.34, 0.55], MSE = 0.20 [0.17, 0.23]

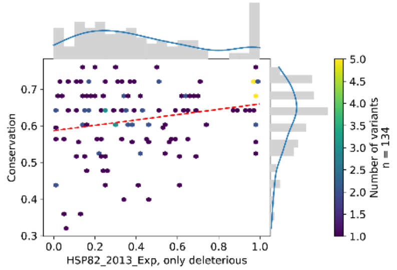

$\rho$ = 0.19 [0.02, 0.37], R = 0.24 [0.10, 0.38], MSE = 0.12 [0.10, 0.14]

## S3j

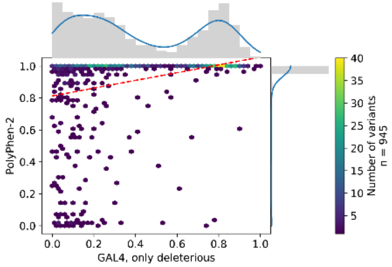

$\rho$ = 0.49 [0.44, 0.54], R = 0.32 [0.28, 0.36], MSE = 0.35 [0.33, 0.37]

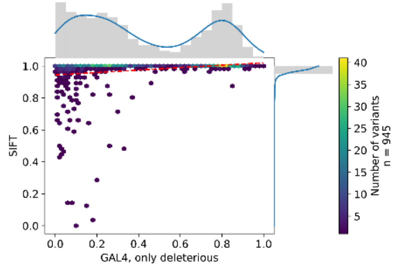

$\rho$ = 0.44 [0.38, 0.49], R = 0.23 [0.21, 0.27], MSE = 0.40 [0.38, 0.42]

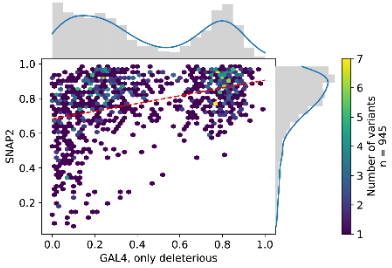

$\rho$ = 0.39 [0.33, 0.44], R = 0.39 [0.34, 0.43], MSE = 0.21 [0.20, 0.22]

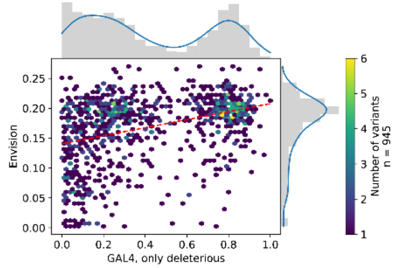

$\rho$ = 0.37 [0.31, 0.43], R = 0.37 [0.32, 0.42], MSE = 0.15 [0.14, 0.16]

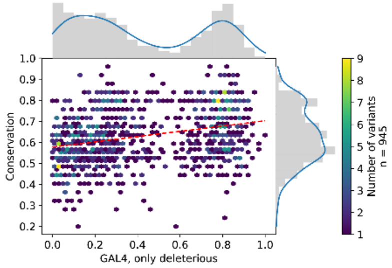

$\rho$ = 0.30 [0.24, 0.35], R = 0.29 [0.24, 0.35], MSE = 0.13 [0.12, 0.14]

## S3k

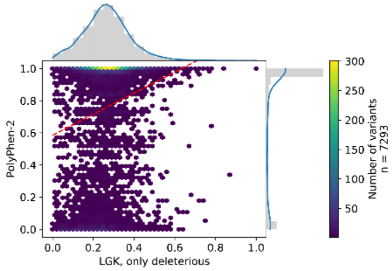

ρ = 0.20 [0.17, 0.22], R = 0.20 [0.18, 0.23], MSE = 0.37 [0.37, 0.38]

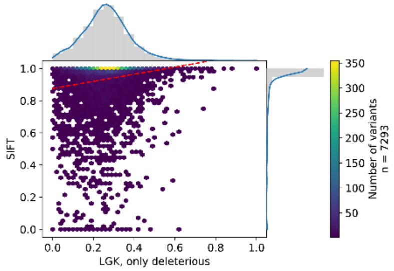

ρ = 0.20 [0.18, 0.22], R = 0.16 [0.14, 0.19], MSE = 0.48 [0.48, 0.48]

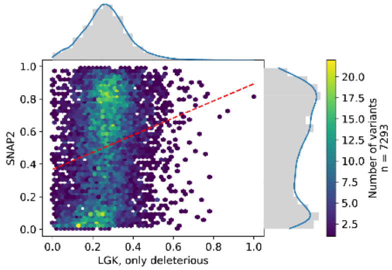

ρ = 0.21 [0.19, 0.23], R = 0.20 [0.18, 0.23], MSE = 0.14 [0.14, 0.15]

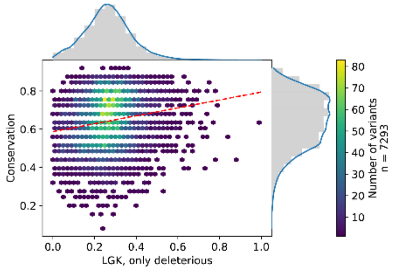

ρ = 0.19 [0.17, 0.21], R = 0.18 [0.15, 0.20], MSE = 0.17 [0.16, 0.17]

## S3l

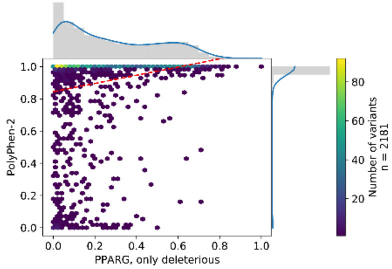

ρ = 0.37 [0.34, 0.41], R = 0.25 [0.21, 0.27], MSE = 0.48 [0.47, 0.50]

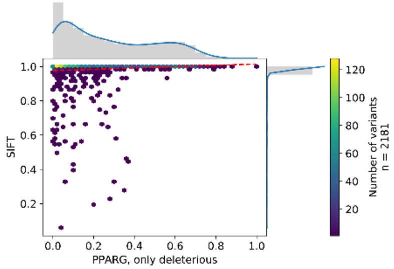

ρ = 0.25 [0.21, 0.28], R = 0.13 [0.10, 0.16], MSE = 0.55 [0.54, 0.56]

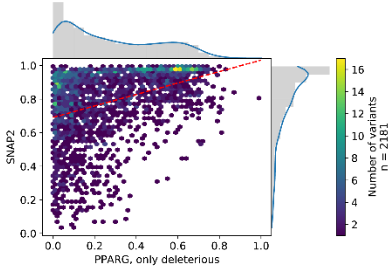

ρ = 0.42 [0.38, 0.45], R = 0.38 [0.35, 0.41], MSE = 0.31 [0.31, 0.32]

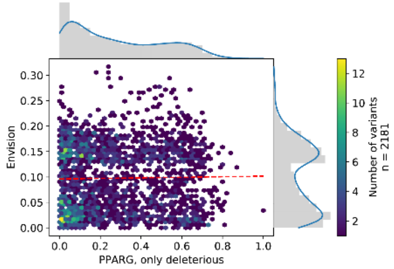

ρ = 0.02 [-0.02, 0.06], R = 0.02 [-0.02, 0.06], MSE = 0.09 [0.08, 0.09]

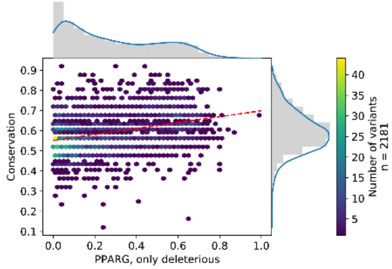

ρ = 0.34 [0.30, 0.38], R = 0.33 [0.29, 0.36], MSE = 0.14 [0.14, 0.15]

## S3m

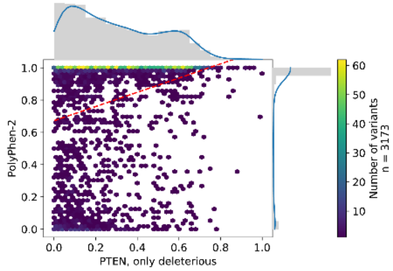

$\rho$ = 0.33 [0.30, 0.36], R = 0.28 [0.25, 0.31], MSE = 0.37 [0.36, 0.38]

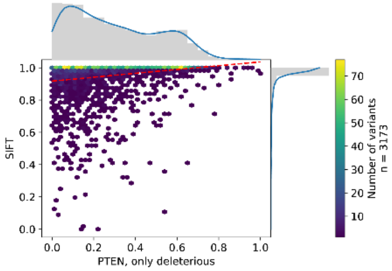

$\rho$ = 0.34 [0.31, 0.37], R = 0.23 [0.20, 0.25], MSE = 0.47 [0.46, 0.48]

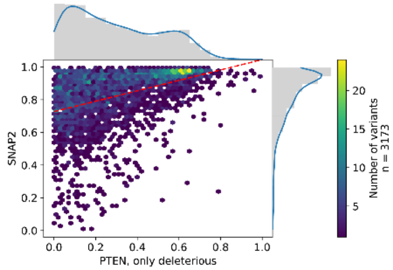

$\rho$ = 0.48 [0.45, 0.51], R = 0.41 [0.39, 0.43], MSE = 0.32 [0.31, 0.32]

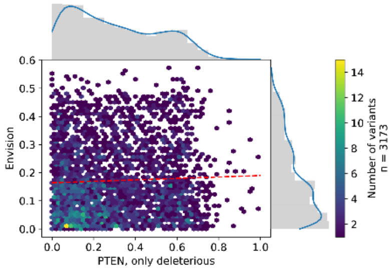

$\rho$ = 0.03 [-0.00, 0.07], R = 0.04 [0.01, 0.08], MSE = 0.08 [0.07, 0.08]

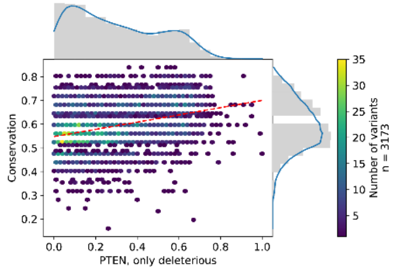

$\rho$ = 0.33 [0.30, 0.36], R = 0.33 [0.29, 0.35], MSE = 0.13 [0.12, 0.13]

## S3n

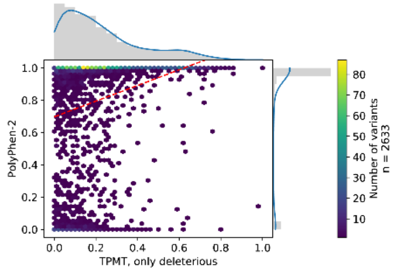

$\rho$ = 0.40 [0.37, 0.44], R = 0.27 [0.24, 0.30], MSE = 0.45 [0.44, 0.47]

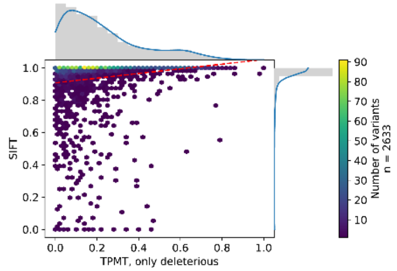

$\rho$ = 0.34 [0.30, 0.37], R = 0.17 [0.15, 0.20], MSE = 0.56 [0.55, 0.57]

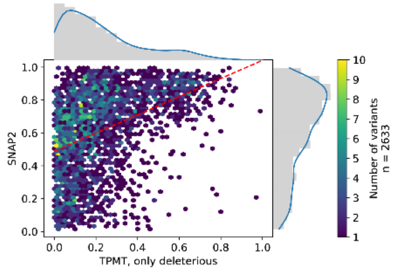

$\rho$ = 0.46 [0.43, 0.49], R = 0.42 [0.39, 0.45], MSE = 0.21 [0.20, 0.22]

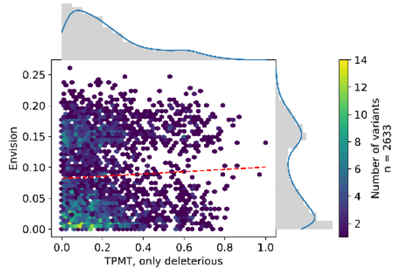

$\rho$ = 0.03 [-0.01, 0.07], R = 0.05 [0.02, 0.09], MSE = 0.06 [0.06, 0.06]

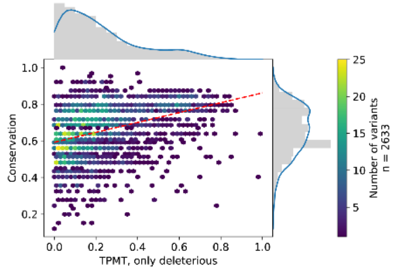

$\rho$ = 0.42 [0.39, 0.45], R = 0.39 [0.35, 0.42], MSE = 0.22 [0.21, 0.22]

S3o

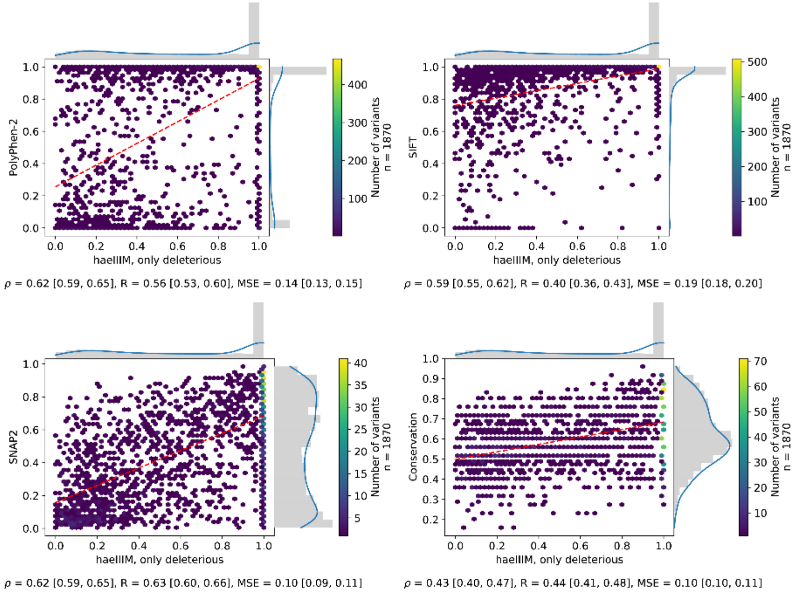
PolyPhen-2
haeIIIM, only deleterious
Number of variants
n = 1870
ρ = 0.62 [0.59, 0.65], R = 0.56 [0.53, 0.60], MSE = 0.14 [0.13, 0.15]
SIFT
haeIIIM, only deleterious
Number of variants
n = 1870
ρ = 0.59 [0.55, 0.62], R = 0.40 [0.36, 0.43], MSE = 0.19 [0.18, 0.20]
SNAP2
haeIIIM, only deleterious
Number of variants
n = 1870
ρ = 0.62 [0.59, 0.65], R = 0.63 [0.60, 0.66], MSE = 0.10 [0.09, 0.11]
Conservation
haeIIIM, only deleterious
Number of variants
n = 1870
ρ = 0.43 [0.40, 0.47], R = 0.44 [0.41, 0.48], MSE = 0.10 [0.10, 0.11]

S3p

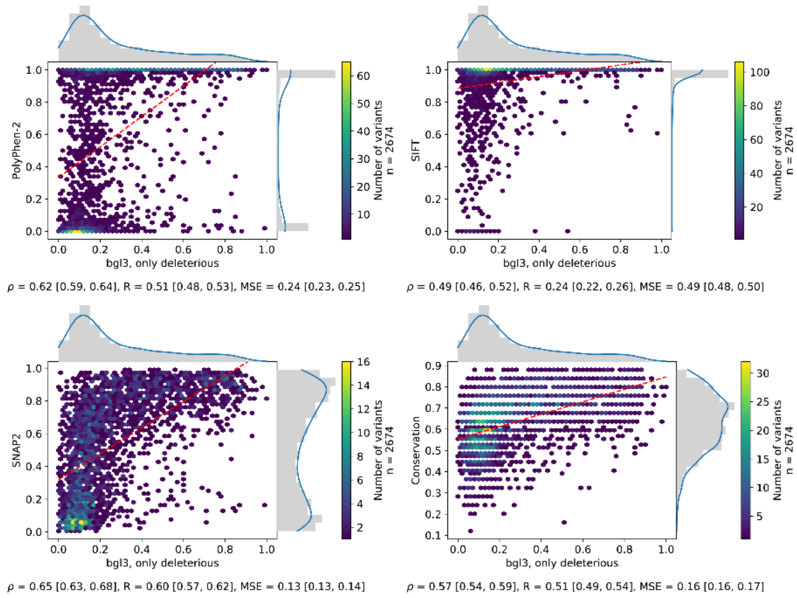
PolyPhen-2
bgl3, only deleterious
Number of variants
n = 2674
ρ = 0.62 [0.59, 0.64], R = 0.51 [0.48, 0.53], MSE = 0.24 [0.23, 0.25]
SIFT
bgl3, only deleterious
Number of variants
n = 2674
ρ = 0.49 [0.46, 0.52], R = 0.24 [0.22, 0.26], MSE = 0.49 [0.48, 0.50]
SNAP2
bgl3, only deleterious
Number of variants
n = 2674
ρ = 0.65 [0.63, 0.68], R = 0.60 [0.57, 0.62], MSE = 0.13 [0.13, 0.14]
Conservation
bgl3, only deleterious
Number of variants
n = 2674
ρ = 0.57 [0.54, 0.59], R = 0.51 [0.49, 0.54], MSE = 0.16 [0.16, 0.17]

## S3q

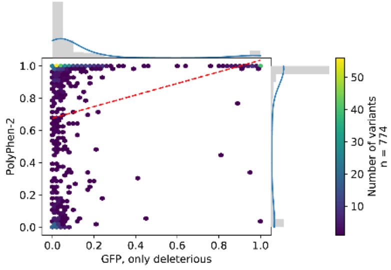

ρ = 0.49 [0.44, 0.55], R = 0.28 [0.24, 0.31], MSE = 0.49 [0.46, 0.52]

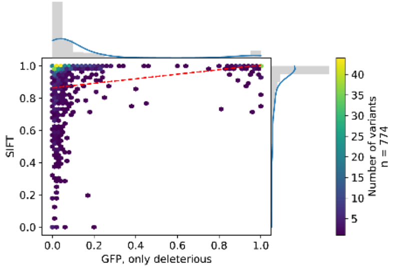

ρ = 0.40 [0.34, 0.46], R = 0.21 [0.18, 0.24], MSE = 0.62 [0.60, 0.64]

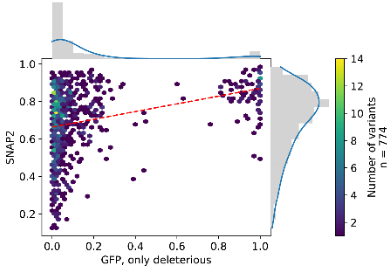

ρ = 0.36 [0.29, 0.43], R = 0.33 [0.29, 0.37], MSE = 0.37 [0.35, 0.38]

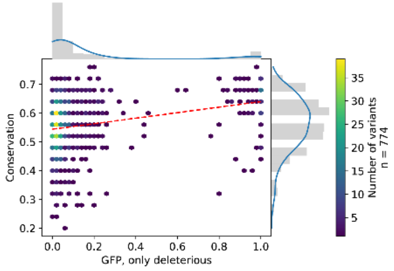

ρ = 0.32 [0.25, 0.38], R = 0.30 [0.23, 0.36], MSE = 0.23 [0.23, 0.24]

## S3r

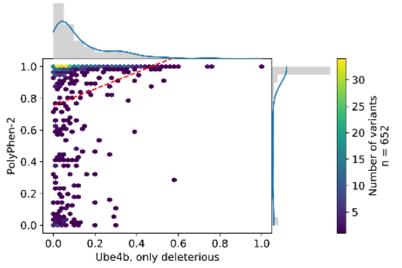

ρ = 0.36 [0.29, 0.43], R = 0.24 [0.19, 0.29], MSE = 0.58 [0.55, 0.60]

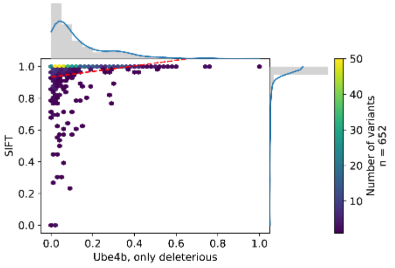

ρ = 0.40 [0.33, 0.45], R = 0.20 [0.18, 0.23], MSE = 0.71 [0.69, 0.73]

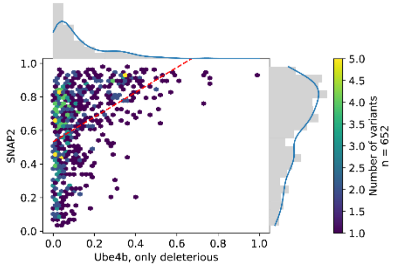

ρ = 0.48 [0.41, 0.54], R = 0.44 [0.39, 0.49], MSE = 0.30 [0.28, 0.31]

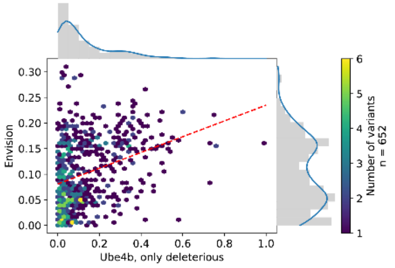

ρ = 0.32 [0.24, 0.39], R = 0.33 [0.27, 0.40], MSE = 0.02 [0.02, 0.02]

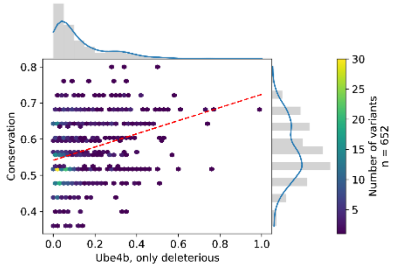

ρ = 0.29 [0.22, 0.36], R = 0.32 [0.24, 0.38], MSE = 0.21 [0.20, 0.22]

## S3s

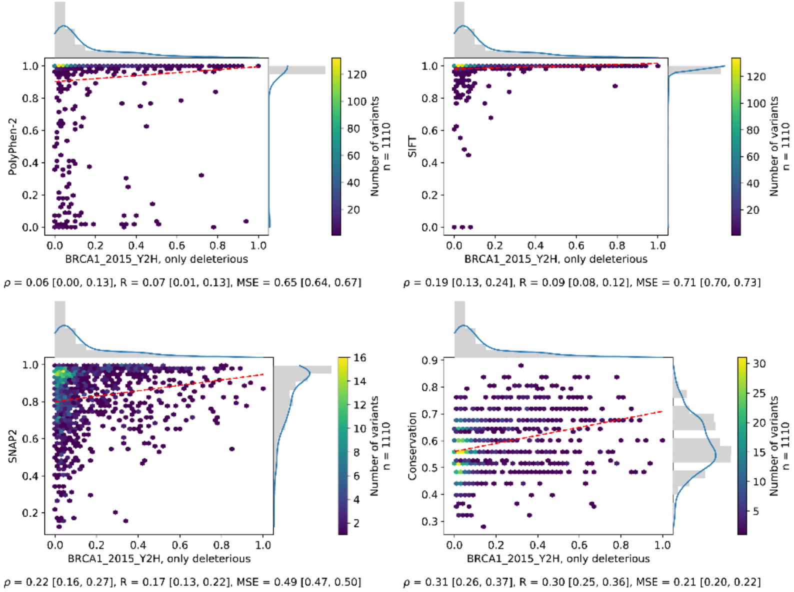
PolyPhen-2
BRCA1_2015_Y2H, only deleterious
Number of variants
n = 1110
ρ = 0.06 [0.00, 0.13], R = 0.07 [0.01, 0.13], MSE = 0.65 [0.64, 0.67]
SIFT
BRCA1_2015_Y2H, only deleterious
ρ = 0.19 [0.13, 0.24], R = 0.09 [0.08, 0.12], MSE = 0.71 [0.70, 0.73]
SNAP2
BRCA1_2015_Y2H, only deleterious
ρ = 0.22 [0.16, 0.27], R = 0.17 [0.13, 0.22], MSE = 0.49 [0.47, 0.50]
Conservation
BRCA1_2015_Y2H, only deleterious
ρ = 0.31 [0.26, 0.37], R = 0.30 [0.25, 0.36], MSE = 0.21 [0.20, 0.22]

## S3t

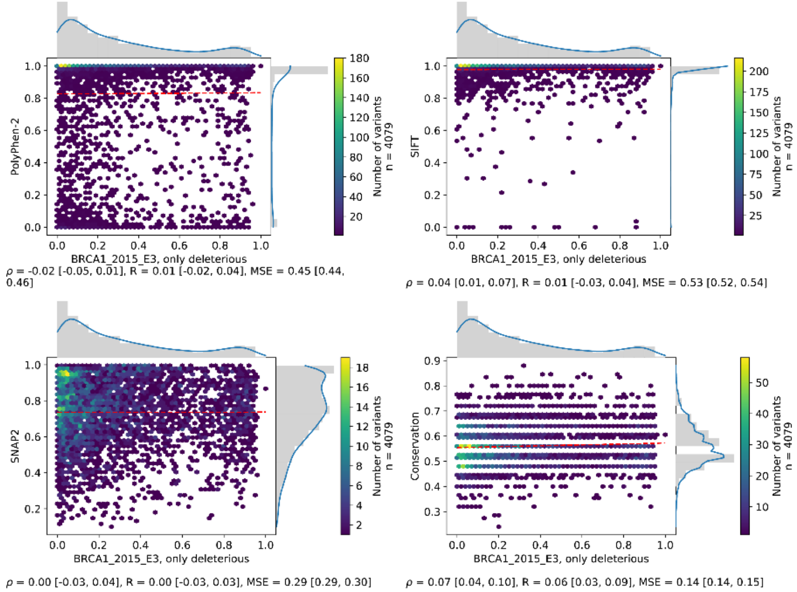
PolyPhen-2
BRCA1_2015_E3, only deleterious
Number of variants
n = 4079
ρ = -0.02 [-0.05, 0.01], R = 0.01 [-0.02, 0.04], MSE = 0.45 [0.44, 0.46]
SIFT
BRCA1_2015_E3, only deleterious
ρ = 0.04 [0.01, 0.07], R = 0.01 [-0.03, 0.04], MSE = 0.53 [0.52, 0.54]
SNAP2
BRCA1_2015_E3, only deleterious
ρ = 0.00 [-0.03, 0.04], R = 0.00 [-0.03, 0.03], MSE = 0.29 [0.29, 0.30]
Conservation
BRCA1_2015_E3, only deleterious
ρ = 0.07 [0.04, 0.10], R = 0.06 [0.03, 0.09], MSE = 0.14 [0.14, 0.15]

S3u

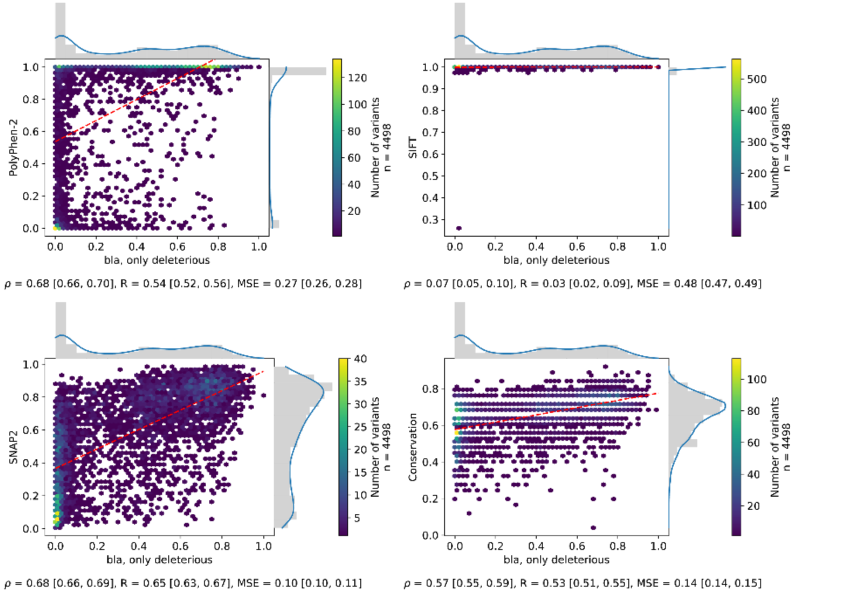
PolyPhen-2
SIFT
SNAP2
Conservation
bla, only deleterious
Number of variants
n = 4498
ρ = 0.68 [0.66, 0.70], R = 0.54 [0.52, 0.56], MSE = 0.27 [0.26, 0.28]
ρ = 0.07 [0.05, 0.10], R = 0.03 [0.02, 0.09], MSE = 0.48 [0.47, 0.49]
ρ = 0.68 [0.66, 0.69], R = 0.65 [0.63, 0.67], MSE = 0.10 [0.10, 0.11]
ρ = 0.57 [0.55, 0.59], R = 0.53 [0.51, 0.55], MSE = 0.14 [0.14, 0.15]

S3v

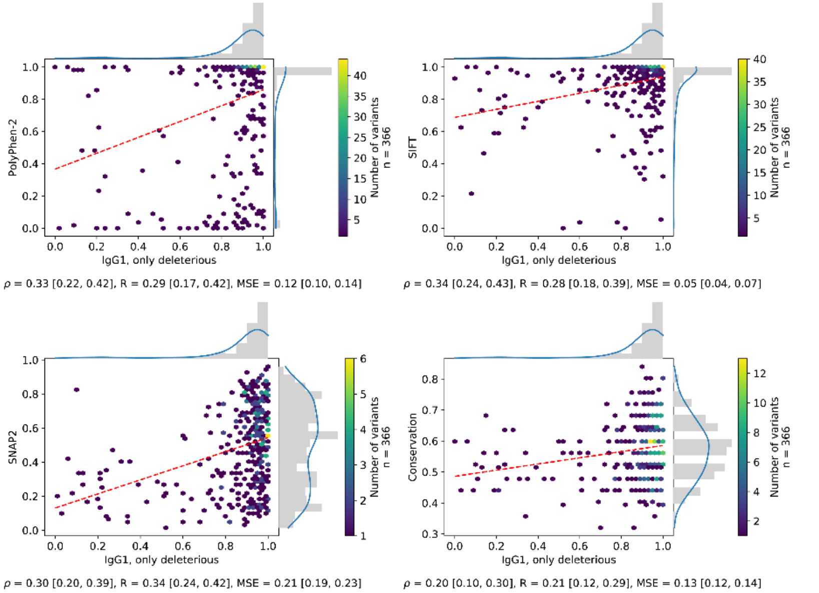
PolyPhen-2
SIFT
SNAP2
Conservation
IgG1, only deleterious
Number of variants
n = 366
ρ = 0.33 [0.22, 0.42], R = 0.29 [0.17, 0.42], MSE = 0.12 [0.10, 0.14]
ρ = 0.34 [0.24, 0.43], R = 0.28 [0.18, 0.39], MSE = 0.05 [0.04, 0.07]
ρ = 0.30 [0.20, 0.39], R = 0.34 [0.24, 0.42], MSE = 0.21 [0.19, 0.23]
ρ = 0.20 [0.10, 0.30], R = 0.21 [0.12, 0.29], MSE = 0.13 [0.12, 0.14]

#### Figure S4: Agreement between prediction methods and beneficial effect scores from experiments on SetCommon and SetCommonSyn95.

Related to Figure 1. Panels (a)-(e) show the performance for five prediction methods through hexbin plots of 15,200 beneficial effect SAVs in SetCommon (see Methods, Processing). Values on axes range from 0 (neutral) to 1 (maximal effect). Dashed red lines give linear least-squared regressions. Marginals denote distributions of experimental and predicted scores with a kernel density estimation overlaid in blue. The footer denotes Spearman ρ, Pearson R and the mean squared error together with the respective 95% confidence intervals. The method scores are given on the y-axes and reveal the method: (a) PolyPhen-2, (b) SIFT, (c) SNAP2, (d) Envision – the only method trained on DMS data, (e) naïve conservation read off PSI-BLAST profiles. Panel (f) gives ROC curves for 12,412 beneficial effect SAVs which were classified into either neutral, defined by the middle 95% of the scores from synonymous variants, or effect (SetCommonSyn95, see Methods). Shaded areas around lines denote 95% confidence intervals. The legend denotes the AUC for each method along with the 95% confidence intervals. Horizontal dashed lines denote the default score threshold used by SNAP2 (blue) and SIFT (green).

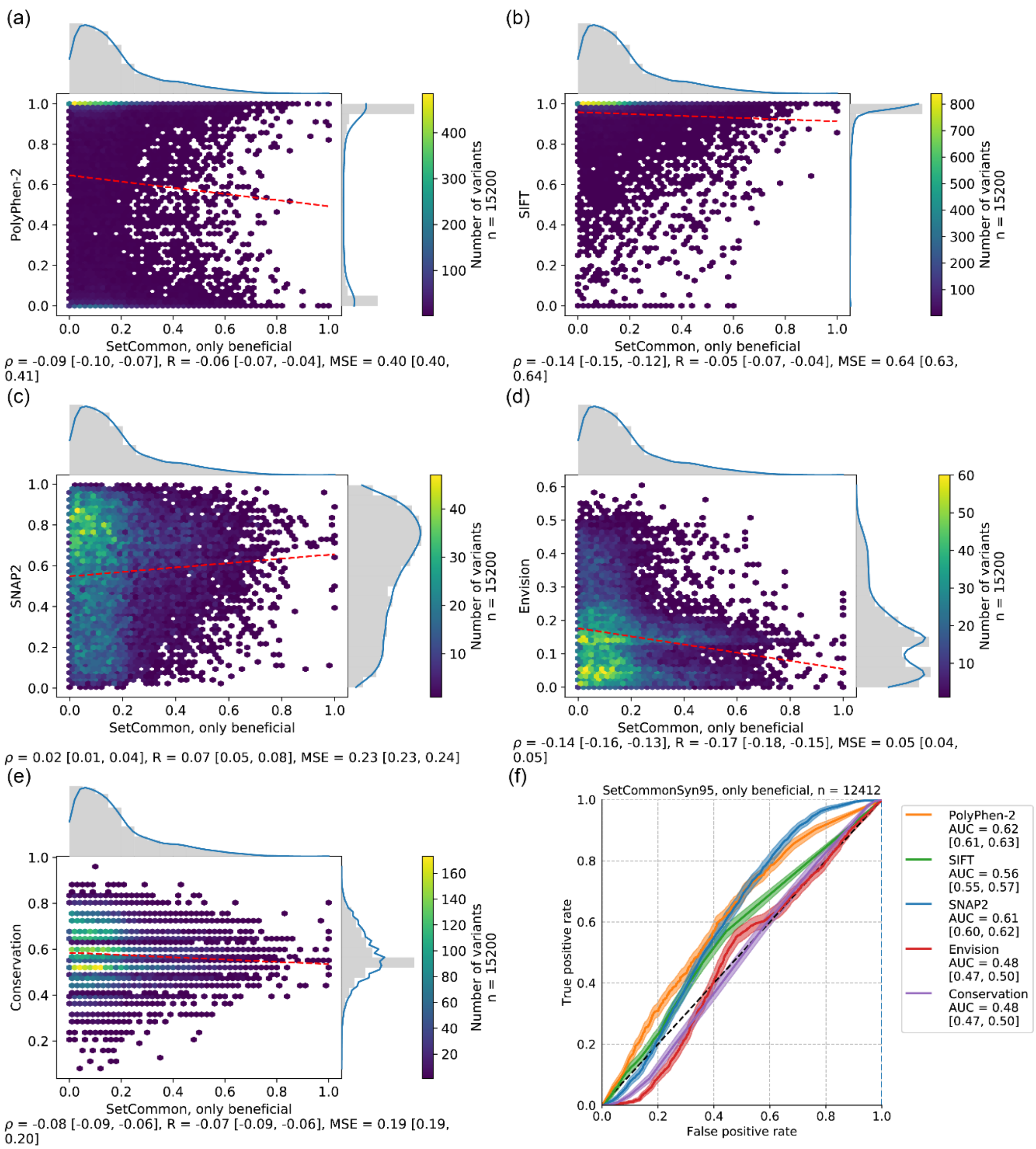
(a)
PolyPhen-2
SetCommon, only beneficial
Number of variants
n = 15200
ρ = -0.09 [-0.10, -0.07], R = -0.06 [-0.07, -0.04], MSE = 0.40 [0.40, 0.41]
(b)
SIFT
SetCommon, only beneficial
Number of variants
n = 15200
ρ = -0.14 [-0.15, -0.12], R = -0.05 [-0.07, -0.04], MSE = 0.64 [0.63, 0.64]
(c)
SNAP2
SetCommon, only beneficial
Number of variants
n = 15200
ρ = 0.02 [0.01, 0.04], R = 0.07 [0.05, 0.08], MSE = 0.23 [0.23, 0.24]
(d)
Envision
SetCommon, only beneficial
Number of variants
n = 15200
ρ = -0.14 [-0.16, -0.13], R = -0.17 [-0.18, -0.15], MSE = 0.05 [0.04, 0.05]
(e)
Conservation
SetCommon, only beneficial
Number of variants
n = 15200
ρ = -0.08 [-0.09, -0.06], R = -0.07 [-0.09, -0.06], MSE = 0.19 [0.19, 0.20]
(f)
SetCommonSyn95, only beneficial, n = 12412
True positive rate
False positive rate
PolyPhen-2 AUC = 0.62 [0.61, 0.63]
SIFT AUC = 0.56 [0.55, 0.57]
SNAP2 AUC = 0.61 [0.60, 0.62]
Envision AUC = 0.48 [0.47, 0.50]
Conservation AUC = 0.48 [0.47, 0.50]

#### Figure S5: Agreements between prediction methods and beneficial effect scores from experiments.

Panels (a)-(u) show the performance for five prediction methods through hexbin plots for all 21 DMS datasets in SetAll (see Fig. S1b, Table S1, Methods). For every dataset only the largest common subset of beneficial effect SAVs for which a prediction was available from every method is analyzed. Missing methods did not perform any predictions at all. The ccdb set was excluded as it does not contain any beneficial effect SAVs. Values on axes range from 0 (neutral) to 1 (maximal effect). Dashed red lines give linear least-squared regressions. Marginals denote distributions of experimental and predicted scores with a kernel density estimation overlaid in blue. The footer denotes Spearman ρ, Pearson R and the mean squared error together with the respective 95% confidence intervals. The method scores are given on the y-axes and reveal the methods: PolyPhen-2, SIFT, SNAP2, Envision (the only method trained on DMS data), naïve conservation read off PSI-BLAST profiles.

S5a

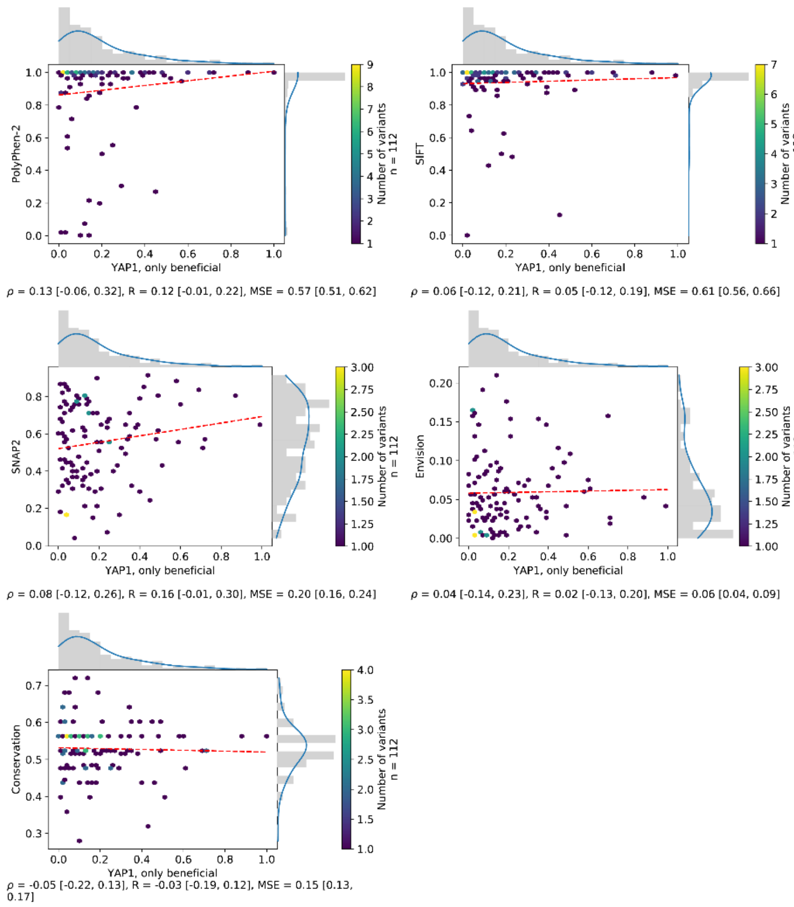
PolyPhen-2
SIFT
SNAP2
Envision
Conservation
YAP1, only beneficial
Number of variants
n = 112
ρ = 0.13 [-0.06, 0.32], R = 0.12 [-0.01, 0.22], MSE = 0.57 [0.51, 0.62]
ρ = 0.06 [-0.12, 0.21], R = 0.05 [-0.12, 0.19], MSE = 0.61 [0.56, 0.66]
ρ = 0.08 [-0.12, 0.26], R = 0.16 [-0.01, 0.30], MSE = 0.20 [0.16, 0.24]
ρ = 0.04 [-0.14, 0.23], R = 0.02 [-0.13, 0.20], MSE = 0.06 [0.04, 0.09]
ρ = -0.05 [-0.22, 0.13], R = -0.03 [-0.19, 0.12], MSE = 0.15 [0.13, 0.17]

S5b

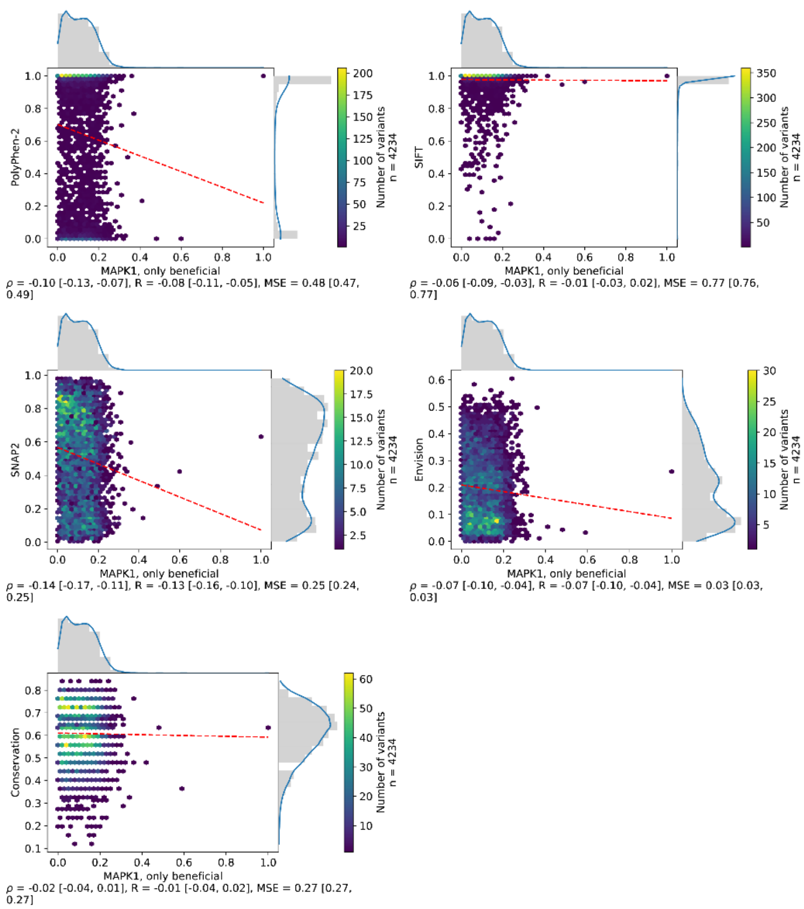
PolyPhen-2
SIFT
SNAP2
Envision
Conservation
MAPK1, only beneficial
Number of variants
n = 4234
ρ = -0.10 [-0.13, -0.07], R = -0.08 [-0.11, -0.05], MSE = 0.48 [0.47, 0.49]
ρ = -0.06 [-0.09, -0.03], R = -0.01 [-0.03, 0.02], MSE = 0.77 [0.76, 0.77]
ρ = -0.14 [-0.17, -0.11], R = -0.13 [-0.16, -0.10], MSE = 0.25 [0.24, 0.25]
ρ = -0.07 [-0.10, -0.04], R = -0.07 [-0.10, -0.04], MSE = 0.03 [0.03, 0.03]
ρ = -0.02 [-0.04, 0.01], R = -0.01 [-0.04, 0.02], MSE = 0.27 [0.27, 0.27]

## S5c

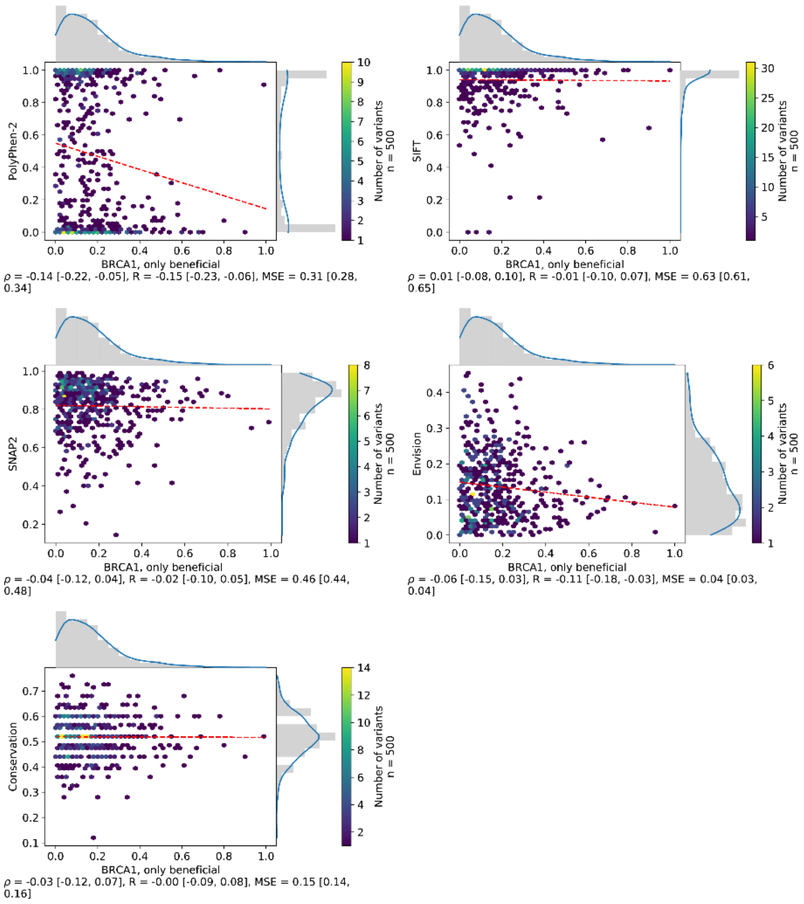
PolyPhen-2
BRCA1, only beneficial
Number of variants n = 500
ρ = -0.14 [-0.22, -0.05], R = -0.15 [-0.23, -0.06], MSE = 0.31 [0.28, 0.34]
SIFT
BRCA1, only beneficial
Number of variants n = 500
ρ = 0.01 [-0.08, 0.10], R = -0.01 [-0.10, 0.07], MSE = 0.63 [0.61, 0.65]
SNAP2
BRCA1, only beneficial
Number of variants n = 500
ρ = -0.04 [-0.12, 0.04], R = -0.02 [-0.10, 0.05], MSE = 0.46 [0.44, 0.48]
Envision
BRCA1, only beneficial
Number of variants n = 500
ρ = -0.06 [-0.15, 0.03], R = -0.11 [-0.18, -0.03], MSE = 0.04 [0.03, 0.04]
Conservation
BRCA1, only beneficial
Number of variants n = 500
ρ = -0.03 [-0.12, 0.07], R = -0.00 [-0.09, 0.08], MSE = 0.15 [0.14, 0.16]

## S5d

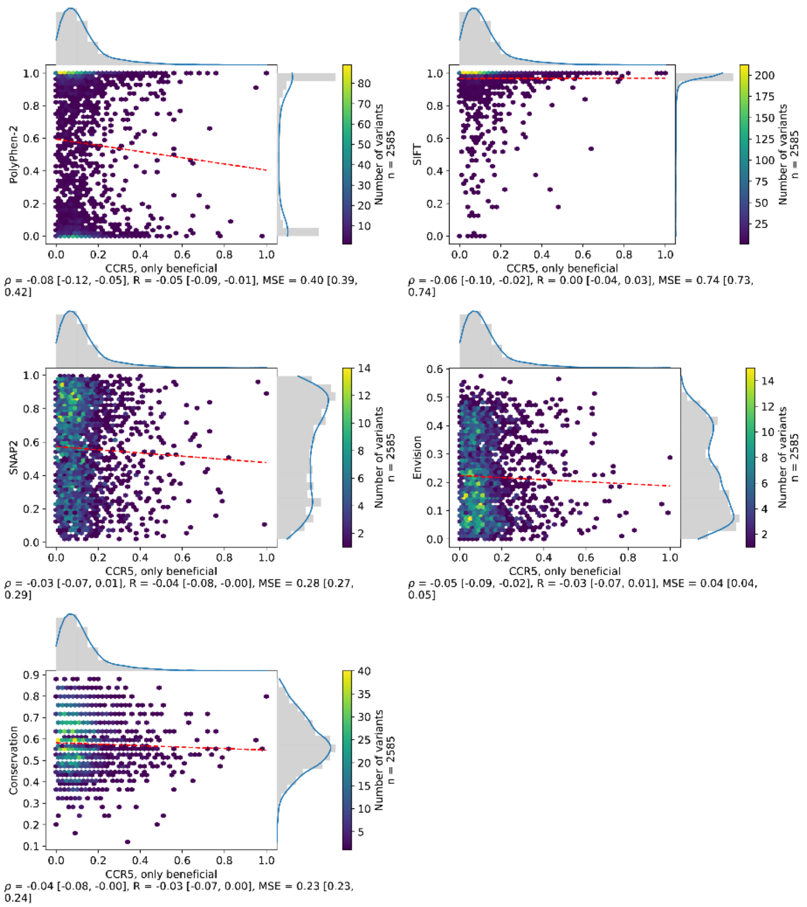
PolyPhen-2
CCR5, only beneficial
Number of variants n = 2585
ρ = -0.08 [-0.12, -0.05], R = -0.05 [-0.09, -0.01], MSE = 0.40 [0.39, 0.42]
SIFT
CCR5, only beneficial
Number of variants n = 2585
ρ = -0.06 [-0.10, -0.02], R = 0.00 [-0.04, 0.03], MSE = 0.74 [0.73, 0.74]
SNAP2
CCR5, only beneficial
Number of variants n = 2585
ρ = -0.03 [-0.07, 0.01], R = -0.04 [-0.08, -0.00], MSE = 0.28 [0.27, 0.29]
Envision
CCR5, only beneficial
Number of variants n = 2585
ρ = -0.05 [-0.09, -0.02], R = -0.03 [-0.07, 0.01], MSE = 0.04 [0.04, 0.05]
Conservation
CCR5, only beneficial
Number of variants n = 2585
ρ = -0.04 [-0.08, -0.00], R = -0.03 [-0.07, 0.00], MSE = 0.23 [0.23, 0.24]

## S5e

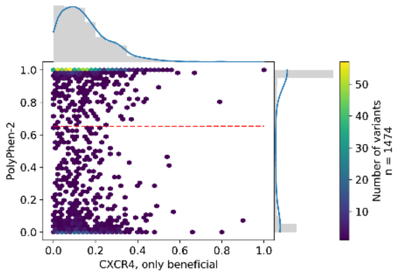

ρ = 0.01 [-0.04, 0.06], R = 0.00 [-0.05, 0.05], MSE = 0.44 [0.42, 0.46]

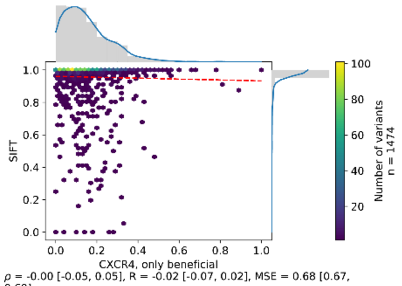

ρ = -0.00 [-0.05, 0.05], R = -0.02 [-0.07, 0.02], MSE = 0.68 [0.67, 0.69]

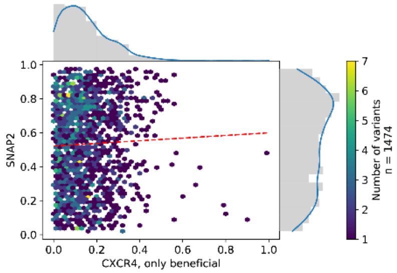

ρ = 0.04 [-0.01, 0.09], R = 0.04 [-0.02, 0.09], MSE = 0.23 [0.22, 0.24]

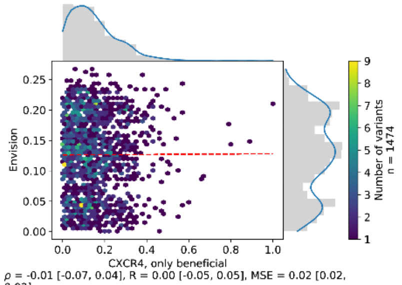

ρ = -0.01 [-0.07, 0.04], R = 0.00 [-0.05, 0.05], MSE = 0.02 [0.02, 0.02]

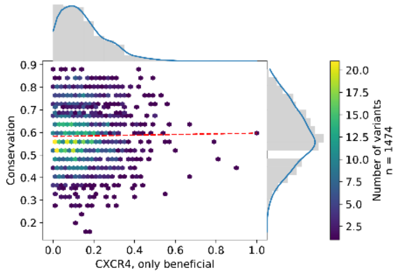

ρ = 0.02 [-0.03, 0.07], R = 0.01 [-0.04, 0.06], MSE = 0.22 [0.21, 0.23]

## S5f

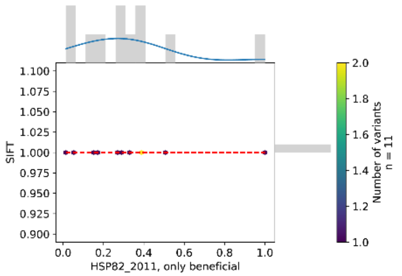

ρ = nan [nan, nan], R = nan [nan, nan], MSE = 0.52 [0.36, 0.67]

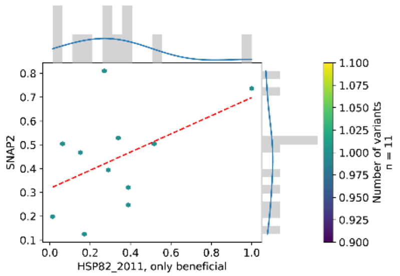

ρ = 0.36 [-0.33, 0.86], R = 0.48 [-0.21, 0.86], MSE = 0.07 [0.02, 0.13]

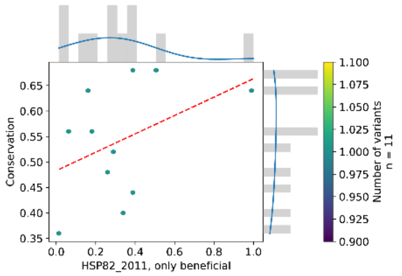

ρ = 0.46 [-0.37, 0.84], R = 0.43 [-0.32, 0.75], MSE = 0.10 [0.05, 0.15]

## S5g

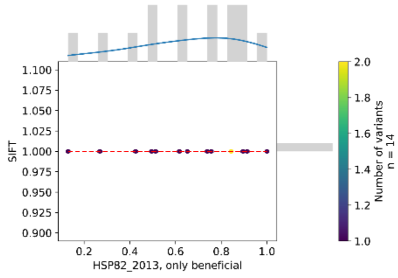

$\rho$ = nan [nan, nan], R = nan [nan, nan], MSE = 0.19 [0.08, 0.32]

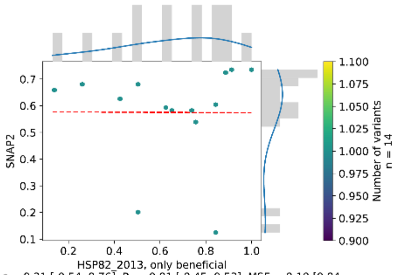

$\rho$ = 0.21 [-0.54, 0.76], R = -0.01 [-0.45, 0.53], MSE = 0.10 [0.04, 0.18]

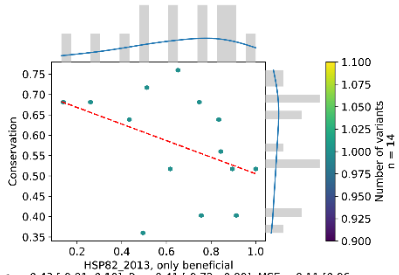

$\rho$ = -0.43 [-0.81, 0.10], R = -0.41 [-0.72, -0.00], MSE = 0.11 [0.06, 0.16]

## S5h

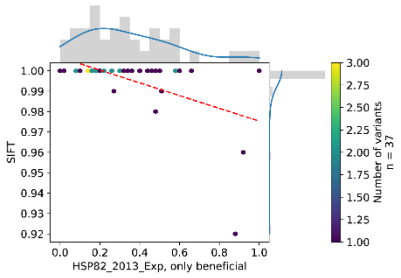

$\rho$ = -0.38 [nan, nan], R = -0.52 [nan, nan], MSE = 0.48 [0.41, 0.58]

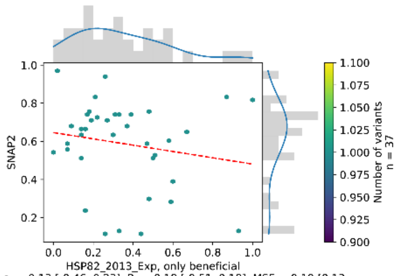

$\rho$ = -0.13 [-0.46, 0.23], R = -0.18 [-0.51, 0.18], MSE = 0.19 [0.13, 0.25]

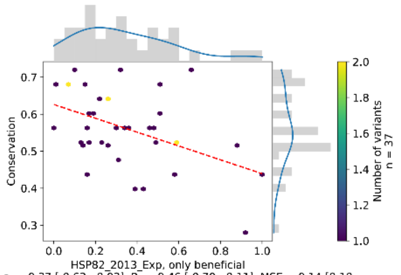

$\rho$ = -0.37 [-0.63, -0.03], R = -0.46 [-0.70, -0.11], MSE = 0.14 [0.10, 0.18]

## S5i

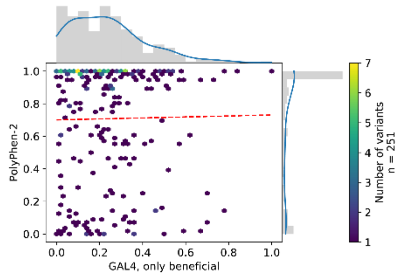

$\rho$ = 0.02 [-0.10, 0.15], R = 0.02 [-0.12, 0.14], MSE = 0.38 [0.34, 0.42]

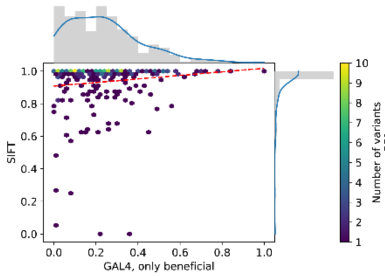

$\rho$ = 0.12 [-0.01, 0.24], R = 0.13 [0.03, 0.23], MSE = 0.53 [0.49, 0.56]

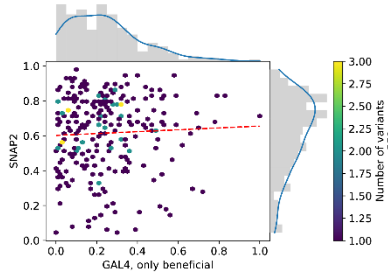

$\rho$ = 0.07 [-0.06, 0.19], R = 0.05 [-0.07, 0.16], MSE = 0.21 [0.19, 0.24]

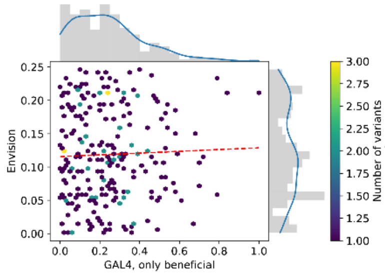

$\rho$ = 0.02 [-0.10, 0.13], R = 0.04 [-0.10, 0.15], MSE = 0.05 [0.04, 0.06]

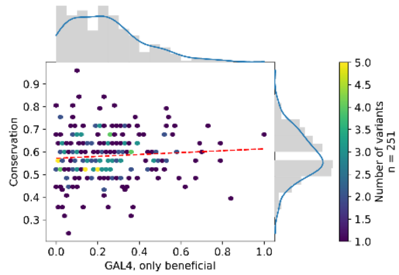

$\rho$ = 0.09 [-0.04, 0.21], R = 0.07 [-0.06, 0.20], MSE = 0.16 [0.14, 0.17]

## S5j

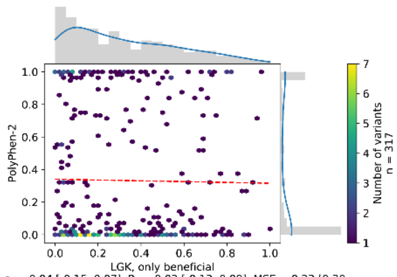

$\rho$ = -0.04 [-0.15, 0.07], R = -0.02 [-0.13, 0.09], MSE = 0.23 [0.20, 0.26]

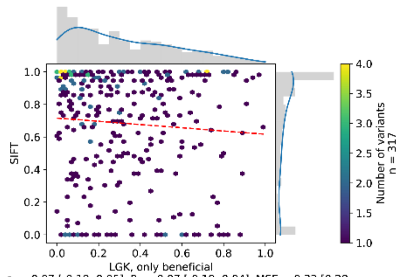

$\rho$ = -0.07 [-0.19, 0.05], R = -0.07 [-0.19, 0.04], MSE = 0.32 [0.28, 0.35]

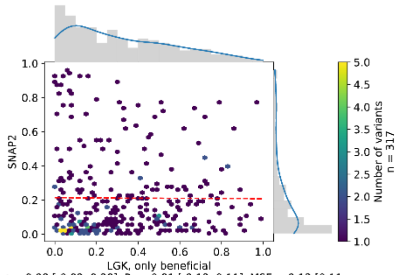

$\rho$ = 0.08 [-0.02, 0.20], R = -0.01 [-0.12, 0.11], MSE = 0.13 [0.11, 0.16]

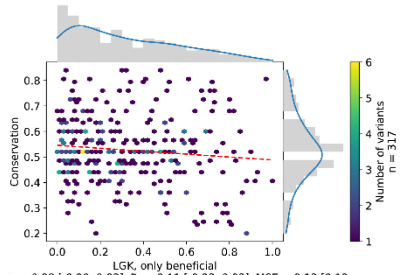

$\rho$ = -0.08 [-0.20, 0.03], R = -0.11 [-0.23, 0.03], MSE = 0.13 [0.12, 0.14]

S5k

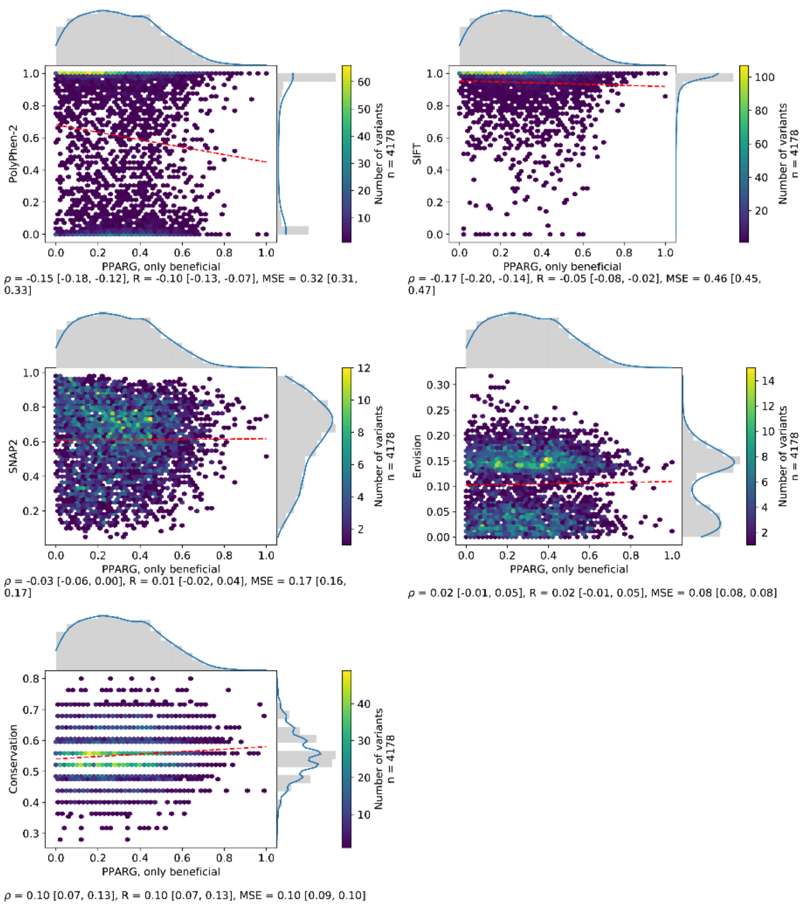
PolyPhen-2
SIFT
SNAP2
Envision
Conservation
PPARG, only beneficial
Number of variants
n = 4178
ρ = -0.15 [-0.18, -0.12], R = -0.10 [-0.13, -0.07], MSE = 0.32 [0.31, 0.33]
ρ = -0.17 [-0.20, -0.14], R = -0.05 [-0.08, -0.02], MSE = 0.46 [0.45, 0.47]
ρ = -0.03 [-0.06, 0.00], R = 0.01 [-0.02, 0.04], MSE = 0.17 [0.16, 0.17]
ρ = 0.02 [-0.01, 0.05], R = 0.02 [-0.01, 0.05], MSE = 0.08 [0.08, 0.08]
ρ = 0.10 [0.07, 0.13], R = 0.10 [0.07, 0.13], MSE = 0.10 [0.09, 0.10]

S5l

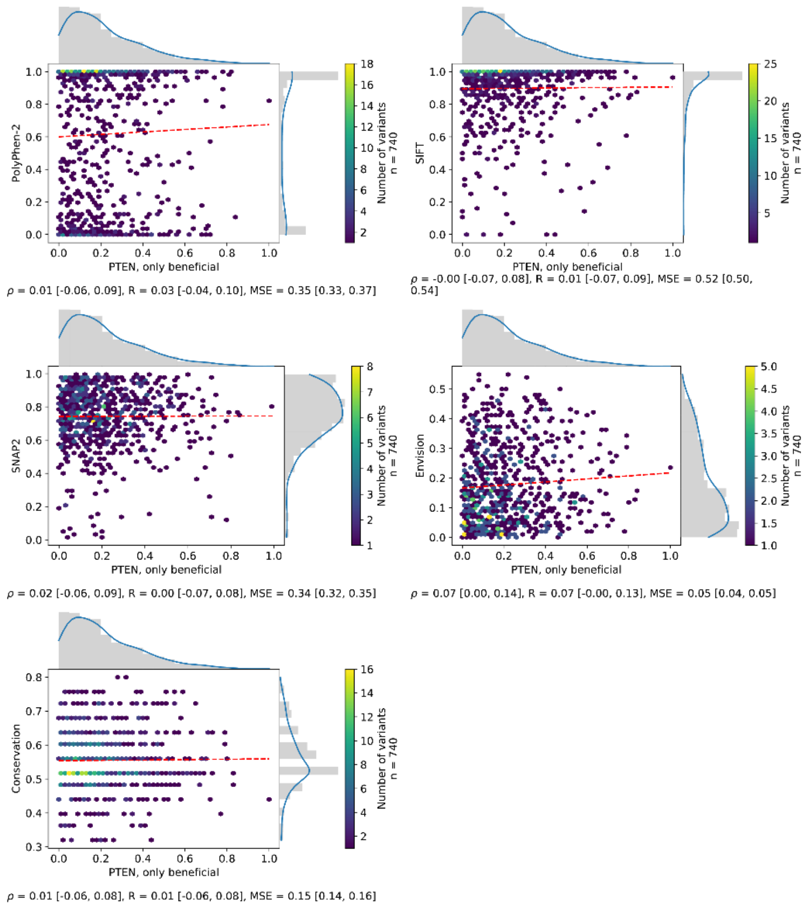
PolyPhen-2
SIFT
SNAP2
Envision
Conservation
PTEN, only beneficial
Number of variants
n = 740
ρ = 0.01 [-0.06, 0.09], R = 0.03 [-0.04, 0.10], MSE = 0.35 [0.33, 0.37]
ρ = -0.00 [-0.07, 0.08], R = 0.01 [-0.07, 0.09], MSE = 0.52 [0.50, 0.54]
ρ = 0.02 [-0.06, 0.09], R = 0.00 [-0.07, 0.08], MSE = 0.34 [0.32, 0.35]
ρ = 0.07 [0.00, 0.14], R = 0.07 [-0.00, 0.13], MSE = 0.05 [0.04, 0.05]
ρ = 0.01 [-0.06, 0.08], R = 0.01 [-0.06, 0.08], MSE = 0.15 [0.14, 0.16]

## S5m

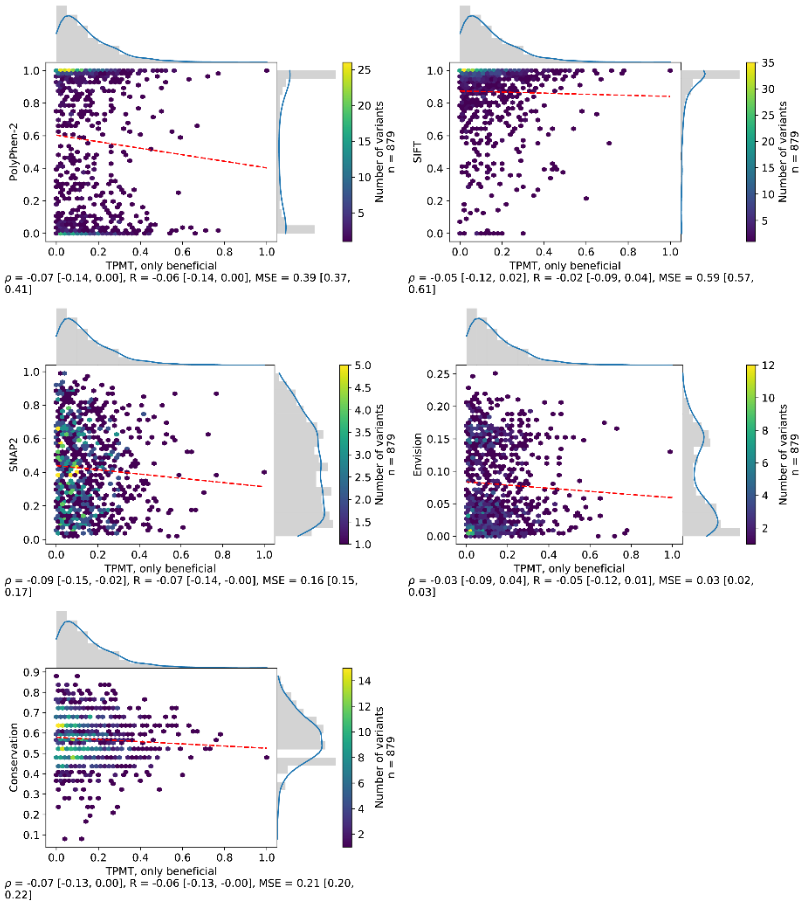
PolyPhen-2
TPMT, only beneficial
Number of variants n = 879
ρ = -0.07 [-0.14, 0.00], R = -0.06 [-0.14, 0.00], MSE = 0.39 [0.37, 0.41]
SIFT
TPMT, only beneficial
Number of variants n = 879
ρ = -0.05 [-0.12, 0.02], R = -0.02 [-0.09, 0.04], MSE = 0.59 [0.57, 0.61]
SNAP2
TPMT, only beneficial
Number of variants n = 879
ρ = -0.09 [-0.15, -0.02], R = -0.07 [-0.14, -0.00], MSE = 0.16 [0.15, 0.17]
Envision
TPMT, only beneficial
Number of variants n = 879
ρ = -0.03 [-0.09, 0.04], R = -0.05 [-0.12, 0.01], MSE = 0.03 [0.02, 0.03]
Conservation
TPMT, only beneficial
Number of variants n = 879
ρ = -0.07 [-0.13, 0.00], R = -0.06 [-0.13, -0.00], MSE = 0.21 [0.20, 0.22]

## S5n

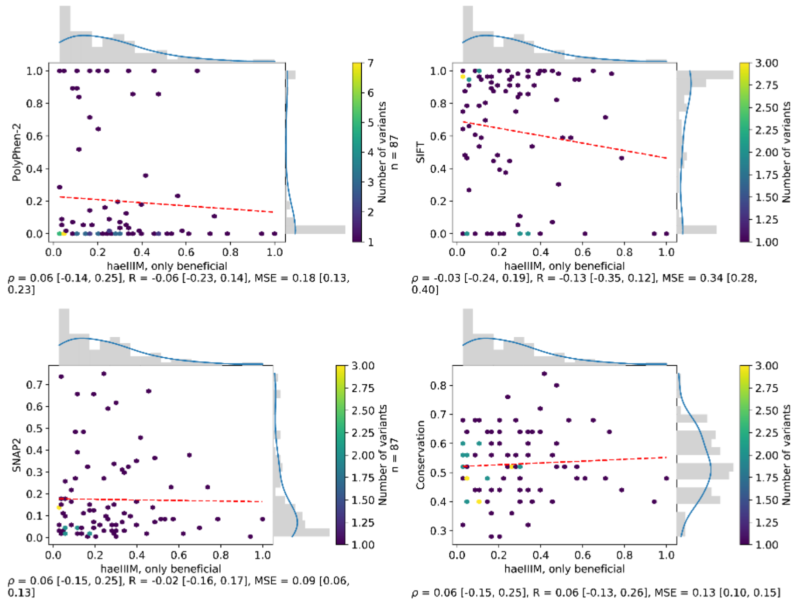
PolyPhen-2
haeIIIM, only beneficial
Number of variants n = 87
ρ = 0.06 [-0.14, 0.25], R = -0.06 [-0.23, 0.14], MSE = 0.18 [0.13, 0.23]
SIFT
haeIIIM, only beneficial
Number of variants n = 87
ρ = -0.03 [-0.24, 0.19], R = -0.13 [-0.35, 0.12], MSE = 0.34 [0.28, 0.40]
SNAP2
haeIIIM, only beneficial
Number of variants n = 87
ρ = 0.06 [-0.15, 0.25], R = -0.02 [-0.16, 0.17], MSE = 0.09 [0.06, 0.13]
Conservation
haeIIIM, only beneficial
Number of variants n = 87
ρ = 0.06 [-0.15, 0.25], R = 0.06 [-0.13, 0.26], MSE = 0.13 [0.10, 0.15]

S5o

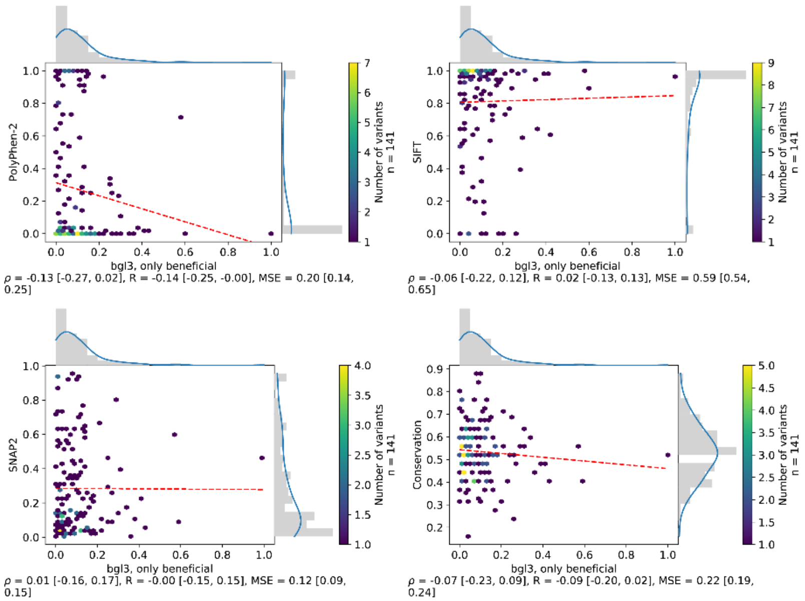
PolyPhen-2
bgl3, only beneficial
Number of variants
n = 141
ρ = -0.13 [-0.27, 0.02], R = -0.14 [-0.25, -0.00], MSE = 0.20 [0.14, 0.25]
SIFT
ρ = -0.06 [-0.22, 0.12], R = 0.02 [-0.13, 0.13], MSE = 0.59 [0.54, 0.65]
SNAP2
ρ = 0.01 [-0.16, 0.17], R = -0.00 [-0.15, 0.15], MSE = 0.12 [0.09, 0.15]
Conservation
ρ = -0.07 [-0.23, 0.09], R = -0.09 [-0.20, 0.02], MSE = 0.22 [0.19, 0.24]

S5p

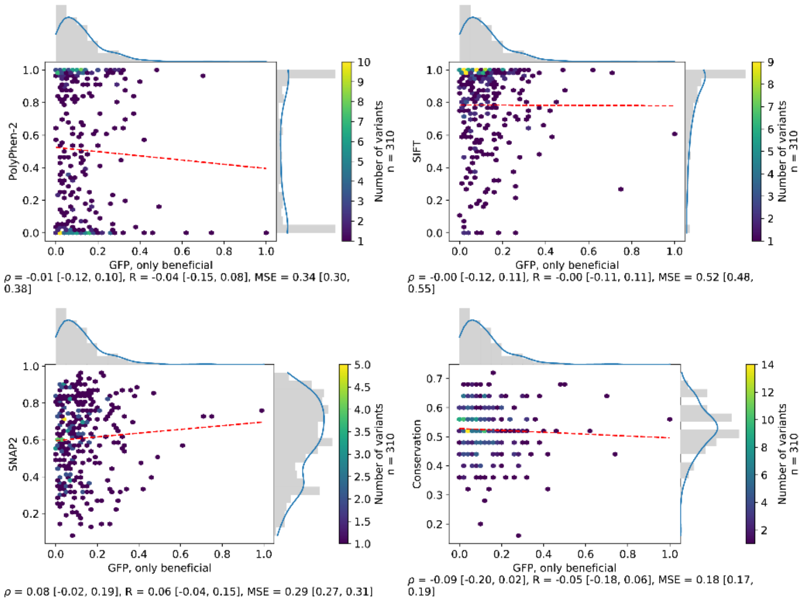
PolyPhen-2
GFP, only beneficial
Number of variants
n = 310
ρ = -0.01 [-0.12, 0.10], R = -0.04 [-0.15, 0.08], MSE = 0.34 [0.30, 0.38]
SIFT
ρ = -0.00 [-0.12, 0.11], R = -0.00 [-0.11, 0.11], MSE = 0.52 [0.48, 0.55]
SNAP2
ρ = 0.08 [-0.02, 0.19], R = 0.06 [-0.04, 0.15], MSE = 0.29 [0.27, 0.31]
Conservation
ρ = -0.09 [-0.20, 0.02], R = -0.05 [-0.18, 0.06], MSE = 0.18 [0.17, 0.19]

# S5q

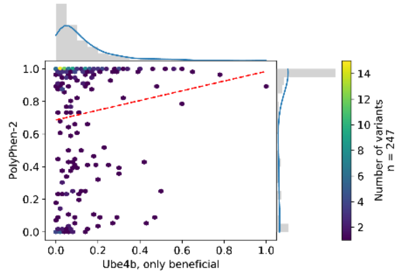

ρ = 0.13 [0.01, 0.25], R = 0.12 [0.01, 0.22], MSE = 0.50 [0.46, 0.55]

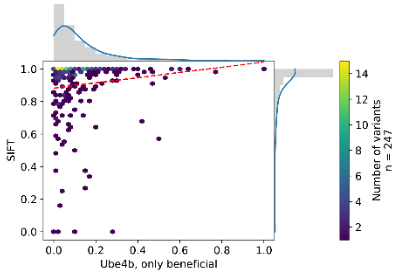

ρ = 0.20 [0.09, 0.32], R = 0.12 [0.01, 0.21], MSE = 0.65 [0.62, 0.69]

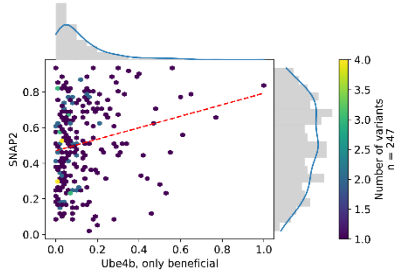

ρ = 0.22 [0.09, 0.34], R = 0.20 [0.09, 0.32], MSE = 0.21 [0.19, 0.23]

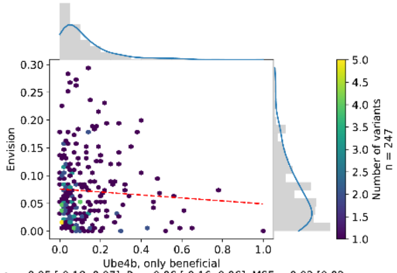

ρ = -0.05 [-0.18, 0.07], R = -0.06 [-0.16, 0.06], MSE = 0.03 [0.02, 0.04]

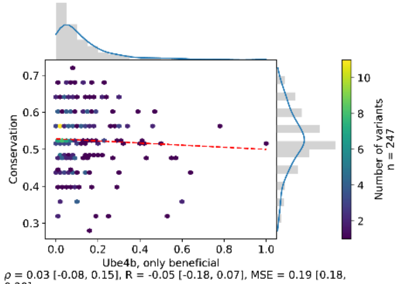

ρ = 0.03 [-0.08, 0.15], R = -0.05 [-0.18, 0.07], MSE = 0.19 [0.18, 0.20]

# S5r

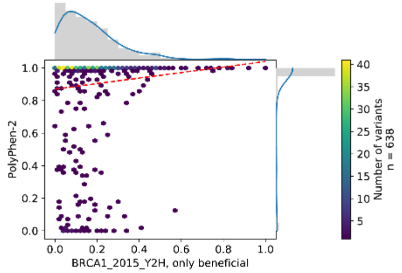

ρ = 0.11 [0.03, 0.19], R = 0.10 [0.04, 0.15], MSE = 0.61 [0.59, 0.63]

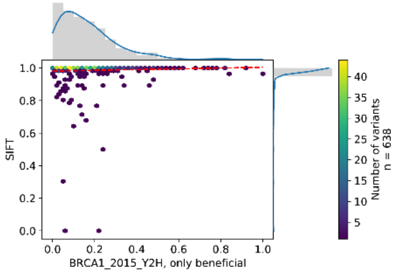

ρ = 0.09 [0.01, 0.17], R = 0.05 [0.00, 0.10], MSE = 0.69 [0.67, 0.70]

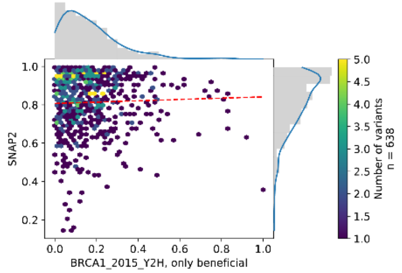

ρ = 0.07 [-0.01, 0.14], R = 0.03 [-0.05, 0.11], MSE = 0.45 [0.44, 0.47]

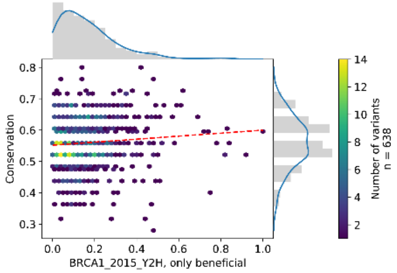

ρ = 0.10 [0.02, 0.18], R = 0.09 [0.01, 0.17], MSE = 0.18 [0.17, 0.19]

## S5s

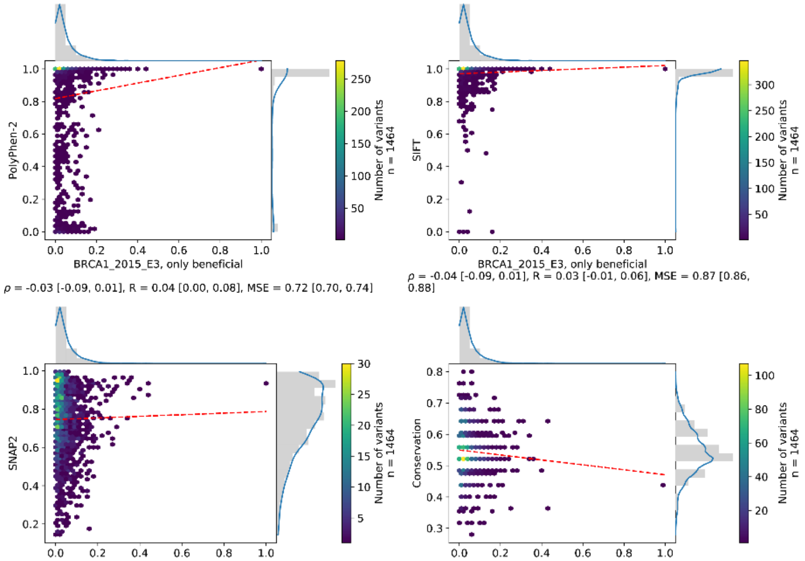

$\rho$ = -0.03 [-0.09, 0.01], R = 0.04 [0.00, 0.08], MSE = 0.72 [0.70, 0.74]

$\rho$ = -0.04 [-0.09, 0.01], R = 0.03 [-0.01, 0.06], MSE = 0.87 [0.86, 0.88]

$\rho$ = -0.06 [-0.11, -0.00], R = 0.01 [-0.04, 0.06], MSE = 0.52 [0.51, 0.54]

$\rho$ = -0.04 [-0.09, 0.01], R = -0.06 [-0.11, -0.01], MSE = 0.26 [0.25, 0.26]

## S5t

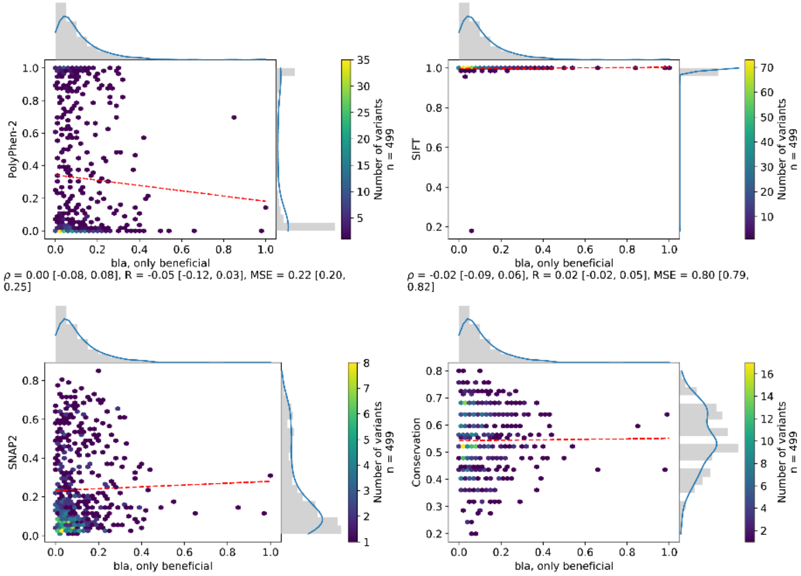

$\rho$ = 0.00 [-0.08, 0.08], R = -0.05 [-0.12, 0.03], MSE = 0.22 [0.20, 0.25]

$\rho$ = -0.02 [-0.09, 0.06], R = 0.02 [-0.02, 0.05], MSE = 0.80 [0.79, 0.82]

$\rho$ = 0.07 [-0.01, 0.15], R = 0.03 [-0.04, 0.10], MSE = 0.07 [0.06, 0.08]

$\rho$ = 0.00 [-0.08, 0.09], R = 0.01 [-0.07, 0.08], MSE = 0.21 [0.20, 0.23]

# S5u

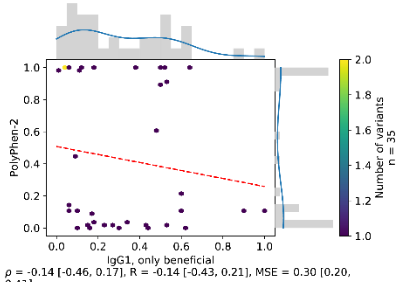

ρ = -0.14 [-0.46, 0.17], R = -0.14 [-0.43, 0.21], MSE = 0.30 [0.20, 0.41]

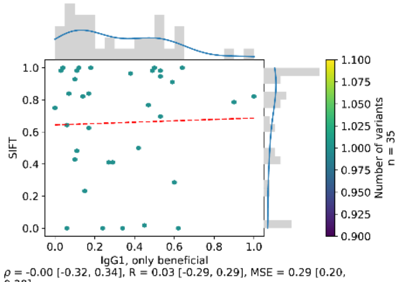

ρ = -0.00 [-0.32, 0.34], R = 0.03 [-0.29, 0.29], MSE = 0.29 [0.20, 0.38]

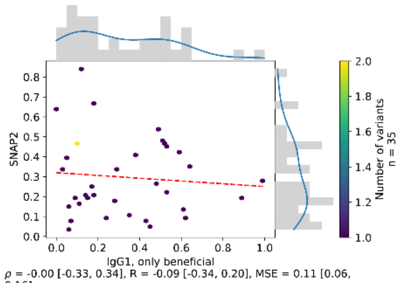

ρ = -0.00 [-0.33, 0.34], R = -0.09 [-0.34, 0.20], MSE = 0.11 [0.06, 0.16]

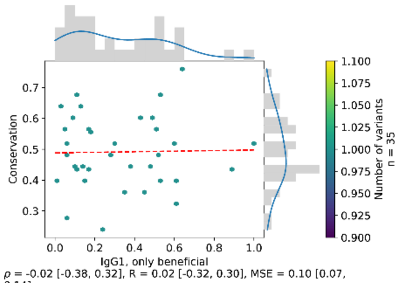

ρ = -0.02 [-0.38, 0.32], R = 0.02 [-0.32, 0.30], MSE = 0.10 [0.07, 0.14]

#### Figure S6: Agreement between independently measured deleterious effect scores.

SAVs from ten DMS experiments on four proteins are taken into account. Hexbin plots show the correlation between two scores from independent experiments on the same protein (see Table S1). Values on axes range from 0 (neutral) to 1 (maximal effect). Dashed red lines give linear least-squared regressions. Marginals denote distributions of experimental scores with a kernel density estimation overlaid in blue. The footer denotes Spearman ρ, Pearson R and the mean squared error.

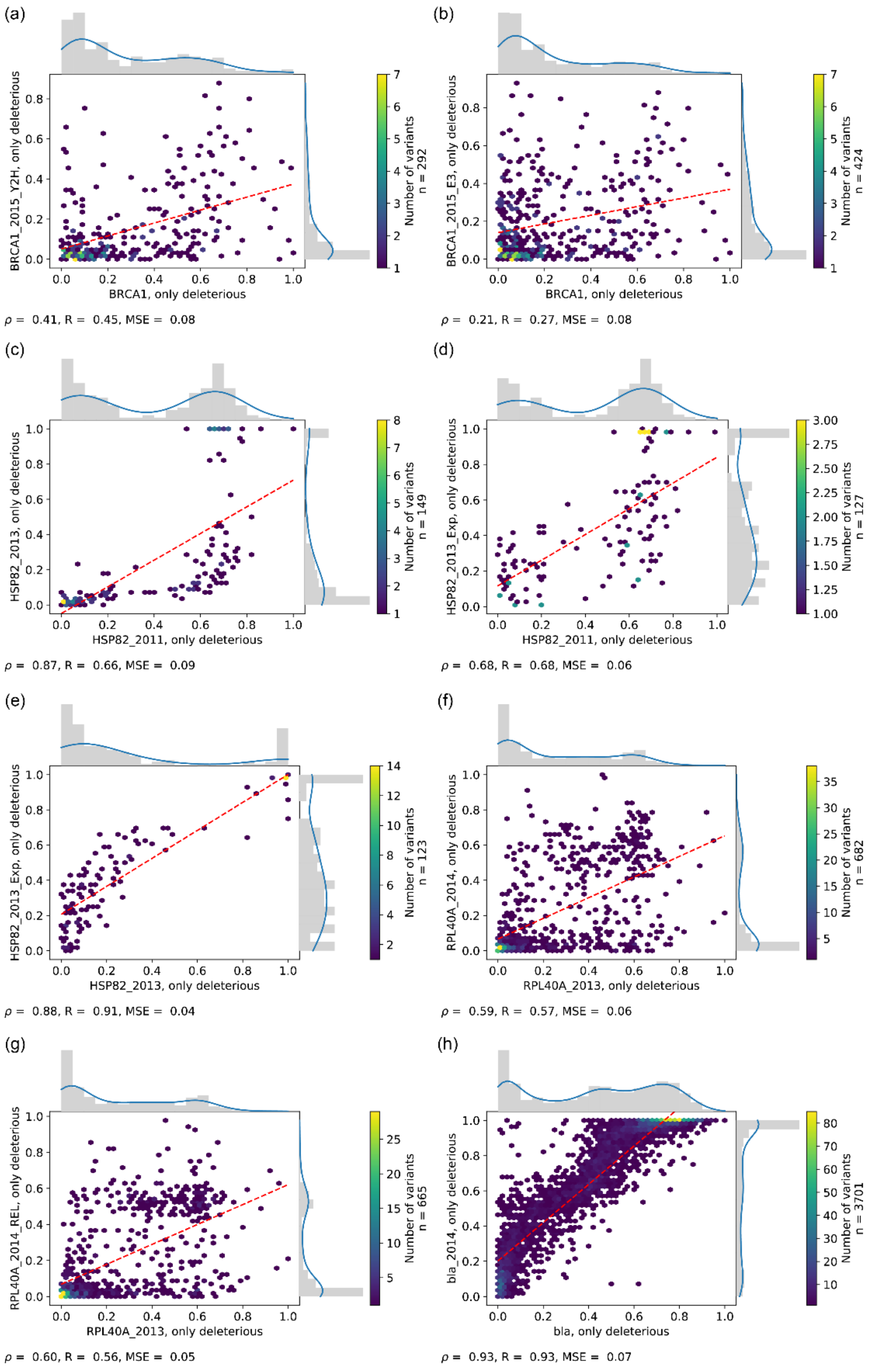
(a)
BRCA1_2015_Y2H, only deleterious
BRCA1, only deleterious
Number of variants
n = 292
ρ = 0.41, R = 0.45, MSE = 0.08
(b)
BRCA1_2015_E3, only deleterious
BRCA1, only deleterious
Number of variants
n = 424
ρ = 0.21, R = 0.27, MSE = 0.08
(c)
HSP82_2013, only deleterious
HSP82_2011, only deleterious
Number of variants
n = 149
ρ = 0.87, R = 0.66, MSE = 0.09
(d)
HSP82_2013_Exp, only deleterious
HSP82_2011, only deleterious
Number of variants
n = 127
ρ = 0.68, R = 0.68, MSE = 0.06
(e)
HSP82_2013_Exp, only deleterious
HSP82_2013, only deleterious
Number of variants
n = 123
ρ = 0.88, R = 0.91, MSE = 0.04
(f)
RPL40A_2014, only deleterious
RPL40A_2013, only deleterious
Number of variants
n = 682
ρ = 0.59, R = 0.57, MSE = 0.06
(g)
RPL40A_2014_REL, only deleterious
RPL40A_2013, only deleterious
Number of variants
n = 665
ρ = 0.60, R = 0.56, MSE = 0.05
(h)
bla_2014, only deleterious
bla, only deleterious
Number of variants
n = 3701
ρ = 0.93, R = 0.93, MSE = 0.07

#### Figure S7: Agreement between independently measured beneficial effect scores.

SAVs from ten DMS experiments on four proteins are taken into account. Hexbin plots show the correlation between two SAV effect scores from independent experiments on the same protein (see Table S1). Values on axes range from 0 (neutral) to 1 (maximal effect). Dashed red lines give linear least-squared regressions. Marginals denote distributions of experimental scores with a kernel density estimation overlaid in blue. The footer denotes Spearman ρ, Pearson R and the mean squared error.

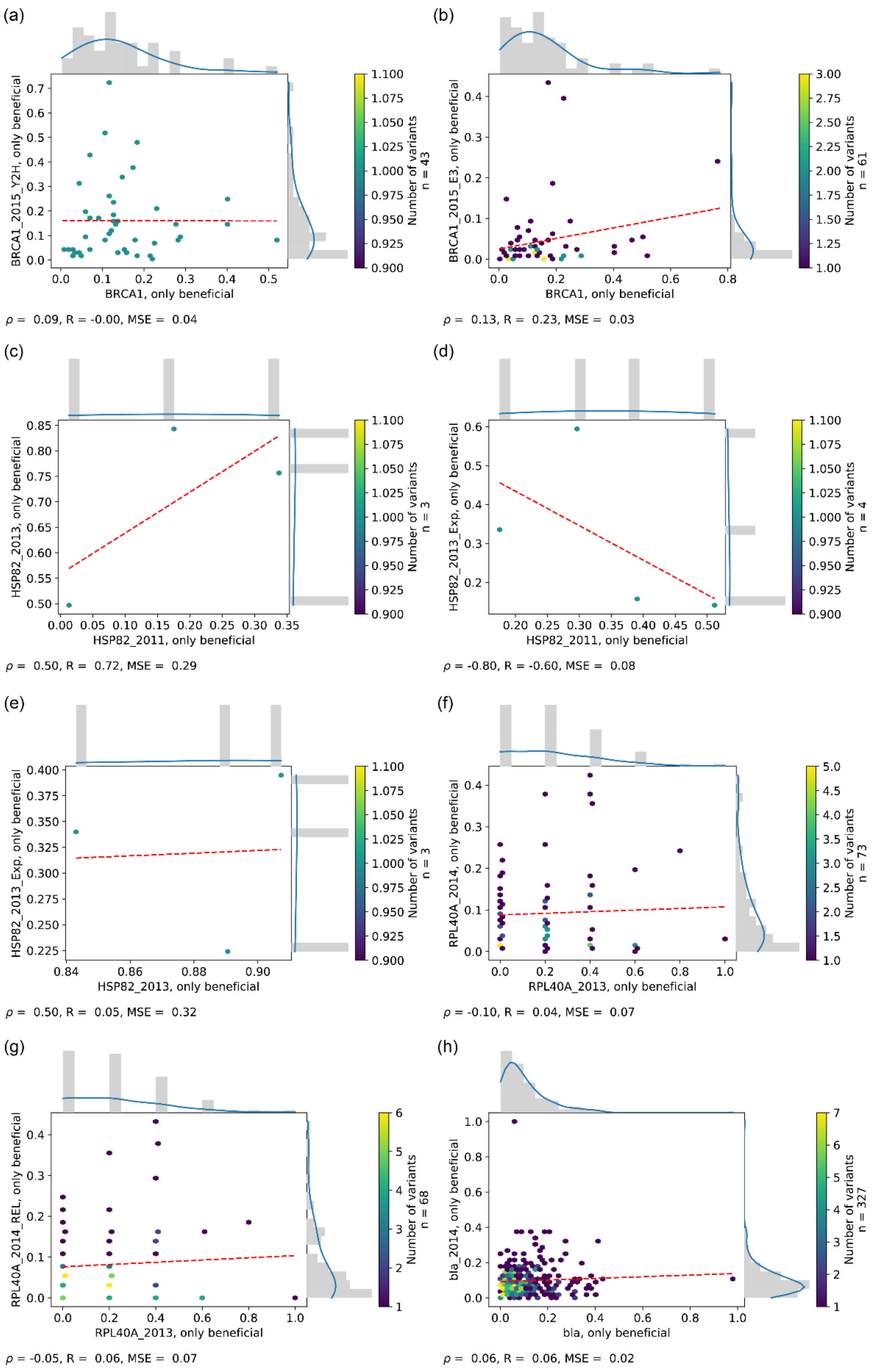
(a)
BRCA1_2015_Y2H, only beneficial
BRCA1, only beneficial
Number of variants n = 43
ρ = 0.09, R = -0.00, MSE = 0.04
(b)
BRCA1_2015_E3, only beneficial
BRCA1, only beneficial
Number of variants n = 61
ρ = 0.13, R = 0.23, MSE = 0.03
(c)
HSP82_2013, only beneficial
HSP82_2011, only beneficial
Number of variants n = 3
ρ = 0.50, R = 0.72, MSE = 0.29
(d)
HSP82_2013_Exp, only beneficial
HSP82_2011, only beneficial
Number of variants n = 4
ρ = -0.80, R = -0.60, MSE = 0.08
(e)
HSP82_2013_Exp, only beneficial
HSP82_2013, only beneficial
Number of variants n = 3
ρ = 0.50, R = 0.05, MSE = 0.32
(f)
RPL40A_2014, only beneficial
RPL40A_2013, only beneficial
Number of variants n = 73
ρ = -0.10, R = 0.04, MSE = 0.07
(g)
RPL40A_2014_REL, only beneficial
RPL40A_2013, only beneficial
Number of variants n = 68
ρ = -0.05, R = 0.06, MSE = 0.07
(h)
bla_2014, only beneficial
bla, only beneficial
Number of variants n = 327
ρ = 0.06, R = 0.06, MSE = 0.02

**Figure S8: Number of neutral and deleterious effect SAVs (syn95).**

Syn95 denotes the thresholding scheme in which SAVs with an effect in the range of 95% of synonymous variants' effect scores are considered neutral. All outside of this range are considered effect (see Methods). ROC curves and AUCs for this thresholding scheme are shown in Fig. S10. SetCommonSyn95, for which every method could perform predictions is a subset of SAVs from BRCA1, TPMT, PTEN and PPARG. LGK does not contain deleterious effect variants using syn90 or syn95, but does using syn99.

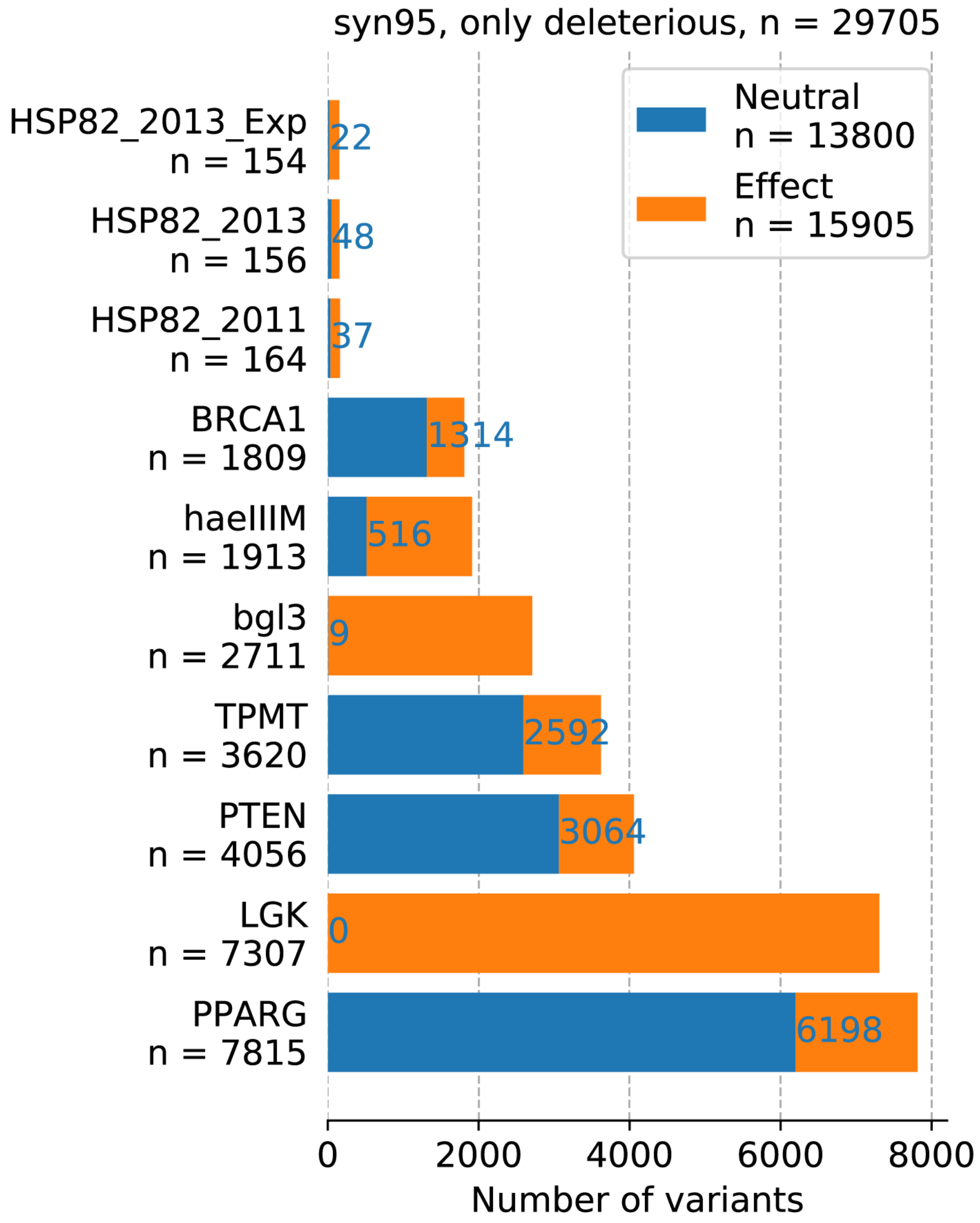

**Figure S9. Precision-Recall curves for classifying deleterious effect SAVs (syn95).**

SAVs are classified as in Figure 1. For SNAP2 the cross-over point is at precision = recall = 0.49, while for Conservation it is at precision = 0.47 and recall = 0.5 For PolyPhen-2 and SIFT, curves never cross, before reaching the point of precision = 1, recall = 0.

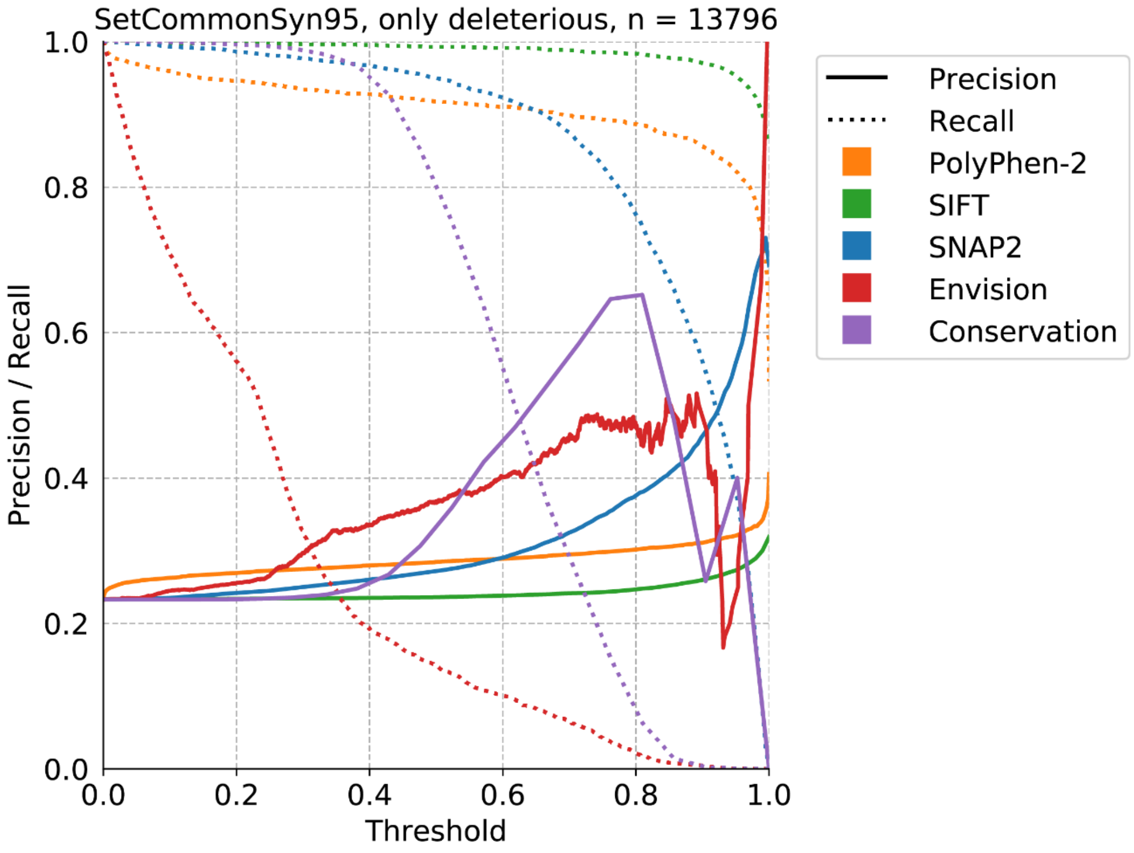

**Figure S10: ROC curves for classifying deleterious effect SAVs (syn95).**

SAVs are classified into either neutral, defined by the middle 95% of synonymous variants' scores, or effect (syn95, see Methods). In every plot the number of SAVs (denoted in the title) is the largest common subset of SAVs for which a prediction is available from every method. Missing methods did not perform any predictions. Shaded areas around lines denote 95% confidence intervals. The legend denotes the AUC for every method together with the 95% confidence intervals. Horizontal dashed lines denote the default score threshold used by SNAP2 (blue) and SIFT (green). Panel (j) shows the combined set of SAVs from four DMS experiments for which every method performed a prediction (SetCommonSyn95).

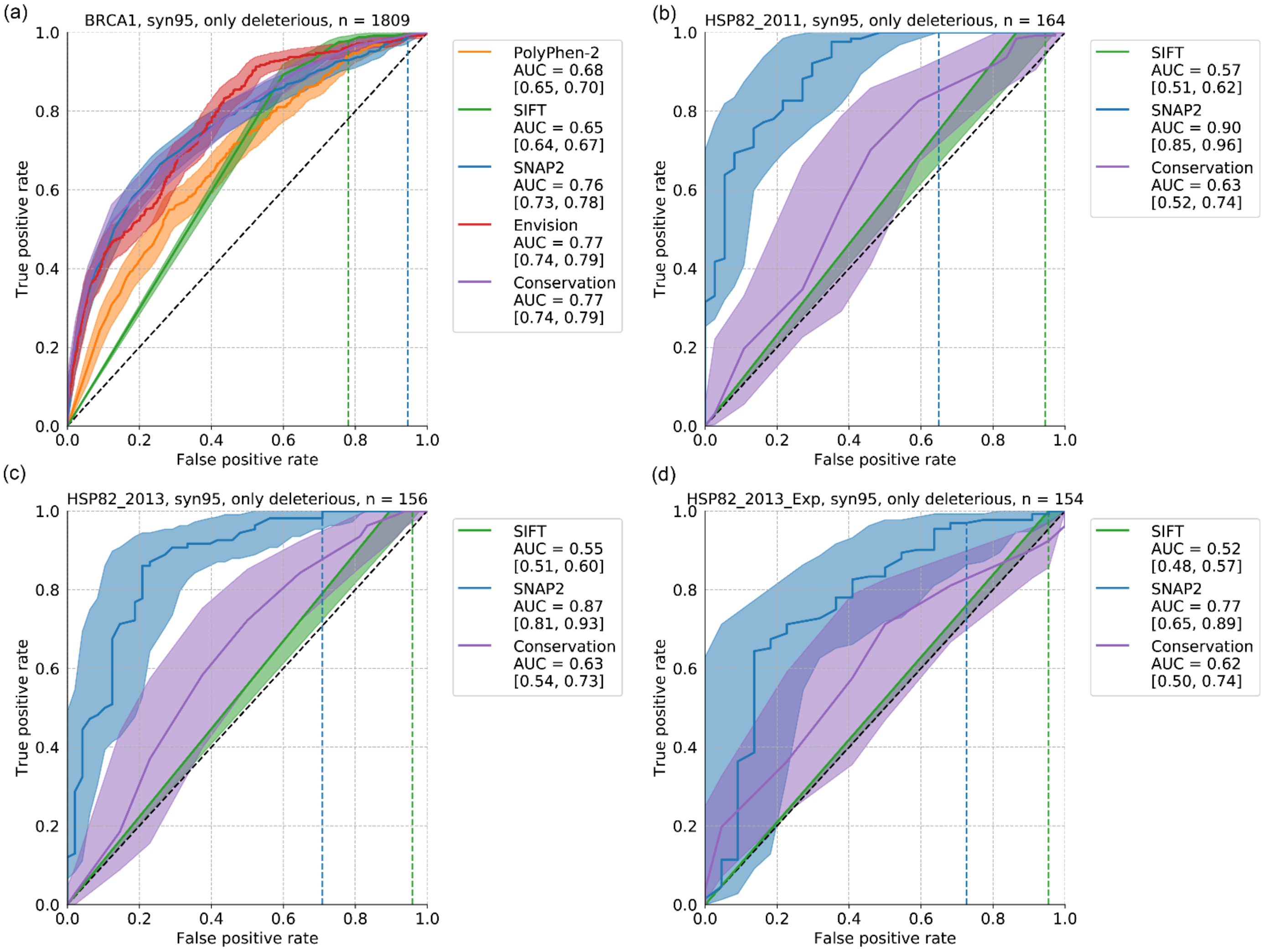

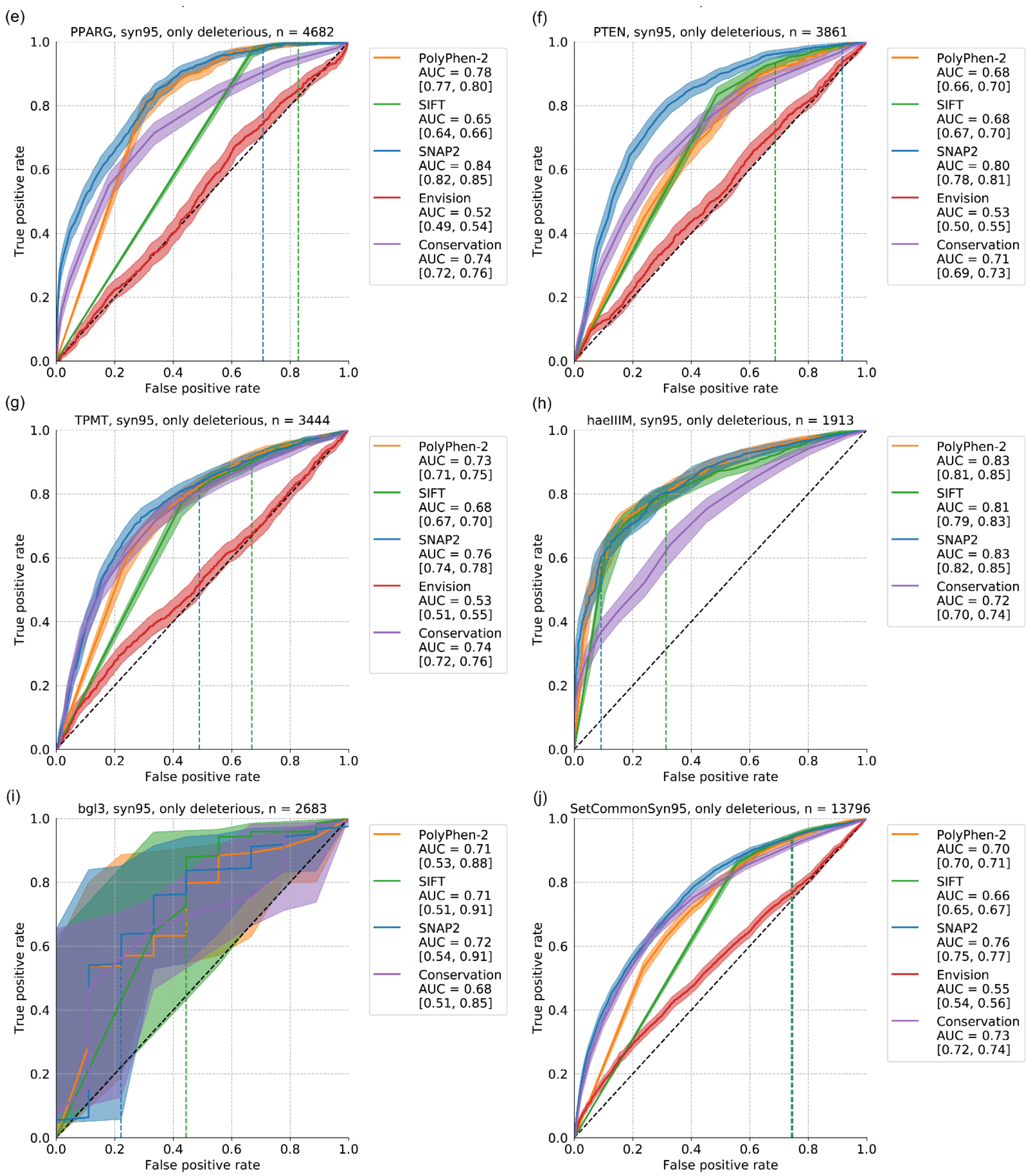
(e)
PPARG, syn95, only deleterious, n = 4682
True positive rate
False positive rate
PolyPhen-2
AUC = 0.78
[0.77, 0.80]
SIFT
AUC = 0.65
[0.64, 0.66]
SNAP2
AUC = 0.84
[0.82, 0.85]
Envision
AUC = 0.52
[0.49, 0.54]
Conservation
AUC = 0.74
[0.72, 0.76]
(f)
PTEN, syn95, only deleterious, n = 3861
True positive rate
False positive rate
PolyPhen-2
AUC = 0.68
[0.66, 0.70]
SIFT
AUC = 0.68
[0.67, 0.70]
SNAP2
AUC = 0.80
[0.78, 0.81]
Envision
AUC = 0.53
[0.50, 0.55]
Conservation
AUC = 0.71
[0.69, 0.73]
(g)
TPMT, syn95, only deleterious, n = 3444
True positive rate
False positive rate
PolyPhen-2
AUC = 0.73
[0.71, 0.75]
SIFT
AUC = 0.68
[0.67, 0.70]
SNAP2
AUC = 0.76
[0.74, 0.78]
Envision
AUC = 0.53
[0.51, 0.55]
Conservation
AUC = 0.74
[0.72, 0.76]
(h)
haeIIIM, syn95, only deleterious, n = 1913
True positive rate
False positive rate
PolyPhen-2
AUC = 0.83
[0.81, 0.85]
SIFT
AUC = 0.81
[0.79, 0.83]
SNAP2
AUC = 0.83
[0.82, 0.85]
Conservation
AUC = 0.72
[0.70, 0.74]
(i)
bgl3, syn95, only deleterious, n = 2683
True positive rate
False positive rate
PolyPhen-2
AUC = 0.71
[0.53, 0.88]
SIFT
AUC = 0.71
[0.51, 0.91]
SNAP2
AUC = 0.72
[0.54, 0.91]
Conservation
AUC = 0.68
[0.51, 0.85]
(j)
SetCommonSyn95, only deleterious, n = 13796
True positive rate
False positive rate
PolyPhen-2
AUC = 0.70
[0.70, 0.71]
SIFT
AUC = 0.66
[0.65, 0.67]
SNAP2
AUC = 0.76
[0.75, 0.77]
Envision
AUC = 0.55
[0.54, 0.56]
Conservation
AUC = 0.73
[0.72, 0.74]

**Figure S11: ROC curves for classifying deleterious and beneficial effect SAVs (syn90, syn99).**

SAVs are classified into either neutral, defined by the middle 90% or 99% of synonymous variants' scores, or effect. SetCommonSyn90 ((a), (c)) and SetCommonSyn99 ((b), (d)) are the respectively classified SAVs from all four DMS experiments for which each method performed a prediction (see Methods). Shaded areas around lines denote 95% confidence intervals. The legend denotes the AUC for every method together with the 95% confidence intervals. Horizontal dashed lines denote the default score threshold used by SNAP2 (blue) and SIFT (green).

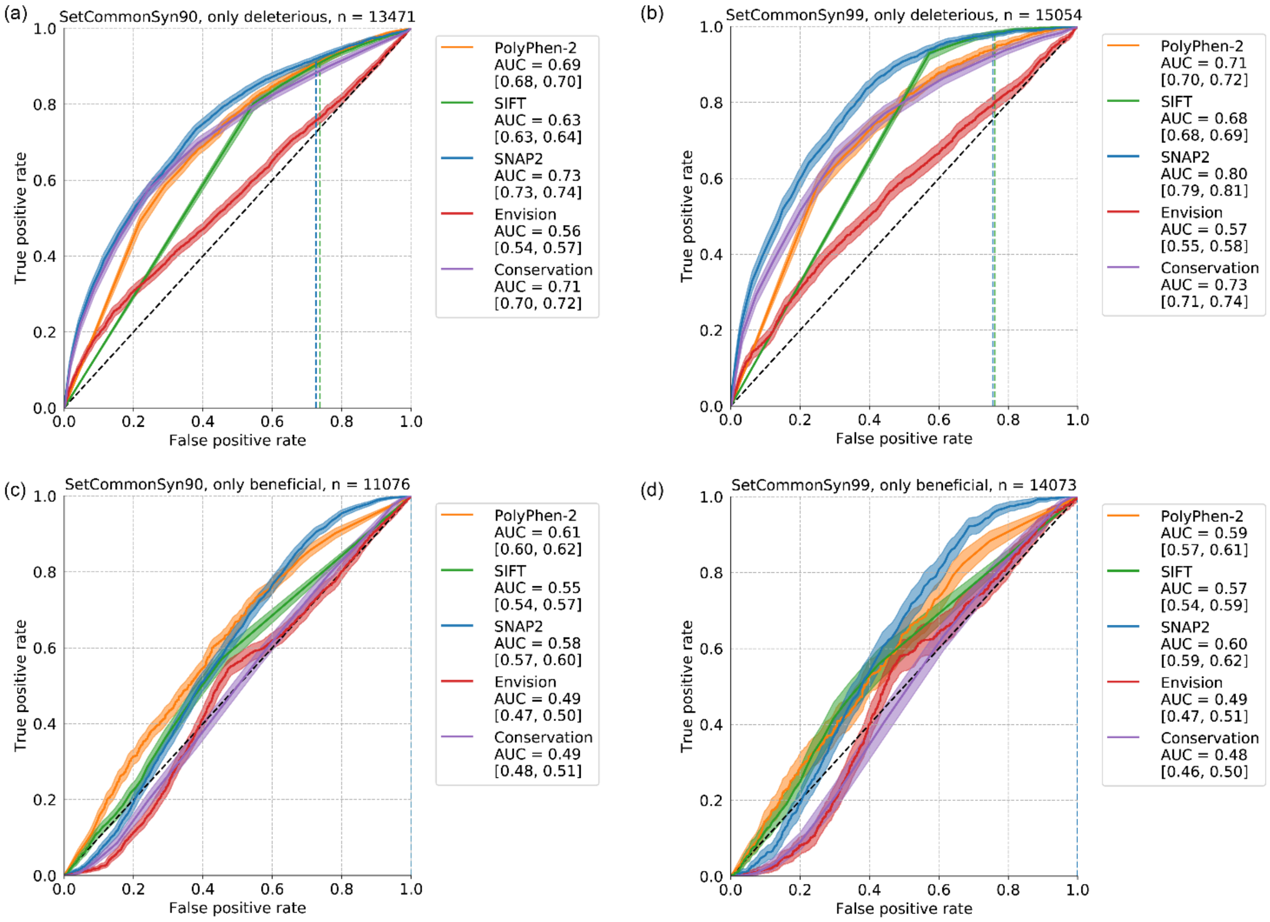

#### Table S1: DMS experiments used throughout this work.

The first 22 entries denote SetAll, below those are other measurements used for additional analyses (see Methods). Dataset identifier denotes the name used for the set of variant scores throughout the manuscript. # variants denotes the count of non-synonymous variants in the dataset, and number of synonymous in parentheses. Coverage is the percentage of residues in the reference sequence (see Table S12) for which at least one SAV is scored. Compl. refers to the percentage of all possible SAVs scored within the region of the protein for which scores are provided. Synonymous mutations are not considered for Coverage and Completeness.

| Dataset identifier | Protein | # variants | Coverage | Compl. | Source org. | Reference |
|---|---|---|---|---|---|---|
| ccdB | Toxin CcdB | 1208 | 99 | 63 | *E. coli* | [1] |
| YAP1 | Transcriptional coactivator YAP1 | 363 | 7 | 4 | *H. sapiens* | [2] |
| MAPK1 | Mitogen-activated protein kinase 1 | 6810 | 100 | 100 | *H. sapiens* | [3] |
| BRCA1 | Breast cancer type 1 susceptibility protein | 1837 (295) | 17 | 5 | *H. sapiens* | [4] |
| CCR5 | C-C chemokine receptor type 5 | 4668 | 100 | 70 | *H. sapiens* | [5] |
| CXCR4 | C-X-C chemokine receptor type 4 | 3877 | 99 | 58 | *H. sapiens* | [5] |
| HSP82_2011 | ATP-dependent molecular chaperone HSP82 | 171 (7) | 1 | 1 | *S. cerevisiae* | [6] |
| HSP82_2013 | ATP-dependent molecular chaperone HSP82 | 170 (8) | 1 | 1 | *S. cerevisiae* | [7] |
| HSP82_2013_Exp | ATP-dependent molecular chaperone HSP82 | 171 (9) | 1 | 1 | *S. cerevisiae* | [8] |
| GAL4 | Regulatory protein GAL4 | 1196 | 7 | 7 | *S. cerevisiae* | [9] |
| LGK | Levoglucosan kinase | 7629 (439) | 100 | 91 | *L. starkeyi* | [10] |
| PPARG | Peroxisome proliferator-activated receptor γ | 9595 (505) | 100 | 100 | *H. sapiens* | [11] |
| PTEN | Phosphatase and tensin homolog | 4112 (156) | 89 | 54 | *H. sapiens* | [12] |
| TPMT | Thiopurine S-methyltransferase | 3689 (139) | 98 | 79 | *H. sapiens* | [12] |
| haeIIIM | Modification methylase HaeIII | 1957 (321) | 100 | 31 | *H. aegyptius* | [13] |
| bgl3 | β-glucosidase | 2857 (479) | 100 | 31 | *Streptomyces* | [14] |
| GFP | Green fluorescent protein | 1114 | 98 | 25 | *Ae. victoria* | [15] |
| Ube4b | Ubiquitin conjugation factor E4 B | 899 | 9 | 4 | *M. musculus* | [16] |
| BRCA1_2015_Y2H | Breast cancer type 1 susceptibility protein | 1748 | 5 | 5 | *H. sapiens* | [17] |
| BRCA1_2015_E3 | Breast cancer type 1 susceptibility protein | 6620 | 16 | 19 | *H. sapiens* | [17] |
| Bla | β-lactamase TEM | 4997 | 92 | 92 | *E. coli* | [18] |
| IgG1 | Immunoglobulin gamma-1 heavy chain | 401 | 24 | 5 | *H. sapiens* | [19] |
| | **Sum** | **66089 (2358)** | | | | |
| RPL40A_2013 | Ubiquitin | 1195 (75) | 59 | 49 | *S. cerevisiae* | [20] |
| RPL40A_2014 | Ubiquitin | 1365 (71) | 59 | 56 | *S. cerevisiae* | [21] |
| RPL40A_2014_REL | Ubiquitin | 1365 (71) | 59 | 56 | *S. cerevisiae* | [21] |
| bla_2014 | β-lactamase TEM | 5160 (269) | 99 | 95 | *E. coli* | [22] |

**Table S2: Beneficial and deleterious variants at the same residue in SetAll.**

| Dataset | Residues with benefical and deleterious variants |
|---|---|
| ccdB | 0% |
| YAP1 | 58% |
| MAPK1 | 80% |
| BRCA1 | 80% |
| CCR5 | 94% |
| CXCR4 | 91% |
| HSP82_2011 | 89% |
| HSP82_2013 | 56% |
| HSP82_2013_Exp | 100% |
| GAL4 | 59% |
| LGK | 28% |
| PPARG | 35% |
| PTEN | 70% |
| TPMT | 88% |
| haeIIIM | 32% |
| bgl3 | 100% |
| GFP | 60% |
| Ube4b | 75% |
| BRCA1_2015_Y2H | 94% |
| BRCA1_2015_E3 | 90% |
| bla | 48% |
| IgG1 | 25% |
| RPL40A_2013 | 64% |
| RPL40A_2014 | 91% |
| RPL40A_2014_REL | 88% |
| bla_2014 | 84% |

**Table S3: p-values for the difference between Spearman ρ on SetCommon.**

Statistical tests for the difference of correlation measures between each pair of prediction methods (TwoDcorR) were performed in R using the WRS package [23, 24]. Alpha = 0.05, bootstrap iterations = 500. P-values given as 0 were too small for reporting in R. Generally, the value of p-values for such large datasets should be questioned and more emphasis be put on the confidence intervals themselves as reported in Tables S6, S7. We report them here to show the clear trends and since some people might expect to see them. SetCommon refers to the set of SAVs for which a prediction is available from every method.

| Method 1 | Method 2 | SetCommon, only deleterious n = 17,781 SAVs | SetCommon, only beneficial n = 15,200 SAVs |
|---|---|---|---|
| PolyPhen-2 | SIFT | 0 | 0 |
| PolyPhen-2 | SNAP2 | 0 | 0 |
| PolyPhen-2 | Envision | 0 | 0 |
| PolyPhen-2 | Conservation | 0 | 0.296 |
| SIFT | SNAP2 | 0 | 0 |
| SIFT | Envision | 0 | 0.604 |
| SIFT | Conservation | 0 | 0 |
| SNAP2 | Envision | 0 | 0 |
| SNAP2 | Conservation | 0 | 0 |
| Envision | Conservation | 0 | 0 |

##### Table S4: Change in performance measures when limiting to subsets of high effect SAVs

SetCommon, containing all 17,781 deleterious and 15,200 beneficial effect SAVs for which every method performs predictions is limited to the subsets with only the 75%, 50% or 25% highest effect variants. The difference in Spearman ρ and mean squared error (MSE) between SetCommon (see Tables [S10LD], S6, S7, S8) and the respective subset is shown. For ρ negative values denote a decrease in performance, for MSE positive values indicated a larger error and hence decreased performance.

| **Change** in | Subset | PolyPhen-2 | SIFT | SNAP2 | Envision | Conservation |
|---|---|---|---|---|---|---|
| Spearman ρ | SetDelStrongest75 | -0.07 | -0.02 | -0.03 | -0.02 | -0.06 |
| | SetDelStrongest50 | -0.2 | -0.09 | -0.12 | -0.13 | -0.17 |
| | SetDelStrongest25 | -0.33 | -0.18 | -0.29 | -0.3 | -0.27 |
| MSE | SetDelStrongest75 | 0.02 | -0.01 | 0.02 | 0.01 | 0.01 |
| | SetDelStrongest50 | 0.05 | -0.01 | 0.05 | 0.02 | 0.02 |
| | SetDelStrongest25 | 0.1 | 0.03 | 0.12 | 0.01 | 0.05 |
| Spearman ρ | SetBenStrongest75 | 0.03 | 0.01 | 0.04 | -0.01 | 0.01 |
| | SetBenStrongest50 | 0.06 | 0.01 | 0.07 | 0.01 | 0 |
| | SetBenStrongest25 | 0.09 | 0.02 | 0.1 | 0.04 | -0.02 |
| MSE | SetBenStrongest75 | 0 | 0.02 | 0.01 | -0.01 | 0.01 |
| | SetBenStrongest50 | -0.01 | 0.03 | 0.01 | -0.01 | 0.02 |
| | SetBenStrongest25 | 0 | 0.05 | 0.02 | -0.01 | 0.04 |

**Table S5: Mean squared error between five variant effect prediction methods and experimentally measured deleterious effect scores from 22 DMS experiments in SetAll.**

If a method did not perform predictions for a given set, cells are left empty. Values in parentheses denote 95% confidence intervals. SetCommon contains 17,781 deleterious effect SAVs from ten proteins for which every method performed predictions (see Tables 1, 2 and Figure 1).

| | Envision | Conservation | PolyPhen-2 | SIFT | SNAP2 |
|---|---|---|---|---|---|
| ccdB | | 0.09 [0.08, 0.11] | 0.22 [0.19, 0.25] | 0.19 [0.17, 0.21] | 0.14 [0.13, 0.16] |
| YAP1 | 0.28 [0.24, 0.31] | 0.09 [0.08, 0.10] | 0.28 [0.24, 0.31] | 0.3 [0.27, 0.34] | 0.09 [0.08, 0.11] |
| MAPK1 | 0.03 [0.03, 0.04] | 0.25 [0.25, 0.26] | 0.56 [0.55, 0.57] | 0.64 [0.64, 0.65] | 0.36 [0.35, 0.37] |
| BRCA1 | 0.05 [0.04, 0.05] | 0.15 [0.14, 0.15] | 0.34 [0.32, 0.36] | 0.59 [0.58, 0.61] | 0.44 [0.43, 0.45] |
| CCR5 | 0.05 [0.05, 0.06] | 0.27 [0.26, 0.27] | 0.5 [0.48, 0.51] | 0.69 [0.68, 0.71] | 0.34 [0.33, 0.35] |
| CXCR4 | 0.03 [0.03, 0.04] | 0.24 [0.23, 0.24] | 0.47 [0.45, 0.48] | 0.67 [0.66, 0.68] | 0.26 [0.25, 0.26] |
| HSP82_2011 | | 0.13 [0.11, 0.16] | | 0.48 [0.42, 0.53] | 0.2 [0.17, 0.23] |
| HSP82_2013 | | 0.23 [0.21, 0.25] | | 0.68 [0.62, 0.73] | 0.36 [0.33, 0.39] |
| HSP82_2013_Exp | | 0.12 [0.10, 0.14] | | 0.42 [0.37, 0.47] | 0.2 [0.17, 0.23] |
| GAL4 | 0.15 [0.14, 0.16] | 0.13 [0.12, 0.14] | 0.35 [0.33, 0.37] | 0.4 [0.38, 0.42] | 0.21 [0.20, 0.22] |
| LGK | | 0.17 [0.16, 0.17] | 0.37 [0.37, 0.38] | 0.48 [0.48, 0.48] | 0.14 [0.14, 0.15] |
| PPARG | 0.09 [0.08, 0.09] | 0.14 [0.14, 0.15] | 0.48 [0.47, 0.50] | 0.55 [0.54, 0.56] | 0.31 [0.30, 0.33] |
| PTEN | 0.08 [0.07, 0.08] | 0.13 [0.12, 0.13] | 0.37 [0.36, 0.38] | 0.47 [0.46, 0.48] | 0.32 [0.31, 0.32] |
| TPMT | 0.06 [0.06, 0.06] | 0.22 [0.21, 0.22] | 0.45 [0.44, 0.46] | 0.56 [0.55, 0.57] | 0.21 [0.20, 0.22] |
| haeIIIM | | 0.1 [0.10, 0.10] | 0.14 [0.13, 0.15] | 0.19 [0.18, 0.20] | 0.1 [0.09, 0.11] |
| bgl3 | | 0.16 [0.16, 0.17] | 0.24 [0.23, 0.25] | 0.49 [0.48, 0.50] | 0.13 [0.13, 0.14] |
| GFP | | 0.23 [0.23, 0.24] | 0.49 [0.47, 0.52] | 0.62 [0.60, 0.64] | 0.37 [0.35, 0.38] |
| Ube4b | 0.02 [0.02, 0.02] | 0.21 [0.20, 0.22] | 0.58 [0.55, 0.60] | 0.71 [0.69, 0.73] | 0.3 [0.28, 0.31] |
| BRCA1_2015_Y2H | | 0.21 [0.20, 0.22] | 0.65 [0.64, 0.67] | 0.71 [0.70, 0.73] | 0.49 [0.47, 0.50] |
| BRCA1_2015_E3 | | 0.14 [0.14, 0.15] | 0.45 [0.44, 0.46] | 0.53 [0.53, 0.54] | 0.29 [0.29, 0.30] |
| bla | | 0.14 [0.14, 0.15] | 0.27 [0.26, 0.28] | 0.48 [0.47, 0.49] | 0.1 [0.10, 0.11] |
| IgG1 | | 0.13 [0.12, 0.14] | 0.12 [0.10, 0.15] | 0.05 [0.04, 0.07] | 0.21 [0.19, 0.23] |
| SetCommon | 0.06 [0.06, 0.07] | 0.19 [0.19, 0.19] | 0.45 [0.45, 0.46] | 0.58 [0.57, 0.58] | 0.3 [0.30, 0.30] |

**Table S6: Spearman ρ between five variant effect prediction methods and experimentally measured deleterious effect scores from 22 DMS experiments in SetAll.**

If a method did not perform predictions for a given set, cells are left empty. Values in parentheses denote 95% confidence intervals. nan values are caused by lack of diversity in prediction scores, making calculation of ρ impossible. SetCommon contains 17,781 deleterious effect SAVs from ten proteins for which every method performed predictions (see Tables 1, 2 and Figure 1).

| | Envision | Conservation | PolyPhen-2 | SIFT | SNAP2 |
|---|---|---|---|---|---|
| ccdB | | -0.11 [-0.20, -0.00] | -0.11 [-0.21, -0.01] | -0.26 [-0.35, -0.17] | -0.18 [-0.27, -0.08] |
| YAP1 | 0.32 [0.19, 0.43] | 0.22 [0.10, 0.34] | 0.17 [0.05, 0.29] | 0.29 [0.17, 0.39] | 0.51 [0.41, 0.60] |
| MAPK1 | 0.39 [0.35, 0.42] | 0.41 [0.37, 0.44] | 0.35 [0.32, 0.38] | 0.15 [0.12, 0.19] | 0.46 [0.43, 0.49] |
| BRCA1 | 0.42 [0.37, 0.45] | 0.44 [0.40, 0.48] | 0.26 [0.21, 0.31] | 0.33 [0.28, 0.37] | 0.39 [0.34, 0.43] |
| CCR5 | 0.11 [0.06, 0.16] | 0.38 [0.34, 0.43] | 0.26 [0.22, 0.31] | 0.19 [0.14, 0.23] | 0.29 [0.25, 0.33] |
| CXCR4 | 0.09 [0.05, 0.13] | 0.29 [0.25, 0.33] | 0.2 [0.16, 0.24] | 0.17 [0.13, 0.21] | 0.28 [0.24, 0.32] |
| HSP82_2011 | | 0.25 [0.08, 0.39] | | 0.27 [nan, nan] | 0.69 [0.59, 0.77] |
| HSP82_2013 | | 0.28 [0.12, 0.42] | | 0.22 [nan, nan] | 0.61 [0.50, 0.70] |
| HSP82_2013_Exp | | 0.19 [0.02, 0.37] | | [nan, nan] | 0.52 [0.38, 0.63] |
| GAL4 | 0.37 [0.31, 0.43] | 0.3 [0.24, 0.35] | 0.49 [0.44, 0.54] | 0.44 [0.38, 0.49] | 0.39 [0.33, 0.44] |
| LGK | | 0.19 [0.17, 0.21] | 0.2 [0.17, 0.22] | 0.2 [0.18, 0.22] | 0.21 [0.19, 0.23] |
| PPARG | 0.02 [-0.02, 0.06] | 0.34 [0.30, 0.38] | 0.37 [0.34, 0.41] | 0.25 [0.21, 0.28] | 0.42 [0.38, 0.45] |
| PTEN | 0.03 [-0.00, 0.07] | 0.33 [0.30, 0.36] | 0.33 [0.30, 0.36] | 0.34 [0.31, 0.37] | 0.48 [0.45, 0.51] |
| TPMT | 0.03 [-0.01, 0.07] | 0.42 [0.39, 0.45] | 0.4 [0.37, 0.44] | 0.34 [0.30, 0.37] | 0.46 [0.43, 0.49] |
| haeIIIM | | 0.43 [0.40, 0.47] | 0.62 [0.59, 0.65] | 0.59 [0.55, 0.62] | 0.62 [0.59, 0.65] |
| bgl3 | | 0.57 [0.54, 0.59] | 0.62 [0.59, 0.64] | 0.49 [0.46, 0.52] | 0.65 [0.63, 0.68] |
| GFP | | 0.32 [0.25, 0.38] | 0.49 [0.44, 0.55] | 0.4 [0.34, 0.46] | 0.36 [0.29, 0.43] |
| Ube4b | 0.32 [0.24, 0.39] | 0.29 [0.22, 0.36] | 0.36 [0.29, 0.43] | 0.4 [0.33, 0.45] | 0.48 [0.41, 0.54] |
| BRCA1_2015_Y2H | | 0.31 [0.25, 0.37] | 0.06 [0.00, 0.13] | 0.19 [0.13, 0.24] | 0.22 [0.16, 0.27] |
| BRCA1_2015_E3 | | 0.07 [0.04, 0.10] | -0.02 [-0.05, 0.01] | 0.04 [0.01, 0.07] | 0 [-0.03, 0.04] |
| bla | | 0.57 [0.55, 0.59] | 0.68 [0.66, 0.70] | 0.07 [0.05, 0.09] | 0.68 [0.66, 0.69] |
| IgG1 | | 0.2 [0.11, 0.30] | 0.33 [0.22, 0.42] | 0.34 [0.24, 0.43] | 0.3 [0.20, 0.39] |
| SetCommon | 0.1 [0.08, 0.11] | 0.29 [0.27, 0.30] | 0.33 [0.32, 0.34] | 0.23 [0.22, 0.25] | 0.41 [0.40, 0.42] |

**Table S7: Spearman ρ between five variant effect prediction methods and experimentally measured beneficial effect scores from 22 DMS experiments in SetAll.**

If a method did not perform predictions for a given set, cells are left empty. Values in parentheses denote 95% confidence intervals. nan values are caused by lack of diversity in prediction scores, making calculation of ρ impossible. SetCommon contains 15,200 beneficial effect SAVs from ten proteins for which every method performed predictions (see Tables 1,2 and Figure S4).

| | Envision | Conservation | PolyPhen-2 | SIFT | SNAP2 |
|---|---|---|---|---|---|
| YAP1 | 0.04 [-0.14, 0.23] | -0.05 [-0.22, 0.13] | 0.13 [-0.06, 0.32] | 0.06 [-0.12, 0.21] | 0.08 [-0.12, 0.26] |
| MAPK1 | -0.07 [-0.10, -0.04] | -0.02 [-0.04, 0.01] | -0.1 [-0.13, -0.07] | -0.06 [-0.09, -0.03] | -0.14 [-0.17, -0.11] |
| BRCA1 | -0.06 [-0.15, 0.03] | -0.03 [-0.12, 0.07] | -0.14 [-0.22, -0.05] | 0.01 [-0.08, 0.10] | -0.04 [-0.12, 0.04] |
| CCR5 | -0.05 [-0.09, -0.02] | -0.04 [-0.08, -0.00] | -0.08 [-0.12, -0.05] | -0.06 [-0.10, -0.02] | -0.03 [-0.07, 0.01] |
| CXCR4 | -0.01 [-0.07, 0.04] | 0.02 [-0.03, 0.07] | 0.01 [-0.04, 0.06] | 0 [-0.05, 0.05] | 0.04 [-0.01, 0.09] |
| HSP82_2011 | | 0.46 [-0.37, 0.84] | | [nan, nan] | 0.36 [-0.33, 0.86] |
| HSP82_2013 | | -0.43 [-0.81, 0.10] | | [nan, nan] | 0.21 [-0.54, 0.76] |
| HSP82_2013_Exp | | -0.37 [-0.63, -0.03] | | -0.38 [nan, nan] | -0.13 [-0.46, 0.23] |
| GAL4 | 0.02 [-0.10, 0.13] | 0.09 [-0.04, 0.21] | 0.02 [-0.10, 0.15] | 0.12 [0.01, 0.24] | 0.07 [-0.06, 0.19] |
| LGK | | -0.08 [-0.20, 0.03] | -0.04 [-0.15, 0.07] | -0.07 [-0.19, 0.05] | 0.08 [-0.02, 0.20] |
| PPARG | 0.02 [-0.01, 0.05] | 0.1 [0.07, 0.13] | -0.15 [-0.18, -0.12] | -0.17 [-0.20, -0.14] | -0.03 [-0.06, 0.00] |
| PTEN | 0.07 [0.00, 0.14] | 0.01 [-0.06, 0.08] | 0.01 [-0.06, 0.09] | 0 [-0.07, 0.08] | 0.02 [-0.06, 0.09] |
| TPMT | -0.03 [-0.09, 0.04] | -0.07 [-0.13, -0.00] | -0.07 [-0.14, -0.00] | -0.05 [-0.12, 0.02] | -0.09 [-0.15, -0.02] |
| haeIIIM | | 0.06 [-0.15, 0.25] | 0.06 [-0.14, 0.25] | -0.03 [-0.24, 0.19] | 0.06 [-0.15 0.25] |
| bgl3 | | -0.07 [-0.23, 0.09] | -0.13 [-0.27, 0.02] | -0.06 [-0.22, 0.12] | 0.01 [-0.16, 0.17] |
| GFP | | -0.09 [-0.20, 0.02] | -0.01 [-0.12, 0.10] | 0 [-0.12, 0.11] | 0.08 [-0.02, 0.19] |
| Ube4b | -0.05 [-0.18, 0.07] | 0.03 [-0.08, 0.15] | 0.13 [0.01, 0.25] | 0.2 [0.09, 0.32] | 0.22 [0.09, 0.34] |
| BRCA1_2015_Y2H | | 0.1 [0.02, 0.18] | 0.11 [0.03, 0.19] | 0.09 [0.01, 0.17] | 0.07 [-0.01, 0.14] |
| BRCA1_2015_E3 | | -0.04 [-0.09, 0.01] | -0.03 [-0.09, 0.01] | -0.04 [-0.09, 0.01] | -0.06 [-0.11, -0.00] |
| bla | | 0 [-0.08, 0.09] | 0 [-0.08, 0.08] | -0.02 [-0.09, 0.06] | 0.07 [-0.01, 0.15] |
| IgG1 | | -0.02 [-0.38, 0.32] | -0.14 [-0.46, 0.17] | 0 [-0.32, 0.34] | 0 [-0.33, 0.34] |
| SetCommon | -0.14 [-0.16, -0.13] | -0.08 [-0.09, -0.06] | -0.09 [-0.10, -0.07] | -0.14 [-0.15, -0.12] | 0.02 [0.01, 0.04] |

**Table S8: Mean squared error between five variant effect prediction methods and experimentally measured beneficial effect scores from 22 DMS experiments in SetAll.**

If a method did not perform predictions for a given set, cells are left empty. Values in parentheses denote 95% confidence intervals. SetCommon contains 15,200 beneficial effect SAVs from ten proteins for which every method performed predictions (see Tables 1,2 and Figure S4).

| | Envision | Conservation | PolyPhen-2 | SIFT | SNAP2 |
|---|---|---|---|---|---|
| YAP1 | 0.06 [0.04, 0.09] | 0.15 [0.13, 0.17] | 0.57 [0.51, 0.62] | 0.61 [0.56, 0.66] | 0.2 [0.16, 0.24] |
| MAPK1 | 0.03 [0.03, 0.03] | 0.27 [0.27, 0.27] | 0.48 [0.47, 0.49] | 0.77 [0.76, 0.77] | 0.25 [0.24, 0.25] |
| BRCA1 | 0.04 [0.03, 0.04] | 0.15 [0.14, 0.16] | 0.31 [0.28, 0.34] | 0.63 [0.61, 0.65] | 0.46 [0.44, 0.48] |
| CCR5 | 0.04 [0.04, 0.05] | 0.23 [0.23, 0.24] | 0.4 [0.39, 0.42] | 0.74 [0.73, 0.74] | 0.28 [0.27, 0.29] |
| CXCR4 | 0.02 [0.02, 0.02] | 0.22 [0.21, 0.23] | 0.44 [0.42, 0.46] | 0.68 [0.67, 0.69] | 0.23 [0.22, 0.24] |
| HSP82_2011 | | 0.1 [0.05, 0.15] | | 0.52 [0.36, 0.67] | 0.07 [0.02, 0.13] |
| HSP82_2013 | | 0.11 [0.06, 0.16] | | 0.19 [0.08, 0.32] | 0.1 [0.04, 0.18] |
| HSP82_2013_Exp | | 0.14 [0.10, 0.18] | | 0.48 [0.41, 0.58] | 0.19 [0.13, 0.25] |
| GAL4 | 0.05 [0.04, 0.06] | 0.16 [0.14, 0.17] | 0.38 [0.34, 0.42] | 0.53 [0.49, 0.56] | 0.21 [0.19, 0.24] |
| LGK | | 0.13 [0.12, 0.14] | 0.23 [0.20, 0.26] | 0.32 [0.28, 0.35] | 0.13 [0.11, 0.16] |
| PPARG | 0.08 [0.08, 0.08] | 0.1 [0.09, 0.10] | 0.32 [0.31, 0.33] | 0.46 [0.45, 0.47] | 0.17 [0.16, 0.17] |
| PTEN | 0.05 [0.04, 0.05] | 0.15 [0.14, 0.16] | 0.35 [0.33, 0.37] | 0.52 [0.50, 0.54] | 0.34 [0.32, 0.35] |
| TPMT | 0.03 [0.02, 0.03] | 0.21 [0.20, 0.22] | 0.39 [0.37, 0.41] | 0.59 [0.57, 0.61] | 0.16 [0.15, 0.17] |
| haeIIIM | | 0.13 [0.10, 0.15] | 0.18 [0.13, 0.23] | 0.34 [0.28, 0.40] | 0.09 [0.06, 0.13] |
| bgl3 | | 0.22 [0.19, 0.24] | 0.2 [0.14, 0.25] | 0.59 [0.54, 0.65] | 0.12 [0.09, 0.15] |
| GFP | | 0.18 [0.17, 0.19] | 0.34 [0.30, 0.38] | 0.52 [0.48, 0.55] | 0.29 [0.27, 0.31] |
| Ube4b | 0.03 [0.02, 0.04] | 0.19 [0.18, 0.20] | 0.5 [0.46, 0.55] | 0.65 [0.62, 0.69] | 0.21 [0.19, 0.23] |
| BRCA1_2015_Y2H | | 0.18 [0.17, 0.19] | 0.61 [0.59, 0.63] | 0.69 [0.67, 0.70] | 0.45 [0.44, 0.47] |
| BRCA1_2015_E3 | | 0.26 [0.25, 0.26] | 0.72 [0.70, 0.74] | 0.87 [0.86, 0.88] | 0.52 [0.51, 0.54] |
| bla | | 0.21 [0.20, 0.23] | 0.22 [0.20, 0.25] | 0.8 [0.79, 0.82] | 0.07 [0.06, 0.08] |
| IgG1 | | 0.1 [0.07, 0.14] | 0.3 [0.20, 0.42] | 0.29 [0.20, 0.38] | 0.11 [0.06, 0.16] |
| SetCommon | 0.05 [0.04, 0.05] | 0.19 [0.19, 0.20] | 0.4 [0.40, 0.41] | 0.64 [0.63, 0.64] | 0.23 [0.23, 0.24] |

**Table S9: Agreement between independent experimentally measured beneficial effect scores from ten experiments on four proteins.**

Only measurements on the same protein from different publications are compared (see Table S1). Only SAVs are taken into account. Values for beneficial variants of Hsp82 datasets were not included in the average since the sets contained only three variants each.

| | **Spearman ρ** | | **Mean squared error** | |
|---|---|---|---|---|
| DMS measures | Deleterious | Beneficial | Deleterious | Beneficial |
| BRCA1<br>BRCA1_2015_Y2H | 0.41 | 0.09 | 0.08 | 0.04 |
| BRCA1<br>BRCA1_2015_E3 | 0.21 | 0.13 | 0.08 | 0.03 |
| HSP82_2011<br>HSP82_2013 | 0.87 | 0.5 [1] | 0.09 | 0.29 [1] |
| HSP82_2011<br>HSP82_2013_Exp | 0.68 | -0.8 [1] | 0.06 | 0.08 [1] |
| HSP82_2013<br>HSP82_2013_Exp | 0.88 | 0.5 [1] | 0.04 | 0.32 [1] |
| RPL40A_2013<br>RPL40A_2014 | 0.59 | -0.1 | 0.06 | 0.07 |
| RPL40A_2013<br>RPL40A_2014_REL | 0.6 | -0.05 | 0.05 | 0.07 |
| bla<br>bla_2014 | 0.93 | 0.06 | 0.07 | 0.02 |
| Average | 0.65 | 0.03 | 0.07 | 0.05 |

[1] Not included in the average due to small size of the data set (n = 3)

**Table S10: p-values for the difference between AUCs on SetCommonSyn sets.**

Statistical tests were performed using the roc.test function from the pROC R package with default settings [24, 25]. P-values given as 0 were too small for reporting in R. Generally, the value of p-values for such large datasets should be questioned and more emphasis be put on the confidence intervals themselves as reported in Figs. 1f, S4f, S10, S11. We report them here to show the clear trends and since some people might expect to see them. SetCommonSyn90|95|99 refers to the set of SAVs for which a prediction is available from every method and the respective thresholding scheme to classify SAVs as neutral or effect could be applied (see Methods). Conserv. denotes the naïve conservation-based prediction method.

| | | deleterious only SetCommon | | | beneficial only SetCommon | | |
|---|---|---|---|---|---|---|---|
| Method 1 | Method 2 | Syn90 | Syn95 | Syn99 | Syn90 | Syn95 | Syn99 |
| PolyPhen-2 | SIFT | 0 | 0 | 0 | 0 | 0 | 0.0188 |
| PolyPhen-2 | SNAP2 | 0 | 0 | 0 | 0.0002 | 0.1106 | 0.2243 |
| PolyPhen-2 | Envision | 0 | 0 | 0 | 0 | 0 | 0 |
| PolyPhen-2 | Conserv. | 0.0048 | 0 | 0.0829 | 0 | 0 | 0 |
| SIFT | SNAP2 | 0 | 0 | 0 | 0 | 0 | 0.0003 |
| SIFT | Envision | 0 | 0 | 0 | 0 | 0 | 0.0002 |
| SIFT | Conserv. | 0 | 0 | 0 | 0 | 0 | 0.0002 |
| SNAP2 | Envision | 0 | 0 | 0 | 0 | 0 | 0 |
| SNAP2 | Conserv. | 0 | 0.0001 | 0 | 0 | 0 | 0 |
| Envision | Conserv. | 0 | 0 | 0 | 0.6074 | 0.7989 | 0.3872 |

**Table S11: The source of all DMS measurements used in this study.**

For reference, MaveDB IDs have been added where existing by October 28th, 2019 [26]. However, no data was obtained from MaveDB.

| Dataset | Source | MaveDB ID |
|---|---|---|
| ccdB | https://www.cell.com/cms/10.1016/j.str.2011.11.021/attachment/95e06292-4986-4fdf-b289-334b9372cf07/mmc2.xls | NA |
| YAP1 [1] | https://www.pnas.org/highwire/filestream/610483/field_highwire_adjunct_files/1/sd01.xls | mavedb:00000002-a |
| MAPK1 | https://www.cell.com/cms/10.1016/j.celrep.2016.09.061/attachment/f92aa769-5a2d-43ea-8618-161907ef54b8/mmc2.xlsx | NA |
| BRCA1 | https://static-content.springer.com/esm/art%3A10.1038%2Fs41586-018-0461-z/MediaObjects/41586_2018_461_MOESM3_ESM.xlsx | NA |
| CCR5 | ftp://ftp.ncbi.nlm.nih.gov/geo/series/GSE100nnn/GSE100368/suppl/GSE100368_enrichment_ratios_CCR5.xlsx | NA |
| CXCR4 | ftp://ftp.ncbi.nlm.nih.gov/geo/series/GSE100nnn/GSE100368/suppl/GSE100368_enrichment_ratios_CXCR4.xlsx | NA |
| HSP82_2011 | https://www.pnas.org/highwire/filestream/605883/field_highwire_adjunct_files/2/sd02.csv | mavedb:00000011-a |
| HSP82_2013 | https://onlinelibrary.wiley.com/action/downloadSupplement?doi=10.1111%2Fevo.12207&file=evo12207-sup-0006-dataS1.xls | mavedb:00000040-a |
| HSP82_2013_Exp | https://doi.org/10.1371/journal.pgen.1003600.s014 | mavedb:00000039-a |
| GAL4 [1] | https://media.nature.com/original/nature-assets/nmeth/journal/v12/n3/extref/nmeth.3223-S2.xlsx | mavedb:00000012-a |
| LGK | https://figshare.com/authors/Justin_Klesmith/792792 | NA |
| PPARG | Author contact (Majithia) | NA |
| PTEN | https://static-content.springer.com/esm/art%3A10.1038 | mavedb:00000013-a |

| | | |
|---|---|---|
| | %2Fs41588-018-0122-z/MediaObjects/41588_2018_122_MOESM3_ESM.txt | |
| TPMT | https://static-content.springer.com/esm/art%3A10.1038%2Fs41588-018-0122-z/MediaObjects/41588_2018_122_MOESM4_ESM.txt | mavedb:00000013-b |
| haeIIIM [1,2] | https://doi.org/10.1371/journal.pcbi.1004421.s003 | NA |
| bgl3 | Author contact (Abate) | NA |
| GFP | https://figshare.com/articles/Local_fitness_landscape_of_the_green_fluorescent_protein/3102154 | NA |
| Ube4b [1] | https://www.pnas.org/highwire/filestream/612049/field_highwire_adjunct_files/1/sd01.xlsx | mavedb:00000004-a |
| BRCA1_2015_Y2H [1] | http://www.genetics.org/lookup/suppl/doi:10.1534/genetics.115.175802/-/DC1/genetics.115.175802-6.xls | mavedb:00000003-b |
| BRCA1_2015_E3 [1] | http://www.genetics.org/lookup/suppl/doi:10.1534/genetics.115.175802/-/DC1/genetics.115.175802-6.xls | mavedb:00000003-a |
| bla [1] | https://ars.els-cdn.com/content/image/1-s2.0-S0092867415000781-mmc1.xlsx | NA |
| IgG1 | Author contact (Traxlmayr) | NA |
| RPL40A_2013 | https://www.sciencedirect.com/science/article/pii/S0022283613000636?via%3Dihub (Table S2) | mavedb:00000037-a |
| RPL40A_2014 | https://www.sciencedirect.com/science/article/pii/S0022283614002587?via%3Dihub (Table S2) | mavedb:00000038 |
| RPL40A_2014_REL | https://www.sciencedirect.com/science/article/pii/S0022283614002587?via%3Dihub (Table S2) | mavedb:00000038 |
| bla_2014 | https://academic.oup.com/mbe/article/31/6/1581/2925654#supplementary-data (Table S2) | |

[1] Data exists in pre-parsed format from a previous analysis. Sources are those used in the original parsing.

[2] Data was originally provided by authors when still unpublished and has not been re-parsed with the files now available.

**Table S12: Best matching protein sequences for every DMS measurement used in this study.**

The sequences denoted here were used as input to the prediction methods. Often indices given for the experimental values have to be shifted by a number of positions to fit to the database sequence. SID is short for sequence identity. Substitutions found in the experimental sequences were introduced to the UniProtKB sequences before submitting them for prediction.

| Dataset | Best match | SID of best match with exp. seq. (excluding non-matched regions) | Substitutions within matched region |
|---|---|---|---|
| ccdB | UniProtKB P62554 (CCDB_ECOLI) | 100 | 0 |
| YAP1 | UniProtKB P46937 (YAP1_HUMAN) | 100 | 0 |
| MAPK1 | UniProtKB P28482 (MK01_HUMAN) | 100 | 0 |
| BRCA1 | UniProtKB P38398 (BRCA1_HUMAN) | 100 | 0 |
| CCR5 | UniProtKB P51681 (CCR5_HUMAN) | 100 | 0 |
| CXCR4 | UniProtKB P61073 (CXCR4_HUMAN) | 100 | 0 |
| HSP82_2011 | UniProtKB P02829 (HSP82_YEAST) | 100 | 0 |
| HSP82_2013 | UniProtKB P02829 (HSP82_YEAST) | 100 | 0 |
| HSP82_2013_Exp | UniProtKB P02829 (HSP82_YEAST) | 100 | 0 |
| GAL4 | UniProtKB P04386 (GAL4_YEAST) | 100 | 0 |
| LGK | UniProtKB B3VI55 (B3VI55_LIPST) | 99.317 | 3 |
| PPARG | UniProtKB P37231 (PPARG_HUMAN) | 100 | 0 |
| PTEN | UniProtKB P60484 (PTEN_HUMAN) | 100 | 0 |
| TPMT | UniProtKB P51580 (TPMT_HUMAN) | 100 | 0 |

| | | | |
|---|---|---|---|
| haeIIIM | UniProtKB P20589 (MTH3_HAEAE) | 98.48 | 5 |
| bgl3 [1] | UniProtKB Q59976 (Q59976_STRSQ) | 99.165 | 4 |
| GFP | UniProtKB P42212 (GFP_AEQVI) | 98.723 | 1 |
| Ube4b | UniProtKB Q9ES00 (UBE4B_MOUSE) | 100 | 0 |
| BRCA1_2015_Y2H | UniProtKB P38398 (BRCA1_HUMAN) | 99.7 | 1 |
| BRCA1_2015_E3 | UniProtKB P38398 (BRCA1_HUMAN) | 99.7 | 1 |
| bla | UniProtKB P62593 (BLAT_ECOLX) | 100 | 0 |
| IgG1 | UniProtKB P0DOX5 (IGG1_HUMAN) | 100 | 0 |
| RPL40A_2013 | UniProtKB P0CH08 (RL40A_YEAST) | 100 | 0 |
| RPL40A_2014 | UniProtKB P0CH08 (RL40A_YEAST) | 100 | 0 |
| RPL40A_2014_REL | UniProtKB P0CH08 (RL40A_YEAST) | 100 | 0 |
| bla_2014 | UniProtKB P62593 (BLAT_ECOLX) | 99.3 | 0 |

[1] The entry from UniProtKB is shorter than the sequence implied by the experimental values. Therefore, the sequence implied by the experimental effect measures was used but without the C-terminal His-Tag (GDPNSSSVDKLAAALEHHHHHH).

**Table S13: The functional scores used from every DMS study.**

Column or worksheet references below are indexed at 1 and refer to the files denoted in Table S11.

| Dataset | Functional score |
|---|---|
| ccdB | MS_seq score (column 10) |
| YAP1 | The only given value, column 5 in the pre-converted file from a previous analysis |
| MAPK1 | log2 fold change of early time point vs. DOX (column 36) |
| BRCA1 | Mean function scores across both replicates, averaged over all codons for each amino acid (column 16) |
| CCR5 | Average expression measured by anti-myc FITC and Alexa stains, each with two replicates (columns 4,5,10,11). Only for cases with >100 reads in the Naive Library1. |
| CXCR4 | Average expression measured by anti-myc FITC and Alexa stains, each with two replicates (columns 4,5,8,9). Only for cases with >100 reads in the Naive Library1. |
| HSP82_2011 | EMPIRIC[1] selection coefficient of doubling time between wt yeast and cells with mutant Hsp82 (column 4). Averaged over all codons for each amino acids. |
| HSP82_2013 | EMPIRIC[1] selection coefficient at 30 degrees (column 3) |
| HSP82_2013_Exp | Average growth with mutant Hsp82 under different promotors (column 11). Scores denotes as <0.034 or <0.014 were set to equal the respective value. |
| GAL4 | log2 enrichment of “Selection C, 64 hours”, i.e. highest concentration of the competitive inhibitor and lack of histidine (worksheet 6). |
| LGK | Only data from the first selection (codon-optimized LGK without amino acid substitutions). Normalized Enrich[2] value (column 3). |
| PPARG | The only score in the file (integrated function score) |
| PTEN | VAMP-seq[3] abundance across all replicates (column 7) |
| TPMT | VAMP-seq[3] abundance across all replicates (column 7) |
| haeIIIM | Relative fitness after 17 rounds of selection. Column 9 for non-synonymous, 8 for synonymous in the respective worksheets. |
| bgl3 | log2 enrichment, the only value in the file. |

| | |
|---|---|
| GFP | log fluorescence (column 3) only of mutants with single variants |
| Ube4b | log2 enrichment normalized to wt (column 3) |
| BRCA1_2015_Y2H | Yeast two-hybrid measurements of BRCA-BARD1 binding (column 2) |
| BRCA1_2015_E3 | E3 ligase activity (column 5) |
| bla | Fitness under ampicillin at highest concentration (2500 ug/ml). Averaged over both replicates (columns 7 and 14) |
| IgG1 | Stability landscape, only values with a frequency >= 0.0005 |
| RPL40A_2013 | EMPIRIC[1] selection coefficient (column 3) |
| RPL40A_2014 | log2 between activity and display (column 3) |
| RPL40A_2014_REL | RPL40A_2014 relative to wild-type (column 4) |
| bla_2014 | “Fitness” from Table S2, column 18 |

[1] EMPIRIC: [27]

[2] Enrich: [28]

[3] VAMP-seq: [12]

**Table S14: Values that denote wild type-like behaviour in the raw DMS measures for every dataset.**

| Dataset | Wild-type score |
|---|---|
| ccdB | 2 |
| YAP1 | 1 |
| MAPK1 | 0 |
| BRCA1 | 0 |
| CCR5 | 0 |
| CXCR4 | 0 |
| HSP82_2011 | 0 |
| HSP82_2013 | 0 |
| HSP82_2013_Exp | 1 |
| GAL4 | 0 |
| LGK | 0 |
| PPARG | 0 |
| PTEN | 1 |
| TPMT | 1 |
| haeIIIM | 1 |
| bgl3 | 0 |
| GFP | 3.7 |
| Ube4b | 0 |
| BRCA1_2015_Y2H | 1 |
| BRCA1_2015_E3 | 1 |
| bla | 1 |
| IgG1 | 1 |
| RPL40A_2013 | 0 |
| RPL40A_2014 | 0 |
| RPL40A_2014_REL | 1 |
| bla_2014 | 1 |

#### Table S15: UniProtKB identifiers used for Envision predictions

Listed identifiers were used to retrieve Envision predictions via the webserver (https://envision.gs.washington.edu/shiny/envision_new/) on January 6$^{th}$, 2019. Predictions for other datasets could not be retrieved because there were no perfect sequence matches in UniProtKB (Romero2015_Bgl3, Klesmith2015_LGK, RockahShmuel2015_MTH3, Sarkisyan2016_GFP, Starita2015_BRCA1), those matches were from UniProt entries of organisms unsupported by Envision and no equal sequence from a supported organism could be found (Adkar2012_CcdB, Stiffler2015_TEM1), or unknown reasons (Hietpas2011_HSP90, Hietpas2013_HSP90, Jiang2013_HSP90, Traxlmayr2012_IgG1_CH3).

| Dataset | UniProtKB identifier |
|---|---|
| BRCA1 | P38398 |
| CXCR4 | P61073 |
| CCR5 | P51681 |
| PPARG | P37231 |
| PTEN | P60484 |
| TPMT | P51580 |
| YAP1 | P46937 |
| MAPK1 | P28482 |
| Gal4 | P04386 |
| Ube4b | Q9ES00 |

#### SOM_Note1: ~25% beneficial effect variants in Envision training set.

The Envision training set is available online (https://github.com/FowlerLab/Envision2017/tree/master/data). The simple R script below reads this file and confirms the overall number of 28,545 variants from the manuscript [29]. After excluding the three DMS measurements that were not used for training, 20,676 variants remain. Of those 5,135 have a scaled effect score larger than 1 which in the normalization scheme employed by Gray et al. denotes beneficial effect.

```
training <- read.csv("dmsTraining_2017-02-20.csv",header = TRUE)

training1 <-training[which( training$mut_type =='missense'),]

training_final <- training1[!(training1$dms_id=='Brca1_E3' |

                                training1$dms_id=='Brca1_Y2H' |

                                training1$dms_id=='Ubiquitin'),]

length(training_final$scaled_effect1)

[1] 20676

sum(training_final$scaled_effect1 > 1.0)

[1] 5135
```

**SOM_Note2: Selection of appropriate performance measures for regression analyses.**

Possible performance measures for regression data could be broadly categorized into correlation measures and error measures. Correlation measures, such as Pearson correlation coefficient R and Spearman rank correlation coefficient ρ evaluate the relationship between, in our case, experimental and predicted variant effect scores. R checks for the existence of a linear relationship but is highly susceptible to outliers and not considered a robust measure [30]. R further assumes that the data is normally distributed which is not the case (Shapiro-Wilk, p <= 1e-7 for all datasets; see also Figs. S3, S5 gray distributions). We report R values regardless to maintain comparability to earlier analyses but caution its use to infer prediction performance.

ρ on the other hand is robust to outliers and measures only whether there is a monotonic relationship between the DMS data and predicted effect. That is, a higher experimental effect should yield a higher predicted effect, however the increase does not need to be linear.

Both measures are invariant to scale and shifts in the data, therefore transforming experimental and predicted effect values to lie between 0 and 1 does not change R or ρ compared to their calculation on the raw values.

The above measures only check for a positive correlation between experimental and predicted values, however they do not evaluate the size of the difference between values, i.e. a method that always predicts high effect variants more like in a classification task can score as well as a method that predicts along the whole possible range of values, more like in a regression task. To account for this, we additionally employed a common error measure, the mean squared error (MSE).

Other related measures exist. For example, given the outliers in our DMS data one could argue to use the mean absolute error (MAE) instead of the MSE as it does not disproportionally weigh outliers. That is, severely over- or underpredicting the effect of some datapoints is not punished when most variants' effect is predicted correctly. A measure even more robust to outliers would be the median absolute error (MedAE). Both are defined below with all variables named as in SOM_Note2 ($x_i$ experimentally measured effect, $y_i$ predicted).

$$\text{Mean absolute error (MAE)} = \frac{1}{n}\sum_{i=1}^{n}|y_i - x_i|$$

$$\text{Median absolute error (MedAE)} = median(|y_1 - x_1|, \dots, |y_n - x_n|)$$

However, we found no major differences in the performance of methods using MAE or MedAE. In particular, the order of methods on deleterious effect SAVs of SetCommon is unchanged from Envision achieving the lowest error followed by Conservation, SNAP2, Polyphen-2 and then SIFT (see below). The same goes for beneficial effect SAVs except SNAP2 being slightly better than Conservation on MedAE. Given these findings we decided to report only the MSE which is likely the best known.

| | SetCommon deleterious SAVs (n = 17781) | | |
|---|---|---|---|
| | MSE | MAE | MedAE |
| PolyPhen-2 | 0.45 [0.45, 0.46] | 0.6 [0.60, 0.61] | 0.68 [0.67, 0.68] |
| SIFT | 0.58 [0.57, 0.58] | 0.73 [0.72, 0.73] | 0.78 [0.78, 0.79] |
| SNAP2 | 0.3 [0.30, 0.30] | 0.5 [0.49, 0.50] | 0.52 [0.51, 0.52] |
| Envision | 0.06 [0.06, 0.07] | 0.19 [0.18, 0.19] | 0.13 [0.13, 0.14] |
| Conservation | 0.19 [0.19, 0.19] | 0.4 [0.40, 0.40] | 0.42 [0.42, 0.43] |
| | SetCommon beneficial SAVs (n = 15200) | | |
| | MSE | MAE | MedAE |
| PolyPhen-2 | 0.4 [0.40, 0.41] | 0.54 [0.53, 0.54] | 0.59 [0.58, 0.61] |
| SIFT | 0.64 [0.63, 0.64] | 0.77 [0.77, 0.77] | 0.83 [0.83, 0.84] |
| SNAP2 | 0.23 [0.23, 0.24] | 0.41 [0.41, 0.41] | 0.4 [0.39, 0.41] |
| Envision | 0.05 [0.04, 0.05] | 0.16 [0.16, 0.16] | 0.12 [0.12, 0.13] |
| Conservation | 0.19 [0.19, 0.20] | 0.4 [0.40, 0.41] | 0.42 [0.42, 0.42] |

For R and ρ, baseline performance is intuitive with no correlation being observed at a value of 0. For MSE, the typical baseline performance is the MSE of a method that always predicts the mean value of the observed (here, experimental DMS) distribution. This concept can be combined in a single score referred to as R2 (https://scikit-learn.org/stable/modules/model_evaluation.html#r2-score). However, calculating R2 we found that values were in almost all cases <0, i.e. prediction methods performed worse than always predicting the mean. However, this is a somewhat unfair comparison: (a) Since the distributions of experimentally determined effect are often highly skewed, hence knowing the true mean is already a large advantage, while prediction methods need to account for a much wider possible range of values. (b) The smaller the spread in values of the experimental effect data (as measured for example by the sample standard deviation), the larger the effect in (a) weighs. We found that R2 tended to be particularly low for experimental datasets that had small standard deviations. As a more realistic baseline, the performance of predictions based solely on PSI-BLAST PSSMs are provided (see Methods).

#### SOM_Note3: Processing applied to experimental effect scores.

First scores were shifted such that the wild-type effect score (Table S14) is at 0. If necessary for the given measurement, scores were then mirrored at 0 such that deleterious effect SAVs have scores with values >0 and beneficial effect scores' values are <0. Next, scores of beneficial effect SAVs were mirrored at 0, such that all values larger than 0 denote effect – either beneficial or deleterious. Finally, scores were interpolated, separately for each of the 22 DMS measurements, to lie between 0 (wild-type) and 1 (highest effect). This interpolation does not affect Spearman ρ or the mean squared error within each dataset.

#### SOM_Note4: Employed performance measures for regression analyses

$$\text{Pearson R (R)} = \frac{n\sum_{i=1}^{n} x_i y_i - \sum_{i=1}^{n} x_i \sum_{i=1}^{n} y_i}{\sqrt{n\sum_{i=1}^{n} {x_i}^2 - (\sum_{i=1}^{n} x_i)^2}\sqrt{n\sum_{i=1}^{n} {y_i}^2 - (\sum_{i=1}^{n} y_i)^2}}$$

$$\text{Spearman ρ (ρ)} = \frac{n\sum_{i=1}^{n} rx_i ry_i - \sum_{i=1}^{n} rx_i \sum_{i=1}^{n} ry_i}{\sqrt{n\sum_{i=1}^{n} {rx_i}^2 - (\sum_{i=1}^{n} rx_i)^2}\sqrt{n\sum_{i=1}^{n} {ry_i}^2 - (\sum_{i=1}^{n} ry_i)^2}}$$

$$\text{Mean squared error (MSE)} = \frac{1}{n}\sum_{i=1}^{n}(y_i - x_i)^2$$

Here, $n$ denotes the number of SAVs, $x_i$ the experimentally measured effect score for SAV $i$ and $y_i$ the respective predicted effect score. For ρ both experimental measurements and predictions are ranked. Then, $rx_i$ denotes the rank of the i-th measurement, $ry_i$ the rank of the i-th prediction.

R and ρ were calculated using the SciPy stats module [31], MSE with the scikit-learn metrics module [32].

**SOM_Note5: Different score scaling schemes for Envision.**

Envision predicts scores between 0 (most severe deleterious effect) to 1 (wild-type like). However, scores >1 are also predicted and given Envision's training data should correspond to beneficial effect [29]. Several options exist to adjust these predictions scores to the scheme used in this work, i.e. values at 0 are wild-type like and everything larger is increasingly severe effect. For all schemes below, it is assumed that the maximum value predicted by Envision is 1.2 (MAX_ENV_SCORE). All predictions performed as part of our analyses yielded scores below this theoretical maximum.

$$Envision_{Lin} = MAX\_ENV\_SCORE - raw\_score$$

$$Envision_{DB} = Envision = \begin{cases} 1 - raw\_score, & raw\_score \leq 1 \\ raw\_score - 1, & raw\_score > 1 \end{cases}$$

$$Envision_{D} = \begin{cases} 1 - raw\_score, & raw\_score \leq 1 \\ NaN, & raw\_score > 1 \end{cases}$$

$$Envision_{DBS} = \begin{cases} 1 - raw\_score, & raw\_score \leq 1 \\ (raw\_score - 1) * 1/(MAX\_ENV\_SCORE - 1), & raw\_score > 1 \end{cases}$$

Envision_Lin: Ignoring the fact, that values larger than 1 denote beneficial effect, simply invert the score. This effectively declares variants predicted to have beneficial effect as the least effect variants (values in [0, 0.2], while variants predicted to have wild-type like effect as slightly higher effect variants (=0.2), followed by variants predicted to have deleterious effect (]0.2, 1])

Envision_DB: Deleterious and beneficial effect predictions are treated separately. After processing, scores within [0, 0.2] are variants originally predicted to have low deleterious effect or to have beneficial effect. All scores in ]0.2,1] were predicted to have deleterious effect. However, the minimal raw score ever predicted in our set is 0.39, therefore the maximum score seen for Envision_DB is just 0.61.

Envision_D: Same as Envision_DB but ignoring all predictions of beneficial effect, hence decreasing the number of samples over which correlation measures can be calculated.

Envision_DBS: Same as Envision_DB, however beneficial effect variants are scaled to be within [0,1], same as the deleterious predictions. This implies that the strongest beneficial effect is comparable to the strongest deleterious effect. Something that the experimental assays used in DMS studies cannot guarantee and which is why analyses in our manuscript treat deleterious and beneficial variants separately.

Envision_D was not found to perform better, even on just deleterious effect SAVs in initial exploratory analyses and would additionally further reduce the largest common subset of SAVs for all dataset. Therefore, Envision_D was disregarded. Among the other schemes, Envision_DB generally showed the same or better performance than the other two sets and was hence chosen as the final approach. It is simply referred to as Envision throughout the manuscript.
